# Design and characterization of peptide-responsive biosensors

**DOI:** 10.64898/2026.10.03.755986

**Authors:** Dylan M. Brown, Alia Clark-ElSayed, Taylor Michelle Mott, Isa Madrigal Harrison, Brice Graham Hendrickson, Julius B. Lucks, Andrew D. Ellington

## Abstract

Gram-positive bacteria use secreted peptides and peptide-responsive transcription factors for quorum sensing. Here, we demonstrate that intracellular expression of these peptide effectors with their cognate transcription factors leads to regulation of gene expression in both *in vitro* and *in vivo* systems. Because both effector and regulatory parts can be readily expressed, complex circuits, including auto-activation schemes, can be developed, with switch-like behaviors. Moreover, the peptide effectors can be fused to proteins, enabling a direct intracellular readout of translation and potentially allowing any protein to encode regulatory interactions in cells. Our results support a foundation for regulatory circuits that extend beyond canonical components such as LacI and TetR.

## Introduction

Bacteria communicate through quorum sensing to synchronize group behaviors^1^. Quorum sensing is frequently mediated by the production and detection of diffusible signaling molecules that accumulate as cell density increases, resulting in coordinated changes in gene expression that regulate collective behaviors such as biofilm formation and the secretion of virulence factors^2–4^. Gram-positive bacteria predominantly use small, ribosomally-produced pheromone peptides^5^ for quorum sensing, which are typically produced as propeptides containing an N-terminal secretion signal to direct their export^6^. The processed, secreted peptides are then detected at the cell surface via either a signal-transduction receptor or a specific oligonucleotide permease, which transports them to the cytoplasm where they interact intracellularly with RRNPP (<u>R</u>ap, <u>R</u>gg, <u>N</u>prR, <u>P</u>lcR, and <u>P</u>rgX) receptors.

The structure and mechanism of the RRNPP family of proteins have been extensively characterized^6–11^. Transcription factors (TFs) in this family have an N-terminal DNA-binding domain with a helix-turn-helix motif. The C-terminal pheromone-peptide-binding domain interacts with the peptide through the tetratricopeptide repeat motifs which mediate oligomerization of the TF^12^. The signaling peptides for these TFs are linear, predominantly hydrophobic, and typically range in length from five to ten amino acids^5^ (see also **Table 1**).

**Table 1.**
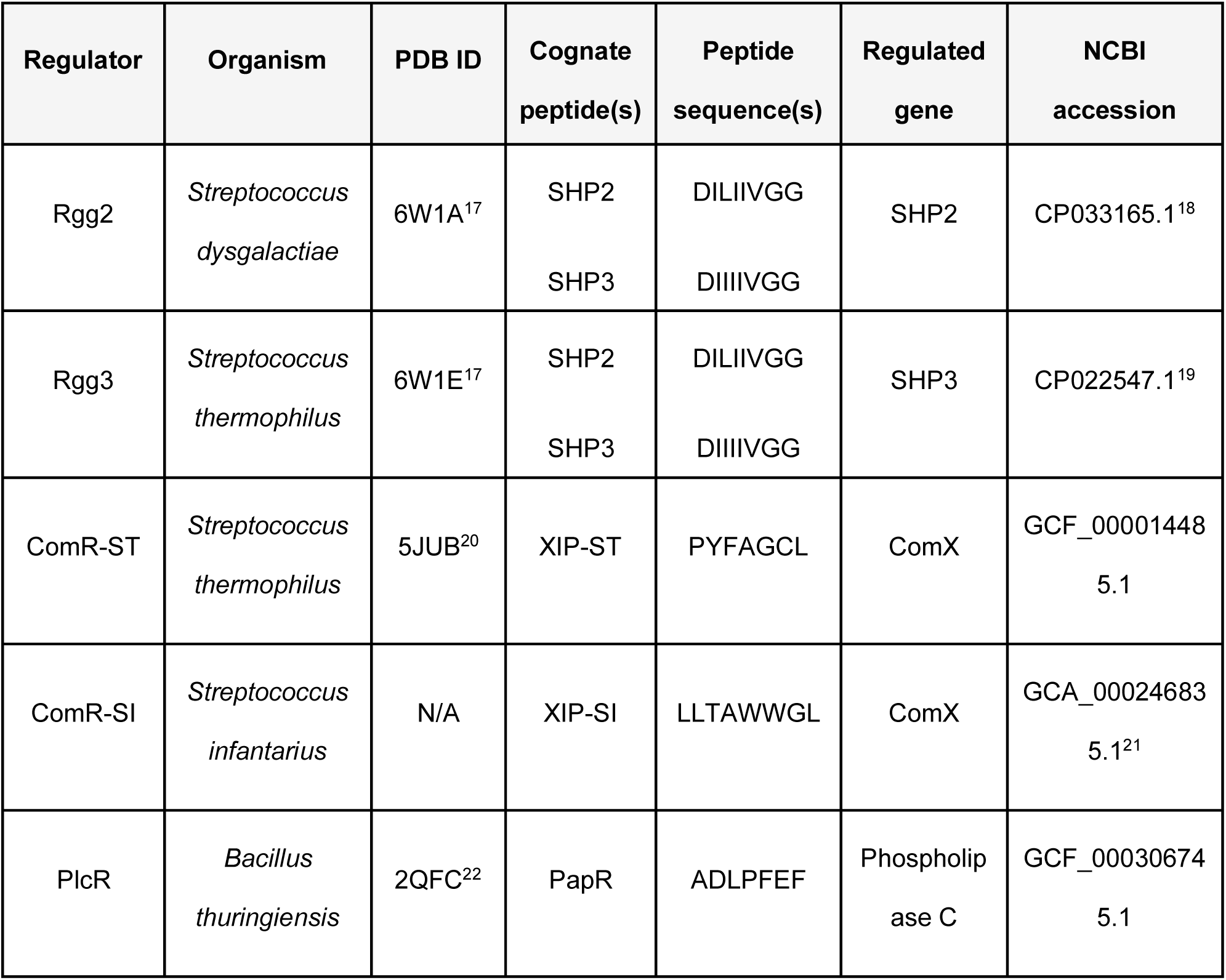
Peptide-responsive transcriptional regulators and their cognate signaling peptide. Listed are the regulator protein, source organism, available Protein Data Bank (PDB) structure, cognate peptide(s), regulated gene, and NCBI accession number.

Synthetic biology approaches are often predicated on the notion that biology can be modularly programmed. However, many parts that are routinely used in synthetic biology have idiosyncratic functions, especially in regulation. While common TFs such as LacI and TetR can be used with external inducers, their ability to participate in complex circuitry is limited by the fact that the production of such inducers often requires numerous biochemical transformations that may not be orthogonal to cellular metabolism. To the extent that peptide-sensing TFs could be adapted to sense intracellularly- or cell-free-produced peptides, this would be a boon for the development of more complex feedback circuits, and for potentially further diversifying what is already an orthogonal regulatory part.

Herein, we attempt to adapt RRNPP family members to function as fully intracellular, peptide-responsive TFs. We find that a number of family members can be adapted to regulate both *in vitro* and *in vivo* genetic circuits, driving GFP expression from their native promoters in response to their cognate peptide pheromone, including via auto-feedback loops. The TF Rgg3 proved to be particularly tractable to further engineering, and its cognate SHP3 peptide activator could be used to create fusion proteins that effectively induced gene expression. This finding opens the way to the intracellular monitoring of protein expression via very small fusion constructs, and to the development of new intracellular signaling systems based on proteins not only carrying out their normal functions, but also encoding new regulatory interactions via peptide fusions.

## Results

### Characterizing the RRNPP-family of transcriptional activators in cell-free systems

In order to quickly determine whether RRNPP-family TFs could be engineered, they were assayed in *Escherichia coli* (*E. coli*) lysate (**Fig. 1**). Rgg2 from *Streptococcus dysgalactiae*, Rgg3 and ComR from *Streptococcus thermophilus* (ComR-ST), and PlcR from *Bacillus thuringiensis* were chosen based on the availability of comprehensive structural data. ComR from *Streptococcus infantarius* (ComR-SI) was chosen based on its reported ability to promiscuously bind peptide pheromones^9^ (**Table 1**).

**Fig. 1.**
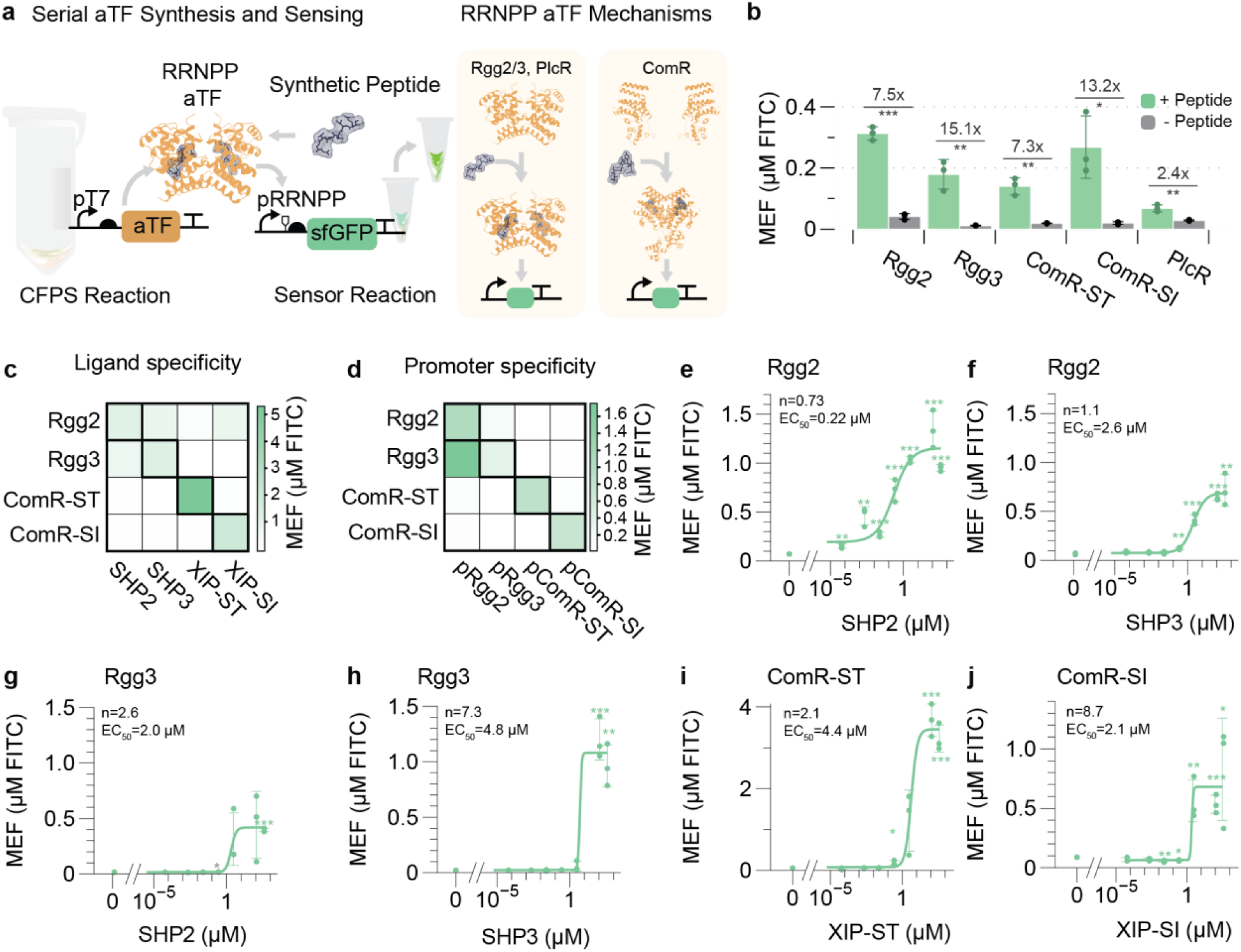
RRNPP-family TFs function in cell-free biosensor reactions. **(a)** Schematic of RRNPP-family TF cell-free biosensor reactions. **(b)** Preliminary functional test of RRNPP biosensors using 1 mg/mL cognate peptide, 9% (v/v) TF CFPS, and 30 nM reporter DNA. **(c)** Ligand specificity heatmap of Rgg2, Rgg3, ComR-ST, and ComR-SI. **(d)** Promoter specificity heatmap of Rgg2, Rgg3, ComR-ST, and ComR-SI. **(e-j)** Dose-response curves for RRNPP TFs and their cognate peptides. **(c-d)** Reactions were carried out using 375 µM peptide, 50 nM reporter DNA, and 9% (v/v) TF CFPS. **(e-j)** Reactions were carried out using 50 nM reporter DNA, 9% (v/v) TF CFPS, and 0, 5.98x10^-5^, 0.0022, 0.025, 0.27, 3.0, 112.5, and 375 µM synthetic peptide. All data were carried out with *n*=3 experimental replicates. Student’s two tailed T-Tests were carried out between the no ligand and induced conditions to determine significance with significant cutoffs of ***p<0.001,**p<0.01, *p<0.05.

For prototyping, a cell-free circuit was designed that should be responsive to the TFs (**Fig. 1a**). The various TFs were synthesized in cell-free protein synthesis (CFPS) reactions overnight and the spent reaction was added to fresh CFPS reactions containing a GFP reporter construct regulated by the identified operator for the RRNPP member. Promoter sequences were identified (**Supplementary Fig. 1**) by first using BLASTP (Protein-Protein Basic Local Alignment Search Tool)^13^ to search for genomes (IDs in **Table S1**) containing TF protein sequences (sequences in **Table S1**), and then operator sequences were extracted based on previously reported DNA-binding targets of each TF (see **Table S1** for references). Reported transcription start sites were obtained from the literature (see **Table S1** for references) and promoter constructs (**Table S1**) were designed to include the region spanning the TF binding site and the transcription start site.

Addition of synthetic peptide was found to activate transcription of the synthetic circuit, and through optimization we found that 0.3 mg/mL of the cognate peptide yielded robust TF activation (**Supplementary Fig. 2**). Activation of GFP expression was observed with almost all of the tested TFs, yielding fold-changes in expression of 7.5x for Rgg2, 15.1x for Rgg3, 7.3x for ComR-ST, 13.2x for ComR-SI, and 2.4x for PlcR (**Fig. 1b**). Due to inconsistent *in vitro* performance, the PlcR sensor was excluded from further experiments. Rgg2 and Rgg3 also functioned in a one-pot reaction system where the TF was co-produced via *in vitro* transcription and translation (**Supplementary Figs. 3** and **4**).

We then tested the orthogonality of the biosensors. Based on previous literature, SHP2 and SHP3, which differ at only one position, are expected to activate both Rgg2 and Rgg3 ^14^. We in fact find that Rgg3 is responsive to SHP2 and SHP3, while ComR-ST and ComR-SI are only responsive to their cognate peptides (**Fig. 1c**). Interestingly, Rgg2 is not only responsive to SHP2 and SHP3, but may respond to ComR-SI’s cognate peptide, XIP-SI (but not XIP-ST, which does not share sequence similarity to XIP-SI) *in vitro*. Cross-reactivity of the promoter sequences was also tested, and it was found that Rgg2 and Rgg3 functioned with each other’s promoter sequences, but not with other RRNPP promoters, while ComR-ST and ComR-SI only function with their cognate promoters (**Fig. 1d)**.

Sensor properties were further characterized by determining dose-responses to synthetic cognate ligands (**Fig. 1e-j**). To assess cooperativity, dose-response measurements were fit to the Hill equation. Dose-response measurements with Rgg2 and Rgg3 were carried out with both SHP2 and SHP3 peptides. Rgg2 was found to be highly sensitive to its cognate peptide SHP2, with a half-maximal effective concentration (EC_50_) value of 0.22 µM and a Hill coefficient (n) of 0.73 (**Fig. 1e**). When Rgg2 is induced with SHP3, we see a shift to higher EC_50_ of 2.6 µM, but a Hill coefficient of 1.1 (**Fig. 1f**). Rgg3 is more sensitive to SHP2 than SHP3 (EC_50_=2.0 µM and 4.8 µM, respectively), but demonstrates stronger positive cooperativity with SHP3 than SHP2 (n=7.3 and 2.6, respectively) (**Fig. 1g-h**). Hill coefficients > 1 were expected, since Rgg2 and Rgg3 predominantly exist as homodimers, with conformational changes being modulated by peptide interactions^6^. ComR-ST similarly showed an EC_50_=4.4 µM and n=2.1 (**Fig. 1i**), while ComR-SI had an extraordinarily high Hill coefficient of n=8.7 (**Fig. 1j**). ComR mostly exists as a monomeric protein in the apo state, with its DNA-binding helices mispositioned and unable to bind DNA effectively until peptide binding triggers a strongly cooperative conformational change and “switch-like” behavior^8^.

The fitted Hill coefficients were substantially higher than the coefficients typically reported for canonical bacterial repressors such as LacI and TetR which are often measured in the range of ∼2^15,16^. The highly cooperative interactions observed likely reflect system-level ultrasensitivity that arises from promoter architecture and TF oligomerization, in addition to cooperativity of peptide binding.

### Genetic circuits with co-translated peptides

We initially used synthetic peptides to characterize RRNPP biosensor properties; however, we next wanted to test whether peptide ligands translated in biological systems were also functional. Using the well-behaved Rgg2 and Rgg3 TFs, a circuit was devised in which reporter expression was impacted by the expression of methionine-modified peptide effectors SHP2 and SHP3, based on both constitutive (**Fig. 2a**) and self-inducing expression of the peptide effector (**Fig. 2d**). For constitutive expression, signal is generated in a template-dependent manner for both Rgg2 (**Fig. 2b**) and Rgg3 (**Fig. 2c**). However, self-induction results in more switch-like behavior (**Fig. 2e-f**) with corresponding increases in Hill coefficients (from n=1.9 to 26.2 for Rgg2, and from 3.1 to 23.8 for Rgg3). ComR-ST and ComR-SI peptides were also found to function when expressed in CFPS reactions containing their cognate TFs (**Supplementary Fig. 5**).

**Fig. 2.**
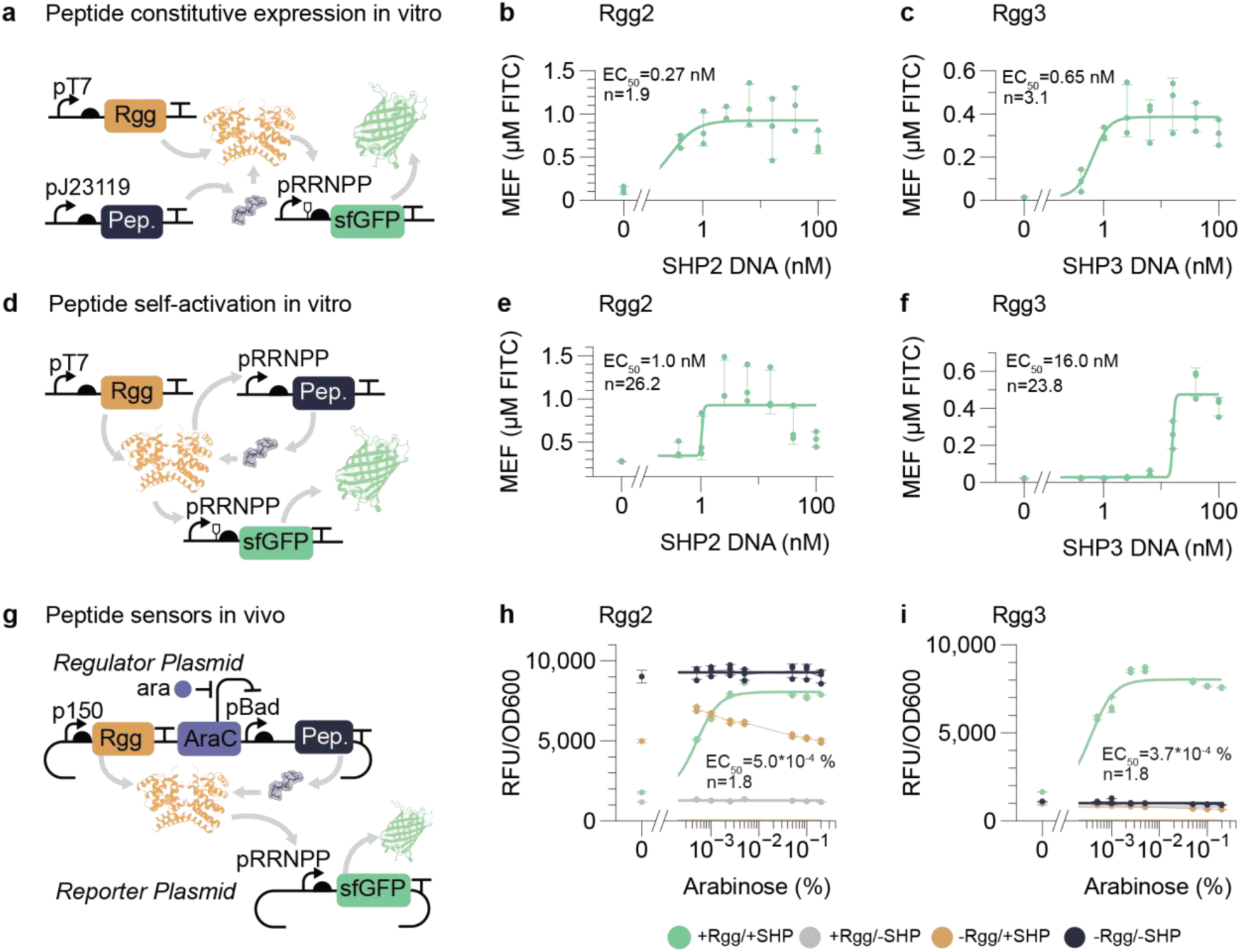
RRNPP-family peptides produced in cell-free and *in vivo* systems activate biosensors. **(a-f)** Cell-free expression assays using peptide-encoding DNA templates. **(a)** Schematic of constitutive *in vitro* peptide expression. **(b, c)** Rgg2 and Rgg3 responses to increasing amounts of pJ23119-SHP2 and pJ23119-SHP3 template DNA, respectively. **(d)** Schematic of *in vitro* peptide self-activation using DNA templates in which an RRNPP-responsive promoter drives peptide expression. **(e, f)** Rgg2 and Rgg3 responses to increasing amounts of pRgg2-SHP2 and pRgg3-SHP3 template DNA, respectively. **(g-i)** *In vivo* assays of RRNPP-type TFs and peptide expression. **(g)** Schematic of the *in vivo* assay. The pBAD-driven peptide activates the transcription factor, changing the sfGFP expression from the reporter plasmid (green). Controls include TF-only (grey), peptide-only (gold), and reporter-only (black). **(h, i)** Rgg2 and Rgg3 responses to increasing arabinose concentrations. Panels b,c,e, and f show raw fluorescence standardized to MEF (µM FITC). Panels h and i show fluorescence in raw fluorescence units (RFU) normalized to cell density (OD) as RFU/OD. All data are carried out with *n*=3 biological replicates for *in vivo* data and experimental replicates for *in vitro* data with error represented as standard deviation (SD).

The ability to quickly establish and tune circuit architectures *in vitro* allows us to design similar architectures for cells. We initially created reporter circuits in which the methionine-augmented peptides were placed under the control of the AraC promoter^23^ (**Fig. 2g**). For both Rgg2 and Rgg3, arabinose-mediated peptide induction yields a dose-dependent increase in signal only when both the transcription factor and peptide are present (**Fig. 2h-i**). No significant response is observed for Rgg3 when the transcription factor, peptide, or reporter is expressed alone (**Fig. 2i**). For Rgg2, the circuit is constitutively on in the absence of the transcription factor, and fully repressed in the presence of TF but in the absence of the peptide (**Fig. 2h**). However, a modest decrease in fluorescence was observed when the SHP2 peptide was expressed in the absence of Rgg2, which may suggest peptide-associated toxicity. Consonant with *in vitro* data, there was little cross-activation in cells by non-cognate peptides, such as XIP (**Supplementary Fig. 6**). Under these circuit conditions, ComR-ST and ComR-SI sensors do not function as anticipated in *E. coli* cells (**Supplementary Fig. 7**).

As in the cell-free system, the pRgg3 promoter in cells was found to be inactive in the absence of its TF; Rgg3 and its effector SHP3 act as a peptide-sensing transcriptional activator (**Fig. 2i**). While *in vitro* the pRgg2 promoter is activated by Rgg2, in cells, however, pRgg2 exhibits basal activity even in the absence of its cognate TF and in the presence of Rgg2 is inhibited, suggesting that Rgg2 acts as a repressor (**Fig. 2h**). These differences between *in vitro* systems and cells highlight the greater complexity of promoter architectures and regulatory interactions in cells, as has been seen with other prokaryotic TFs (e.g., AraC, OmpR)^24,25^. To determine whether promoter architecture was the primary determinant of regulatory function, pRgg2 was assayed with Rgg3, and pRgg3 was similarly assayed with Rgg2; in each case, the promoter configuration (not the TF) determined whether de-repression (pRgg2) or activation (pRgg3) was observed (**Supplementary Fig. 8**).

### Activating gene expression with peptide tags

In order to determine whether the short peptide tag could be fused to other proteins or peptides, essentially turning them into activators or repressors of transcription, we initially fused SHP3 to either terminus of mScarlet. These fusions were expressed alongside a circuit in which Rgg3 controlled activation of GFP expression. In fact, mScarlet fusions at either terminus proved capable of activating GFP expression, both *in vitro* (**Fig. 3a-c**) and *in vivo* (**Fig. 3d-f**).

**Fig. 3.**
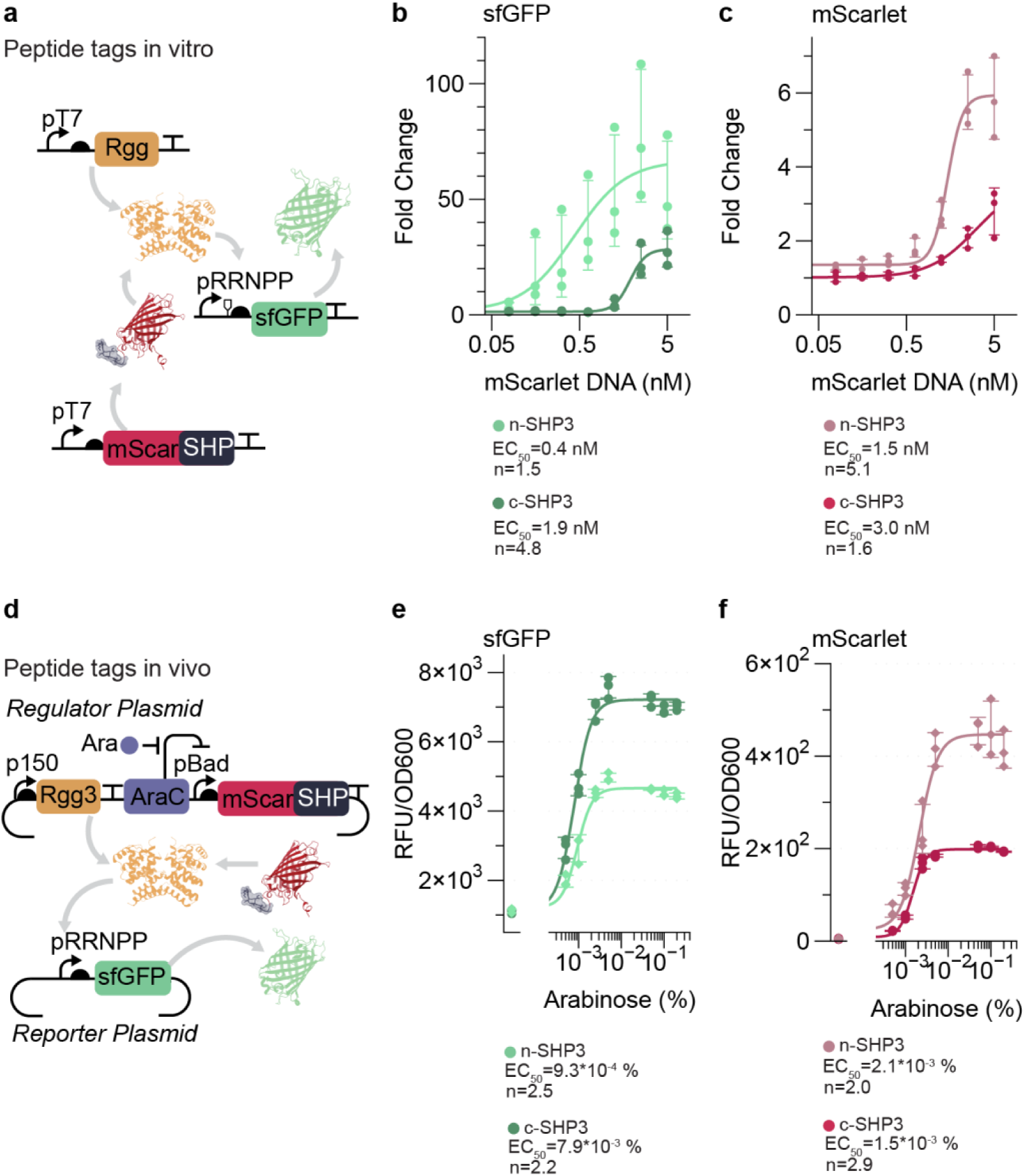
SHP-tagged proteins activate gene expression and alter sensor properties in a terminus-dependent manner. **(a)** Schematic of an *in vitro* expression system for TF activation by a peptide-fusion protein. **(b, c)** Fold change in GFP and RFP signal as a function of mScarlet-peptide fusion DNA, respectively, *in vitro*. **(d)** Schematic of an *in vivo* expression system for TF activation by a peptide-fusion protein. **(e, f)** Normalized GFP and RFP output from Rgg-regulated GFP expression with peptide-tagged mScarlet, respectively, *in vivo*. All data are carried out with *n*=3 biological replicates for *in vivo* data and experimental replicates for *in vitro* data with error represented as SD.

**Fig. 4.**
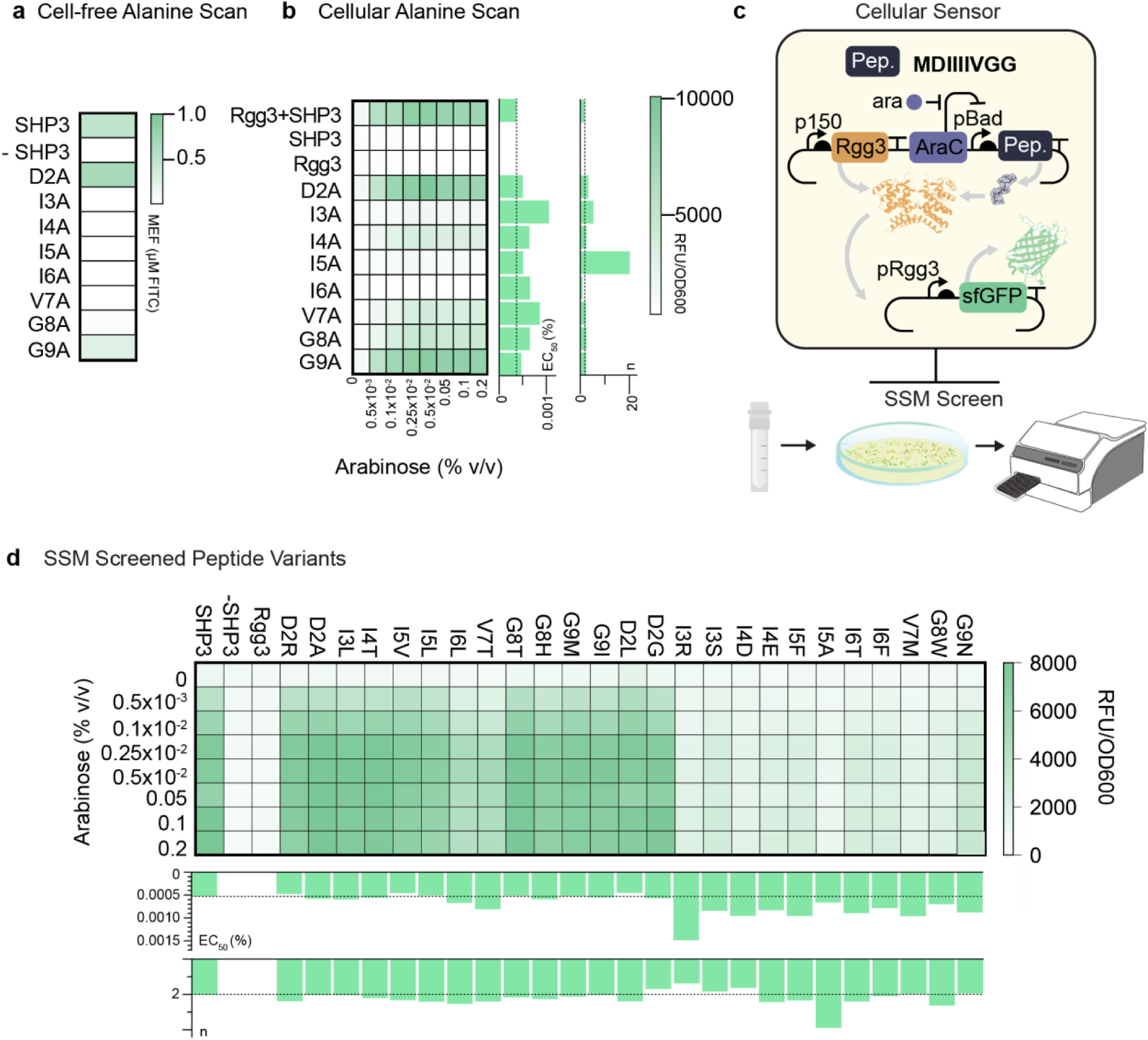
Alanine scanning mutagenesis of SHP3 identifies hydrophobic peptide residues as important for TF activation, and SSM identifies peptide variants with near wild-type activation. **(a)** Cell-free translated alanine scan library. Heat map showing the activity of alanine-scan library variants at a template DNA concentration of 50 nM, peptide DNA concentration of 80 nM, and 9% (v/v) Rgg3 CFPS**. (b)** Cellular-translated alanine scan library. **(c)** Schematic of SSM library screen. The libraries were transformed into cells containing the pRgg3-sfGFP reporter plasmid and plated onto solid media containing 0.005% arabinose. The fluorescent colonies were assayed for fold change in fluorescence in the presence of arabinose. **(d)** Dose-response measurements of identified library variants from the high-activity peptide screen and the medium-activity peptide screen. All data are carried out with *n*=3 biological replicates for *in vivo* data and experimental replicates for *in vitro* data with error represented as SD.

### Determining Rgg3 ligand specificity via SHP3 alanine scan and site-saturation mutagenesis

SHP3 contains an N-terminal acidic residue followed by seven hydrophobic amino acids. Structural studies of the Rgg3-SHP3 complex have revealed extensive interactions that explain how such a small peptide achieves high affinity and specificity for its TF^6^: the N-terminal aspartate side chain and the C-terminus of SHP3 form salt bridges with arginines on Rgg3, while the remaining residues mediate extensive hydrophobic contacts.

To determine how individual residues contributed to signaling function, we carried out an alanine scan of SHP3 both *in vitro* and *in vivo*. The results of the two scans were consistent with one another (**Fig. 4a-b** and **Supplementary Figs. 10-11**). Consistent with the importance of the hydrophobic residues, alanine substitution at any of the four isoleucines (I3, I4, I5, and I6) generally disrupts TF activation. Mutations at residues involved at the N- and C-termini are more readily tolerated, and surprisingly, a D2A mutation increased TF activation relative to the wild-type SHP3 sequence, even though the aspartate side chain is involved in a salt bridge.

The alanine scan of SHP3 indicates that the peptide is amenable to some sequence substitutions. We therefore sought to identify new peptide variants that might also activate Rgg3. Single-position, site-saturation mutagenesis (SSM) libraries at each peptide position were individually cloned and transformed into *E. coli* electrocompetent cells containing the reporter plasmid (**Fig. 4c**). Functional peptide variants were identified by plating libraries onto solid media with and without arabinose, and then picking colonies that fluoresced under blue-light illumination. Colonies that initially showed either high fluorescence (**Supplementary Table 2**) or moderate fluorescence (**Supplementary Table 3**) were subsequently regrown, and dose-response curves were generated as a function of peptide effector induction.

For highly fluorescent variants, substitutions that demonstrated increased activation were identified at every position of the peptide (**Fig. 4d**). Consistent with alanine-scanning mutagenesis, the hydrophobic interactions contributed by the residues at positions 3-6 proved to be most important for TF activation. Substitutions recovered at these positions were largely conservative (for example, I3L, I4T, I5V, I5L, and I6L) and retained activation levels similar to those of the wild-type peptide. By contrast, less responsive variants contained fewer conservative substitutions, extending beyond branched-chain residues to include aromatic and charged amino acids (for example, I5F, I3R, I4D, and I4E). The N-terminus was also found to tolerate less conservative substitutions while maintaining near-wild-type activity (for example, D2R and D2A). In contrast, the C-terminus showed greater sequence diversity, including multiple nonconservative substitutions at glycine residues 8 and 9 (for example, G8T, G8H, G9M, and G9I).

### Peptide variants are functional in vitro and in vivo applications

To determine whether individual peptide variants might impact both *in vitro* and *in vivo* applications, we down-selected a small number of variants for further assay. First, seven peptides from the SSM screen that displayed variable activities *in vivo* were further assayed *in vitro* as self-activating circuits under the control of either Rgg2 or Rgg3. All seven peptides minimally activated or were unable to activate Rgg2 (**Supplementary Fig. 12**). In contrast, all retained activity with Rgg3, with diverse EC_50_ values, Hill coefficients, and dynamic ranges (**Supplementary Fig. 12**). Three variants (I4T, I5V, and V7T) were found to yield either larger (V7T, n=35.1) or smaller (I4T, n=1.8; I5V, n=4.4) Hill coefficients with Rgg3 than the wild-type tag (n= 23.8; **Fig. 5a**). Second, three different variants (G8T, V7T, and G9N) were assayed for their ability to function as C-terminal fusions of mScarlet, again using a simple activation assay with Rgg3. Consonant with their previously determined activities as peptides alone, two of the tags, G8T and V7T, effectively activated Rgg3 (with Hill coefficients and EC_50_ values similar to SHP3), while G9N showed no activity as a fusion tag (**Fig. 5b**). In general, the results observed with peptides selected *in vivo* play out consistently in both *in vitro* and *in vivo* applications.

**Fig. 5.**
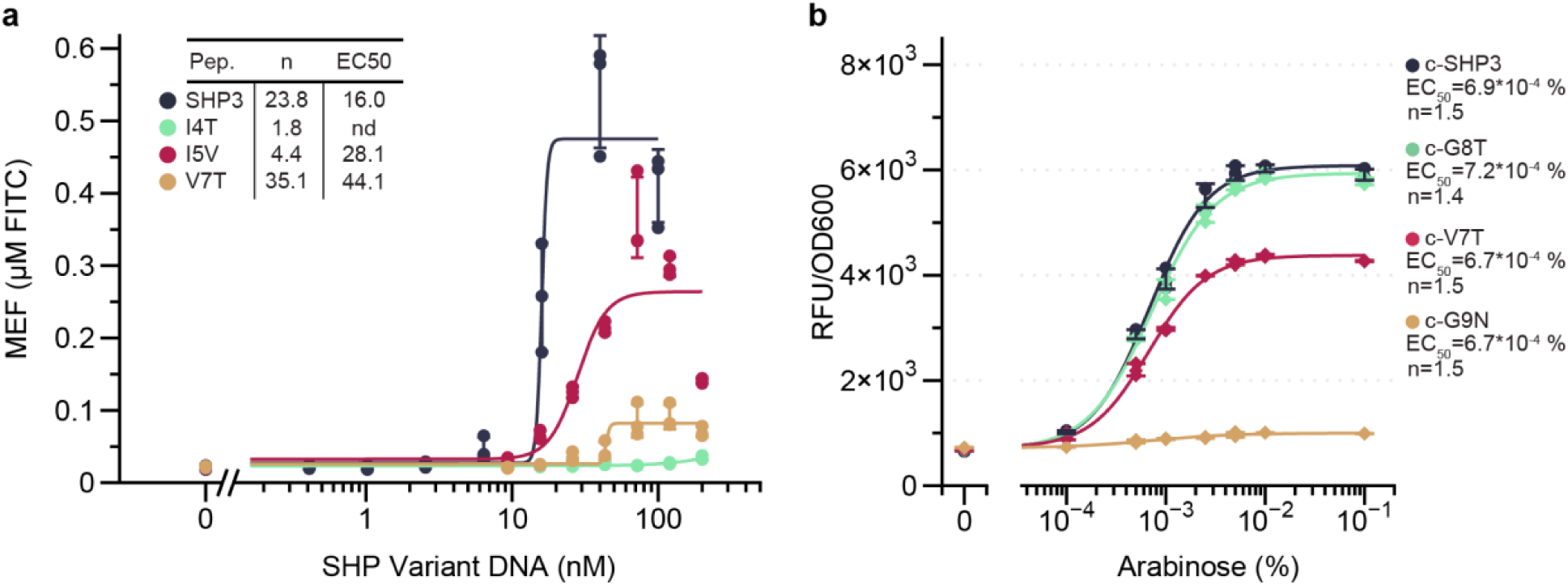
Peptide variants function as protein tags for activating gene expression. **(a)** Cell-free expression of select peptide variants under a self-inducing circuit. **(b)** *In vivo* sfGFP expression in response to a C-terminal peptide variant-tagged mScarlet. All data are carried out with *n*=3 biological replicates for *in vivo* data and experimental replicates for *in vitro* data with error represented as SD.

## Discussion

Herein, we report the adaptation of peptide-responsive TFs to both *in vitro* and *in vivo* expression systems. A key innovation in this system was engineering the peptide effector to avoid peptide export, extracellular accumulation, or re-import^26–29^, a feature that should also allow much broader application across multiple chassis. While individual transcription factor:promoter interactions remain somewhat idiosyncratic, the peptide:transcription factor interface has proven surprisingly malleable and engineerable. Peptide effectors can be fused to whole proteins, providing opportunities to measure protein synthesis and potentially leading to novel intracellular regulatory mechanisms. In particular, the use of peptide effector tags may allow *in vivo* alternatives to more cumbersome analytical methods such as ELISAs (especially since the peptide effector tags are of roughly the same size as FLAG^30^ or Myc^31^ tags). While monitoring protein expression *in vivo* is possible, it generally requires the use of fluorescent proteins that may have undesirable properties^32,33^. Self-labeling systems such as HaloTag^34^ or SNAP-tag^35^ require fusion of additional protein domains of approximately 33 or 22 kDa, respectively, while our system only requires a short (8 amino acid) peptide tag, greatly reducing any perturbations to native proteins. Taken together, these results establish programmable peptide-responsive TFs as a tool for coupling intracellular peptide production to downstream gene regulation.

The peptide-responsive regulatory circuits can potentially go well beyond commonly used regulatory parts such as LacI and TetR, especially since these ligands are not endogenously produced and often lack orthogonality (e.g., IPTG, whose induction strength is coupled to host carbon metabolism and endogenous regulatory networks)^36–38^. While commonly used repressors often rely on small molecule effectors in the micromolar range^39^, the Rgg2/3 peptide-sensing system has previously shown sensitivities to peptides at the nM level^6^. Using synthetic peptides as effectors we similarly see nM sensitivity for Rgg2, although other RRNPP TFs showed EC_50_ values on the order of ∼1-10 μM, a commonly reported range for TF-based cell-free biosensors ^40–45^. To the extent that DNA templates can be considered analytes in cell-free reactions, added DNA has EC_50_ values of 0.27 nM and 0.65 nM for peptide DNA for Rgg2 and Rgg3, respectively. The generally low concentrations required for activation suggest advantages for using endogenous peptide-responsive regulators over more conventional parts that rely on exogenous small-molecule effectors.

Transcription control over both the effector (peptide) and target (regulated cistron) provides further opportunities for generating feedback loops and cross-regulatory schemes that can be almost fully embedded within designed DNA sequences. Genetic circuits with different peptides and architectures exhibited very different Hill coefficients, from ∼2 to upwards of ∼26 (**Fig. 2a-f**). In particular, peptide-regulated circuits with self-inducing architectures had even greater ‘switch-like’ behaviors: for example, a feedback loop with wild-type SHP3 gave a Hill coefficient of 24 while for the V7T peptide the Hill coefficient was 35 (**Supplementary Fig. 12**). The wide-ranging control we have demonstrated over circuit architectures and performance finds its apotheosis in the use of SHP as a fusion tag, allowing any protein to essentially control its own (or others’) production.

An additional advantage of peptide effectors for regulating protein expression is the possibility that they can be multiplexed based on sequence recognition by Rgg3 and other TFs. In this regard, we assessed the criticality of individual residues via alanine scans with both synthetic and *in vitro* translated peptides. Further, SSM screens in cells showed that conservative mutations generally retain function and even less conservative substitutions can still activate transcription. These experiments indicate that the interface between peptides and their cognate TF is somewhat fungible, and therefore could potentially become targets for future screens and selections, especially for new, orthogonal peptide:transcription factor pairs.

The various proven advantages of using peptide-responsive regulators should prove to be a boon for metabolic engineering applications. For example, it should be possible to identify the *in vivo* concentrations of tagged proteins in a pathway (similar to **Fig. 3d-f**), and thereby develop transfer curves between inducers and enzyme amounts for rate-limiting steps. Further opportunities for developing entirely genetically encoded regulatory mechanisms are manifest and offer unique opportunities relative to small molecule induction regimes and regulatory element engineering^46–49^. The use of self-regulating circuitry (**Fig. 2d**) should allow feedback loops to be readily programmed, including using fusion proteins to control gene expression across wide swaths of cellular metabolism. Importantly, the ability to change peptide specificities and develop orthogonal peptide-responsive regulators should allow for cross-control of metabolic pathways, with peptides and proteins essentially serving as information-rich, readily programmable ‘second messengers’.

## Supporting information

Sequence Information

## Materials and methods

### Strains and growth medium

*E. coli* NEB 10-beta or NEB 5-alpha (New England Biolabs) competent cells were used for routine cloning, while DH10B was used for all *in vivo* assays. Lysogeny broth with 1.5% agar was used for routine cloning and for transformation plating in assays. Lysogeny broth supplemented with appropriate antibiotic(s) was used as the growth medium for all cloning and assay purposes. Cells were grown in Superior Broth supplemented with appropriate antibiotic(s) to prepare both chemically and electrocompetent cells. Compounds used include 20% arabinose (Teknova) diluted to the concentrations indicated in the text. Carbenicillin (100 µg/mL) and chloramphenicol (25 µg/mL) were used where appropriate.

### Plasmids and genetic parts assembly

DNA oligonucleotides for cloning were synthesized by Integrated DNA Technologies (IDT). Genes encoding all genetic parts were synthesized by Twist Bioscience as non-clonal gene fragments with adaptor sequences. Plasmids were assembled using Golden Gate cloning by combining DNA fragments with BsaI-HF v2, T7 DNA ligase, and T4 DNA ligase buffer (all New England Biolabs) in reactions. DNA parts were supplied as PCR-amplified fragments or synthetic gene blocks for the *de novo* assembly of complete vectors. Plasmid sequences were confirmed with long-read DNA sequencing (Plasmidsaurus). Relevant plasmid sequences are provided in **Supplementary Information Sequences**.

For cell-free reactions, linear expression templates (found in **Supplementary Information Sequences**) were generated for use in *E. coli* lysate experiments. Reporter and TF linear expression templates were made using PCR from constructed plasmids. Peptide producing DNA templates for lysate-based cell-free experiments were obtained via gBlock from IDT, using PCR to generate higher concentrations of linear DNA. Sixteen 50 µL PCR reactions were combined with 5x the volume of binding buffer (Buffer PB Qiagen) and added to a single Qiagen cleanup column. PCR products were then purified using Qiagen QIAquick PCR cleanup kits following manufacturer protocol and eluted twice with 30 µL of MiliQ water by applying water, centrifuging, reapplying product to the column, and spinning again. DNA concentrations were quantified by Nanodrop before being used in cell-free lysate experiments.

### Cell-based dose-response assays

Chemically competent *E. coli* DH10B cells containing the reporter plasmid were transformed with 1 µl of the regulator plasmid DNA, recovered in 100 µl of SOC medium at 37°C for 1 h, and plated onto LB agar containing chloramphenicol and carbenicillin. After overnight incubation at 37°C, three individual colonies were inoculated into 900 µl of LB containing chloramphenicol and carbenicillin in a 96 deep-well round-bottom plate (Thermo Scientific) and sealed with an AeraSeal film (Excel Scientific). Plates were incubated at 37°C with shaking at 1000 r.p.m (VWR Incubating Mini Shaker) for 16 hr.

For induction, 20 µl of each overnight culture was inoculated into 900 µl of fresh LB containing chloramphenicol and carbenicillin in a 96 deep-well plate, sealed with an AeraSeal film, and incubated at 37°C with shaking for 2 hr. Cultures were then induced with 80 µl of LB and arabinose. The total volume per well was 1 mL, with arabinose solution added to water so that the final water volume accounted for 1% (v/v) of the well volume. Cells were incubated for 4 hr at 37°C with shaking, pelleted by centrifugation, and resuspended in 900 µl of phosphate-buffered saline. A 100 µl aliquot of each resuspension was transferred to a 96-well black, clear-bottom microplate. Fluorescence using excitation/emission of 475/509 for sfGFP and 552/595 for mScarlet nm and OD were measured using a Tecan plate reader.

### SHP3 library construction and screening

Peptide libraries were constructed using mutagenic primers that have NNS at the target amino acid position. To randomize the entire peptide sequence with single-site mutations, eight primer sets were designed. As previously described, each forward primer contained a region of sequence homology to the plasmid backbone for annealing, an NNS codon for diversification, and a BsaI recognition site^50^. The corresponding reverse primers contained a homologous region for reverse annealing and a BsaI recognition site. PCR amplification with these primers produced linear fragments containing a single randomized codon flanked by BsaI sites that generate complementary overhangs upon digestion. Purified PCR products were assembled into circular plasmids using Golden Gate cloning. The assembled libraries were purified using the Monarch PCR & DNA Cleanup Kit (New England Biolabs) and transformed into electrocompetent *E. coli* cells containing the reporter plasmid. Following electroporation, cells were recovered in SOC medium for 1 hr at 37°C with shaking, then inoculated into LB and grown overnight at 37°C for 16 hr. Before inoculation, an aliquot of each reaction was serially diluted and plated to determine total library size. Library sizes ranged from approximately 1x10⁵ to 1x10⁶ transformants. After overnight growth, glycerol stocks of each library were prepared, flash-frozen in liquid nitrogen, and stored at -80°C.

To isolate peptide variants spanning a range of activities, mutagenized libraries were screened using two colony-selection stringencies. In both screens, libraries were diluted 10², 10⁴, and 10⁶ in LB medium, and 100 µl of each dilution was plated onto LB-agar plates supplemented with 0.005% arabinose or no arabinose using sterile plating beads. This enabled comparison of colony fluorescence under inducing and non-inducing conditions. Plates were incubated at 37°C for 16 h, after which colonies were visualized under blue light illumination. For the high-stringency screen, the brightest fluorescent colonies were selected visually, cultured overnight in LB at 37°C with shaking, and quantitatively assayed for reporter output in the presence and absence of 0.005% arabinose as described in the “*in vivo* dose-response assays” section. Variants showing the largest fold-change responses were sequenced, subcloned, and assayed across an arabinose series to determine theur full activation profiles.

For the lower-stringency screen, dimmer fluorescent colonies were intentionally selected to recover variants with moderate activity. These colonies were similarly cultured and quantitatively assayed, and variants with fold changes of approximately 2.5-4.5-fold changes were sequenced. Variants containing D2* mutations or a single-nucleotide deletion that generated an extended open reading frame were excluded from further analysis. Non-redundant peptide sequences were then subcloned into the expression vector and assayed across an arabinose dose-response series to determine their full activation profiles.

### Linear DNA preparation

Linear DNA was prepared by PCR from gBlocks from IDT or plasmids. For each preparation, a minimum of eight and a maximum of 96 50 μL PCR reactions were prepared using the NEB Phusion HiFi PCR Protocol. PCR products were purified using the Qiagen QIAquick columns and pooled with 5x the volume of binding buffer (Buffer PB, Qiagen) was added to the reactions. The volumes were run through purification columns (16 PCR reactions’ worth of volume per column) and washed with 2x with Buffer PE (Qiagen) using a manifold system. Columns are then spun down at maximum speed for 1 min to remove residual wash buffer, then rehydrated with 30 μL of MiliQ water per column, and spun again in a fresh Eppendorf tube to collect. Effluent was then pooled again, and the DNA concentration in the total volume was quantified using a NanoDrop. This results in linear DNA template concentrations ranging from 600 to 1500 ng/μL.

### E. coli extract preparation

*E. coli* BL21(DE3)* was used for lysate preparation. Preparation of *E. coli* BL21(DE3)* extract was carried out using previously published methods^16^. Briefly, BL21(DE3)* cells were plated on LB agar, and a single colony was cultured overnight in 2xYT+P media, followed by inoculation into 1 L of 2xYT+P media. One liter of this medium uses 16 g tryptone, 10 g yeast extract, 5 g NaCl, 7 g potassium phosphate dibasic, and 3 g potassium phosphate monobasic. Cells were then grown with shaking at 220 rpm at 37°C and harvested at an OD of 3.0, after approximately 4 hr. Once an OD of 3.0 was reached, the cells were processed in accordance with the previous protocol up through dialysis^16^.

### Cell-free reaction preparation

All cell-free reactions contained in the paper were set up as a modified phosphoenolpyruvate, amino acids, NAD+, and oxalic acid (PANOx)^17^ protocol using 30% (v/v) cell extract, 30% (v/v) PANOx reaction components, and 40% (v/v) water with additions, such as peptide and DNA. The reaction components are made up of concentrated stock solutions: 15X salt solution, a 15X transcriptional “master mix,” a 12.5X cofactor “reagent mix,” 50 mM amino acids, and 1 M phosphoenolpyruvate (PEP) solutions. The following concentrations per cell-free reaction are 8 mM salt solution, containing 8 mM magnesium glutamate, 10 mM ammonium glutamate, and 130 mM potassium glutamate. The transcription master mix contains 1.2 mM ATP, 0.850 mM GTP, 0.850 mM UTP,0.850 mM CTP, 72 μM folinic acid, and 0.171 mg/mL tRNA. Reagent mix cofactor solution contains 0.33 mM NAD, 0.27 mM CoA, 4 mM oxalic acid, 1 mM putrescine, 1.5 mM spermidine, 57 mM HEPES.^16^ The amino acid solution contains 2mM of each amino acid. Finally, 30 mM PEP solution is used as an energy source.

### Cell-free biosensor assay reactions

Cell-free master mix reactions were created as described in cell-free reaction preparation. To create biosensor reactions, 70 uL overnight CFPS reactions were constructed using 10 nM TF linear expression templates. These TF CFPS reactions were then placed in a 2 mL eppendorf tube and incubated in a Vortemp without shaking for 20 hr at 30 °C. For biosensor assays in **Fig. 1**, fresh cell-free reactions were constructed using 9% (v/v) of TF CFPS and 50 nM reporter DNA linear expression template with the exception of **Fig. 1b**, where 30 nM reporter template was used before sensor optimization found in **Supplementary Fig. 2**. To induce biosensor expression, synthetic peptide was added to the reactions at 1 mg/mL for **Fig. 1b**, 0.3 mg/mL for **Fig. 1c, 1d**, or at 0, 5.98x10^-5^, 0.0022, 0.025, 0.27, 3.0, 112.5, and 375 µM synthetic peptide concentrations for dose response curves in **Fig. 1e-j**. Unless stated otherwise, all proceeding reactions were constructed using the same TF CFPS percentage with 50nM reporter template and varied peptide producing DNA or synthetic peptide concentrations. All cell-free reactions were run on a plate reader for 10 h at 30 °C using excitation/emission 485/520 for GFP and 572/596 for mScarlet.

### Hill equation fits

Dose response curves were fit using the Datagraph arbitrary fit function using a Hill function of the form:

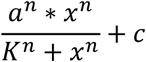

where n represents the Hill coefficient, K represents the EC_50_ of peptide or DNA inducer, a represents the upper asymptote, and c is the baseline expression.

## Data availability

Data for all main text and supplementary information figures are provided in Supplementary Excel files.

## Code availability

Code used for data analysis, along with data tables, can be found at https://github.com/LucksLab/Brown_ElSayed_Peptide_Biosensors_2026

## Contributions

A.D.E. conceived the project. A.C.E., D.M.B., B.G.H., T.M.M., and I.M.H curated the data. A.C.E. and D.M.B. carried out the investigations, devised the methodology, and carried out the formal analysis. A.D.E and J.B.L. supervised the project. A.C.E. and D.M.B. wrote the original draft. A.C.E., D.M.B., J.B.L., and A.D.E. reviewed and edited the manuscript.

## Acknowledgements

This work was supported by the National Science Foundation (2310382 to J. B. L.). The views, opinions, and/or findings expressed are those of the authors and should not be interpreted as representing the official views or policies of the Department of Defense, the National Science Foundation, or the U.S. Government. Research reported in this publication was supported by the National Institutes of Health (under award number T32GM139796 to A.C.E. and 1R01 GM146093-01 A1 to A.D.E). The content is solely the responsibility of the authors and does not necessarily represent the official views of the National Institutes of Health. The authors acknowledge financial support from the Welch Foundation (F-1654 to A.D.E.).

