## Supplementary Information for "Design and characterization of peptide-responsive biosensors"

#### Supplementary Figures

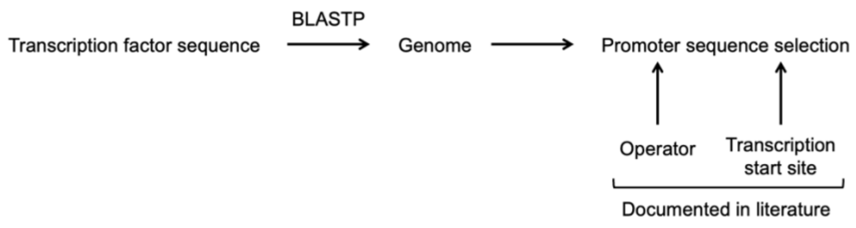

**Supplementary Figure 1. Generation of transcription factor-responsive promoter sequences.** TF protein sequences were searched against the database using BLASTP to identify genomes containing the TF. Based on known operators and transcription start sites in the literature, the promoter sequences were extracted.

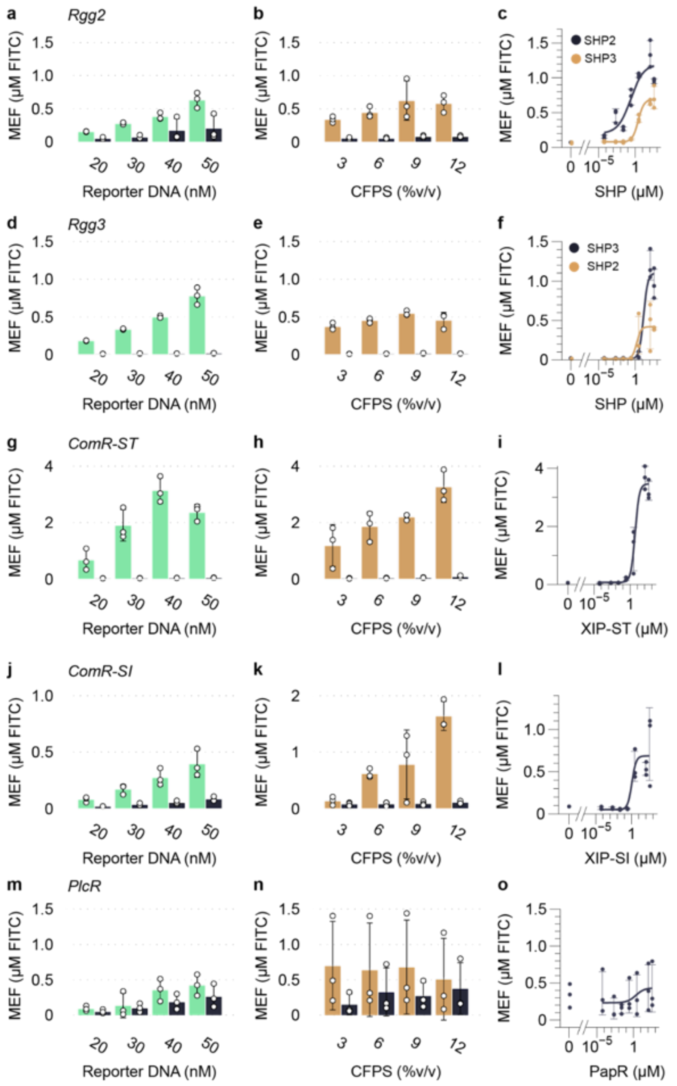

**Supplementary Figure 2. Two pot optimizations of RRNPP-family biosensors. (a)** Rgg2 reporter DNA titration. **(b)** Rgg2 overnight CFPS titration. **(c)** Rgg2 dose-response curves for SHP2 and SHP3 ligands. **(d)** Rgg3 reporter DNA titration. **(e)** Rgg3 overnight CFPS titration. **(f)** Rgg3 dose-response curves for SHP2 and SHP3 ligands. **(g)** ComR-ST reporter DNA titration. **(h)** ComR-ST overnight CFPS titration. **(i)** ComR-ST XIP-ST dose response curve. **(j)** ComR-SI reporter DNA titration. **(k)** ComR-SI overnight CFPS titration. **(l)** ComR-SI XIP-SI dose-response curve. **(m)** PlcR reporter DNA titration. **(n)** PlcR overnight CFPS titration. **(o)** PlcR PapR dose response curve. For **(a, d, g, j, m)** 9 %v/v aTF CFPS and 1 mg/mL cognate peptide was used in experiments. For **(b,e,h,k,n)** 1 mg/mL cognate peptide and 50 nM reporter DNA were used in experiments. For **(c,f,i,l,o)** 9 %v/v aTF CFPS and 50nM reporter DNA was used for each dose-response curve.

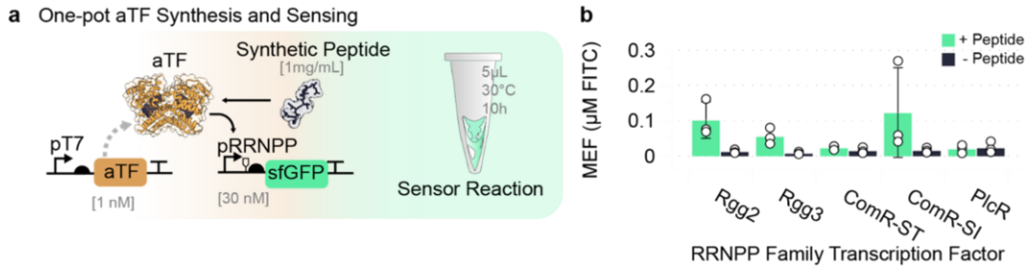

**Supplementary Figure 3. One-pot transcription factor synthesis and sensing show biosensor activity. (a)** Schematic of the one-pot transcription factor synthesis regime. **(b)** Endpoint fluorescence of biosensor reactions for Rgg2, Rgg3, ComR-ST, ComR-SI, and PlcR. 1 nM of pT7-aTF plasmid was added to cell-free reactions with 30 nM pRRNPP reporter DNA and 1 mg/mL cognate peptide. Reactions were then run on a plate reader for 10 hr, with reads taken every 5 min at excitation/emission of 485/520. One-pot aTF synthesis and peptide sensing gave variable results, with lower fluorescence than pre-synthesized aTF. ComR-ST, ComR-SI, and PlcR are notably not reliably functional through one pot synthesis mechanisms; however, ComR-ST and ComR-SI are functional using the overnight CFPS approach.

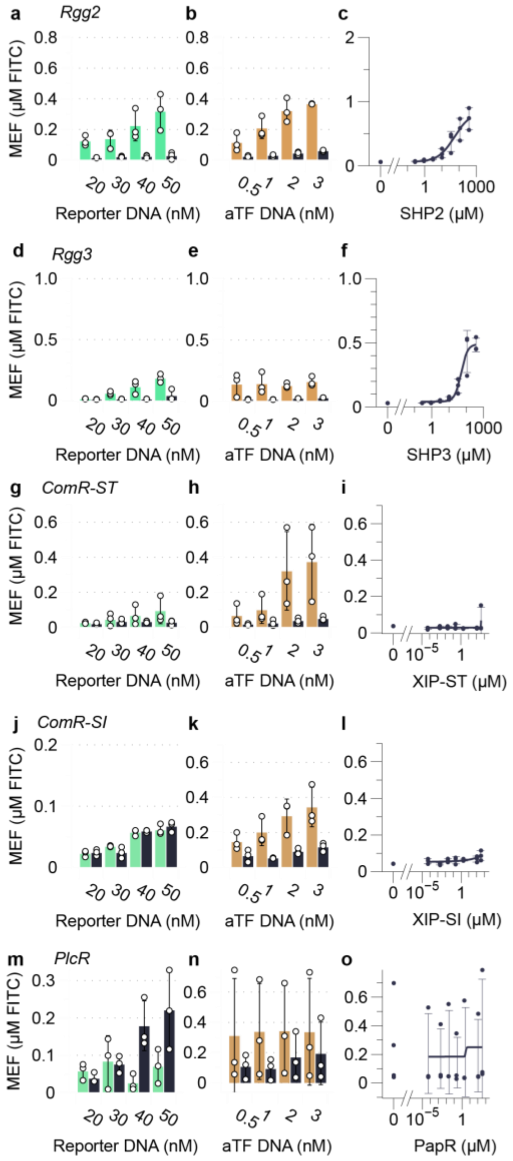

**Supplementary Figure 4.** One-pot optimizations. **(a)** Rgg2 reporter DNA titration. **(b)** pT7-Rgg2 DNA titration. **(c)** Rgg2 dose response curves for the SHP2 ligand. **(d)** Rgg3 reporter DNA titration. **(e)** pT7-Rgg3 DNA titration. **(f)** Rgg3 dose response curves for the SHP3 ligand. **(g)** ComR-ST reporter DNA titration. **(h)** pT7-ComR-ST DNA titration. **(i)** ComR-ST XIP-ST dose response curve. **(j)** ComR-SI reporter DNA titration. **(k)** pT7-ComR-SI DNA titration. **(l)** ComR-SI XIP-SI dose response curve. **(m)** PlcR reporter DNA titration. **(n)** pT7-PlcR DNA titration. **(o)** PlcR PapR dose response curve. For **(a, d, g, j, m)** 1 nM aTF DNA and 1 mg/mL cognate peptide was used in experiments. For **(b,e,h,k,n)** 1 mg/mL cognate peptide and 50 nM reporter DNA was used in experiments. For **(c,f,i,l,o)** 2 nM aTF DNA and 50nM reporter DNA were used for each dose response curve.

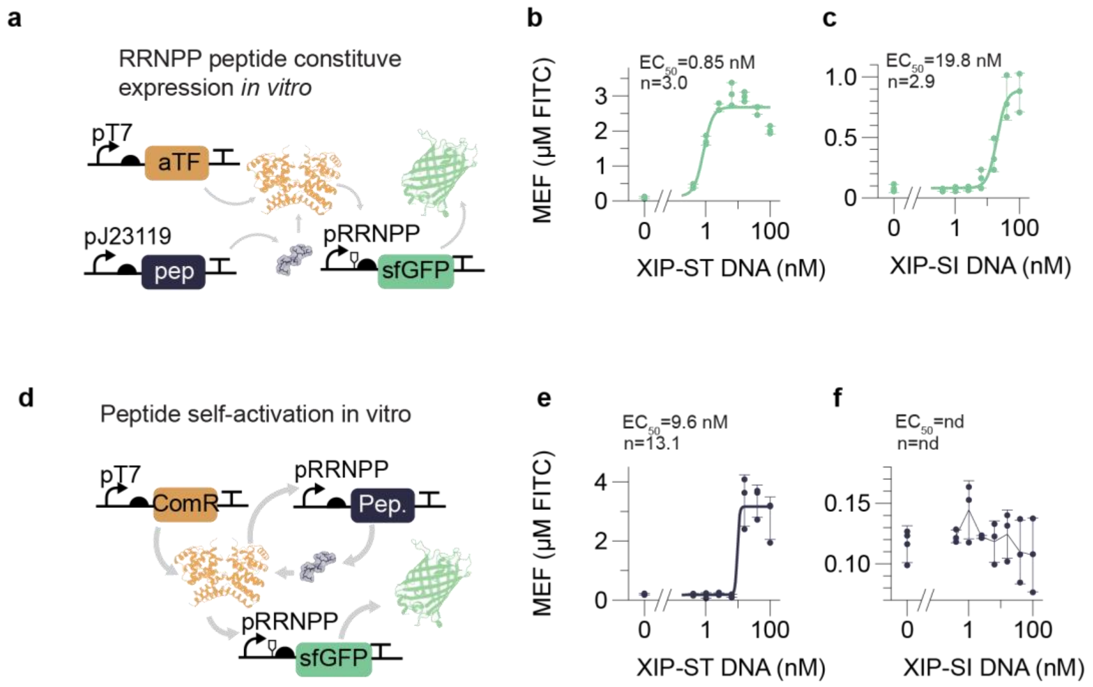

**Supplementary Figure 5.** XIP-SI and XIP-ST expression with cognate transcription factor *in vitro*. **(a)** Schematic of constitutively expressed peptide ligands for activation of ComR transcription factors. **(b)** Dose response of ComR-ST with pJ23119-XIP-ST DNA. **(c)** Dose response of ComR-SI with pJ23119-XIP-SI DNA. **(d)** Schematic of self-inducing peptide expression for activating ComR transcription factors. **(e)** Dose response of ComR-ST with pComR-ST controlled XIP-ST expression. **(f)** Dose response of ComR-SI with pComR-SI controlled XIP-SI expression. All reactions contained 9% (v/v) ComR aTF CFS, 50 nM pComR-sfGFP reporter DNA, and 0, 0.4, 1.0, 2.56, 6.4, 16, 40, and 100 nM XIP DNA. All data were carried out with *n*=3 experimental replicates and averages were fitted with a Hill function to determine $EC_{50}$ and Hill coefficient values. Error is represented by standard deviation. Reactions were run on a plate reader for 10 hr at 30 °C using excitation/emission of 485/520.

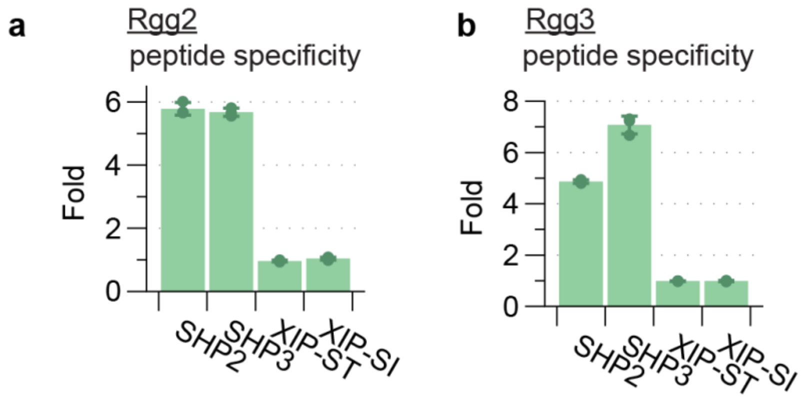

**Supplementary Figure 6.** Rgg2 and Rgg3 cross-activation with XIP *in vivo*. **(a)** Rgg2 ligand specificity *in vivo*. **(b)** Rgg3 ligand specificity *in vivo*. Both TFs activate gene expression in the presence of SHP2 and SHP3, with no activation shown when exposed to XIP peptides. Cells were induced with either 0% or 0.005 % (v/v) arabinose for four hr at 37 °C before sfGFP fluorescence was measured using excitation/emission of 475/509, and $OD_{600}$ was measured. Fold change was calculated by dividing the normalized fluorescent (RFU/OD) values of the induced sample by the uninduced sample.

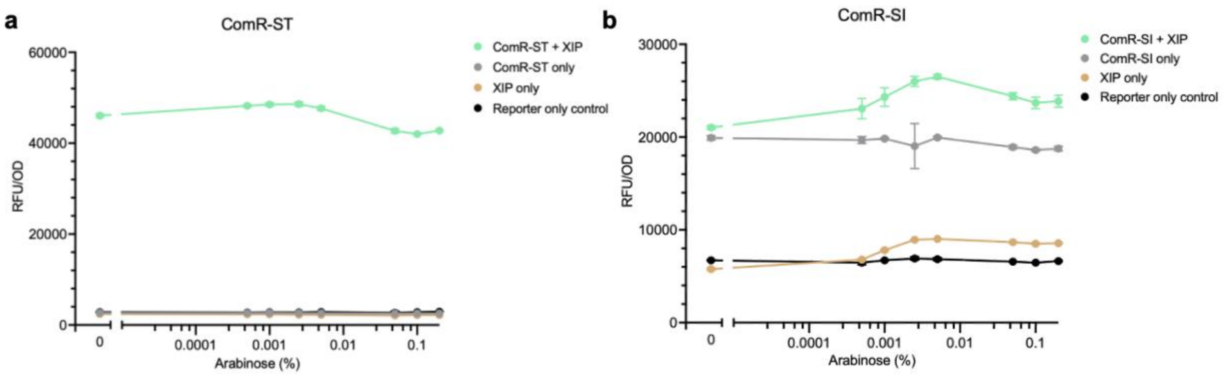

**Supplementary Figure 7. ComR-ST and ComR-SI assays *in vivo*.** The pBAD-driven peptide activates the TF, which in turn regulates sfGFP expression from the reporter plasmid. **(a)** ComR-ST activation *in vivo.* **(b)** ComR-SI activation *in vivo.* Conditions include TF and peptide (green), TF only (grey), peptide only (gold), and reporter-only control (black). Cells were induced with varying levels of arabinose for four hr at 37 °C before sfGFP fluorescence was measured using excitation/emission of 475/509, and $OD_{600}$ was measured.

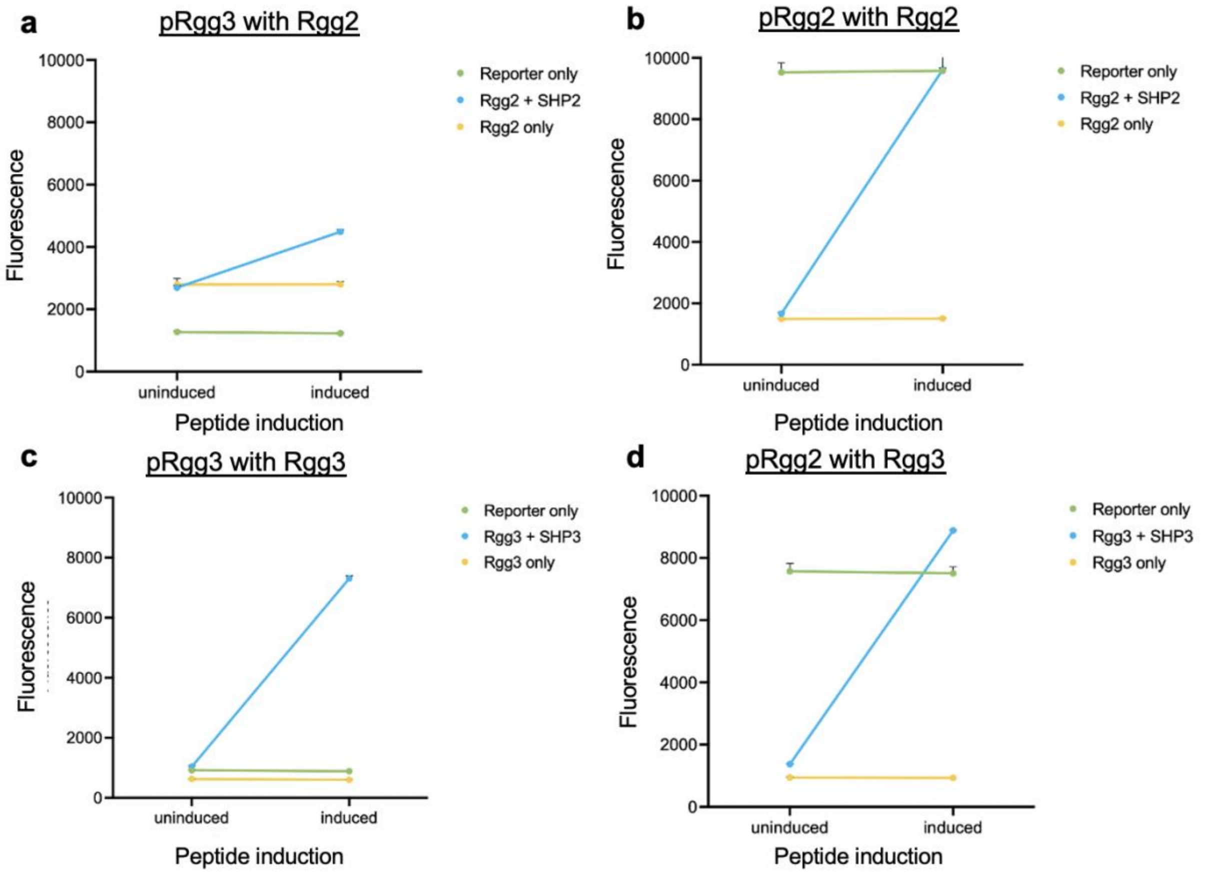

**Supplementary Figure 8. Basal-level expression of pRgg2 and pRgg3 *in vivo* and response to Rgg2 and Rgg3.** *In vivo* expression of Rgg2 with **(a)** pRgg3 and **(b)** Rgg2 promoters. *In vivo* expression of Rgg3 with **(c)** pRgg3 and **(d)** Rgg2 promoters. pRgg2 exhibits basal activity in the absence of a TF, yet both transcription factors alone can repress it. Upon induction of the peptide, the promoter is turned on. pRgg3 shows no basal activity in the absence of a TF, minimal activity with the TF, and activation upon induction of the peptide with the TF.

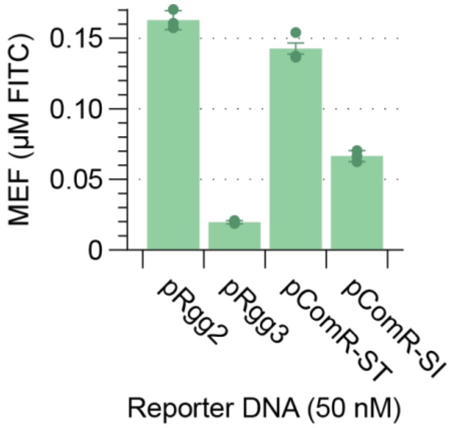

**Supplementary Figure 9. Background reporter levels in the absence of RRNPP transcription factors**. 50nM of reporter DNA for each RRNPP family member was added to fresh cell-free reactions to assess background levels of expression independent of the presence of aTF and ligand. All data are carried out with *n*=3 experimental replicates. Error is represented by standard deviation. Reactions were run on a plate reader for 10 hr at 30 °C using excitation/emission of 485/520 nm.

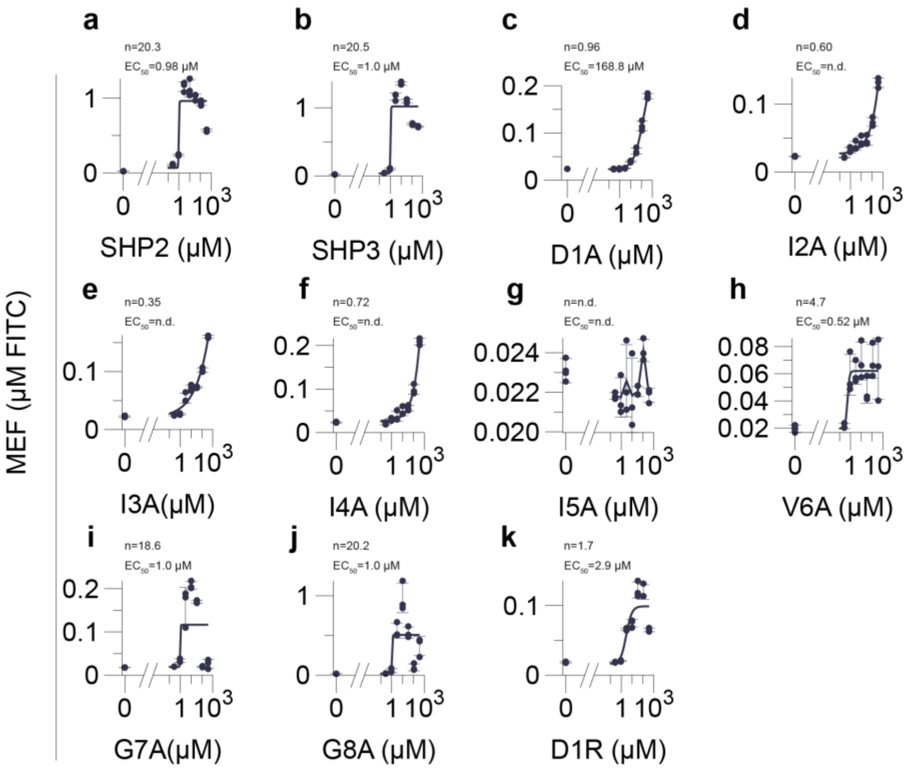

**Supplementary Figure 10. Synthetic peptide alanine scan dose response curves for Rgg3**. **(a-k)** Dose responses curves for Rgg3 with synthetic peptide alanine scan variants compared to wt SHP2 and SHP3 sequences. All reactions contained 9% (v/v) Rgg3 aTF CFS, 50 nM pRgg3-sfGFP reporter DNA, and 0 to 375 μM synthetic peptide. All data are carried out with *n*=3 experimental replicates and averages are fitted with a Hill function to determine $EC_{50}$ and Hill coefficient values. Error is represented by standard deviation. Reactions were run on a plate reader for 10 hr at 30 °C using excitation/emission of 485/520 nm.

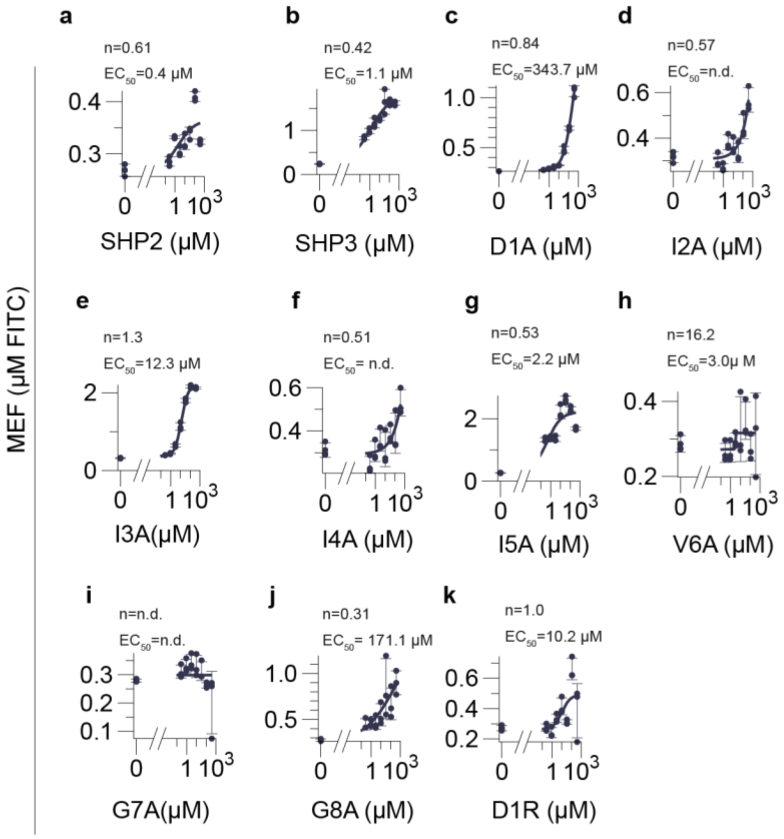

**Supplementary Figure 11. Synthetic peptide alanine scan dose response curves for Rgg2. (a-k)** Dose responses curves for Rgg2 with synthetic peptide alanine scan variants compared to wt SHP2 and SHP3 sequences. All reactions contained 9% (v/v) Rgg2 aTF CFS, 50 nM pRgg2-sfGFP reporter DNA, and 0 to 375 µM synthetic peptide. All data are carried out with *n*=3 experimental replicates and averages are fitted with a Hill function to determine $EC_{50}$ and Hill coefficient values. Error is represented by standard deviation. Reactions were run on a plate reader for 10 hr at 30 °C using excitation/emission of 485/520 nm.

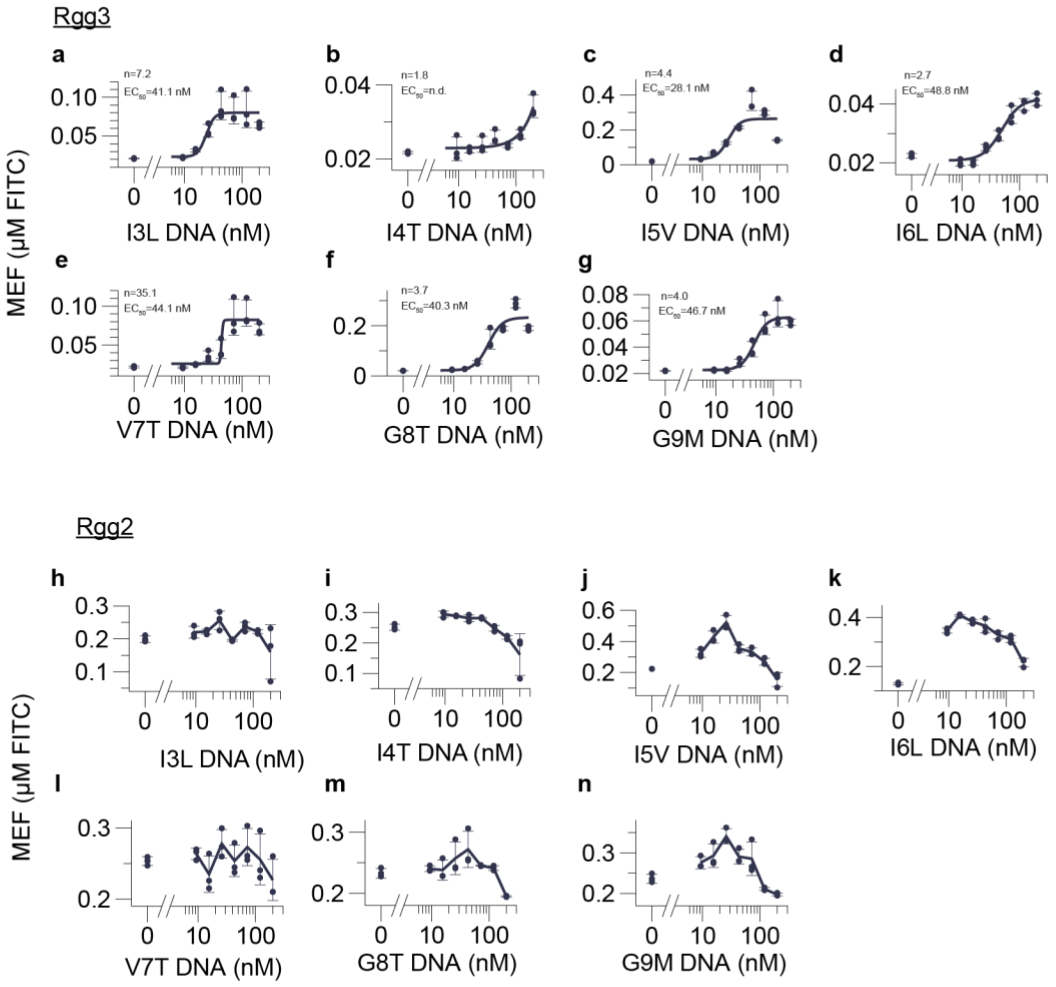

**Supplementary Figure 12. Self-induction of SSM peptide variants *in vitro*. (a-g)** Rgg3 dose response curves over peptide variant DNA concentrations. **(h-n)** Rgg2 dose response curves over peptide variant DNA concentrations. Peptide DNA for both Rgg3 and Rgg2 dose response curves contains the cognate promoter of Rgg3 (pRgg3) while the reporter DNA for Rgg3 and Rgg2 dose response curves contain their respective cognate promoters (pRgg3, pRgg2 respectively). All reactions contained 9% (v/v) Rgg2/3 aTF CFS, 50 nM pRgg2/3-sfGFP reporter DNA, and 0 to 200 nM peptide variant DNA under the pRgg3 promoter. All data are carried out with *n*=3 experimental replicates and averages are fitted with a Hill function to determine $EC_{50}$ and Hill coefficient values. Error is represented by standard deviation. Reactions were run on a plate reader for 10 hr at 30 °C using excitation/emission of 485/520 nm.

### Supplementary Tables

**Supplementary Table 1. Summary of sequences used in this study.**

| Transcription factor | Organism | Transcription factor | Promoter sequence | Transcription start site | Regulated gene | Operator |
|---|---|---|---|---|---|---|
| Rgg2 (GCF_009650275.1) | *Streptococcus dysgalactiae* | MDKELGKTLRRLRQGKQVSISSLADEHLSKSQISRFERGESEISCSRLLNLLDKLNITIDEFVSTHSKTHTHFFTLLSRVRKYYAEKNVAKLLKLLEDYAHKDYESTMIKAILSSIEPTVEPSEEEVTRLTDYLFSVEQWGYYEIILLGNCSRFINYNTLFLLTKEMVTSFAYSEQNKTNKTLVTQLSINCLIISIDYSYFDHSHYLIEKIEFLLRDELNFYEKTVFLYVHGYYKLKQGQVSGKDDMRQALQIFKYLGEDALYYSYKEHYRKEVLGDEEDEEEG | TGTTTTCCTTGGCGTAGTCGCCTTAAGGTTTTACCGAGTTCTTTTTCCATTTTTCCCACTTTCACAACAAAAAATTCAAATCCGTCAAATCCATTATATAATACAAGTA | AAGTA[6] | SHP2[6] | TTTTCCCACTTTCACAACAAAAA[42] |
| Rgg3 (GCF_000011825.1) | *Streptococcus thermophilus* | MKSKLGSTLRKVRNGKQISICSVADEHLSKSQISRFERGESEISCIRLINILDKLHITLDEFLILHDEDYTKTESFANLVQYIRKQYSLQNINNIQSLLSDSSNYTLDPFEKTMVKSILHTMDSSIIPSDDELLQLADYLFKVEKWGYY | TATGGCTTTCCTTTTCGAACTTTTCTAAGTGTGACACCAAGTTTTGATTTCATATTTTCCCACTTTCCCAACAAAAATCAATTCCGTTTTTTTTTATTG | ATAAA[6] | SHP3[6] | TTTTCCCACTTTCCCAACAAAAA[42] |

| | | | | | | |
|---|---|---|---|---|---|---|
| | | EIILLGNCVRTIDYNSVFLLTKEMLNNYIYSSLNKTNKRIVTQLAINCLILSIDMEEFTNCFYLIDEIKALLDNELNFYEQTVFLYATGYFEFKRWQSTSGIEKMKQAIQVLDILGEDNLKLHYTIHFDKLINNK | TACAATAAAACG | | | |
| ComR-ST (GCF_000014485.1) | *Streptococcus thermophilus* | MNLKDSIGLRIKTERERQQMSREVLCLDGAELTVRQLIRIEKGESLPSLDRLSYIAKRLGKSMTELLDQDNITIPDEYYEMKNRLIKFPTYRNPDRIKSKLTLIEEVYEKFFDILPEEELLTLDILENILSFTSWEESPKVEEIYEDLFEQVKRKRKFSTNDLLVIDYYFFHLYGRKQYDKKLFERIIKRVLNQEIWTDDVYNIVLFNDLMAIAALKIFHNSFSDFLTVVDKALAVIEKSQLYSYKPSVFVLKAKYELLHKENKKEAAENYDKAIVFASVLEDSVLEESIKAGKLADGL | GTGGTTTTGACTCTATCTCGCTTATTTAATTGTTTTAGTCAAGTTATTTATAGTGACATATATGTCTCTATTTTATTTTTCTACATTCTTTTATAATAGGTCTA | AGGTC[43] | ComX[44] | TAGTGACATATATGTCTCTA[43] |
| ComR-SI (GCA_000246835.1) | *Streptococcus infantarius* | MEKFGEKIRLMREEKGISREEFCGDETELSVRQLARIELNQSIPNLSKASFIANRLGVKLGTLTDGDSLELPKRYKELKYLLLRTPTYGDQVR | AAGCGGTAGACGGAGAGCAACTCAGATGAGTTGCTTTTTTTGTGCATCAATTTGGACATTTTTGT | AACGT (Used in this reporter construct) | ComX[44] | TTTGGACATTTTTGTCATGT |

| | | | | | | |
|---|---|---|---|---|---|---|
| | | LDRKNDYFDEI AEVFYDVIPEE ERLIIDCLQSK FDVHFSEDVN FGEGILNDYF GQVNRKKVFT INDLILIDLYFA CLSSAKKFEGI YSLDFYDDLM KRLVNQKHVS PETDLILNNVL LNNIDLAFQFK REYYIERVITIS EKIMTEIHDFQ RRPILSLVEW KYYLKLKHDF ALAEQSFTNA TLFARLVGDT YLENKLKEEW QLDIDAVE | CATGTTC CTTTTCC CTAATCC TTTCTATA ATAAGGG TA | | | |
| PlcR (GCF_0003067 45.1) | *Bacillus thuringiensis* | MQAEKLGSEI KKIRVLRGLTQ KQLSENICHQ SEVSRIESGA VYPSMDILQGI AAKLQIPIIHFY EVLIYSDIERK KQFKDQVIML CKQKRYKEIY NKVWNELKKE EYHPEFQQFL QWQYYVAAY VLKKVDYEYCI LELKKLLNQQL TGIDVYQNLYI ENAIANIYAEN GYLKKGIDLFE QILKQLEALHD NEEFDVKVRY NHAKALYLDS RYEESLYQVN KAIEISCRINS MALIGQLYYQ RGECLRKLEY EEAEIEDAYKK ASFFFDILEMH AYKEALVNKIS R | AAATGAA AGTGTAT TCATTCAT TATATTCA CTGTGTA TAAAGTTA TAATGATA TGAACAT TTGCATAT TTTAATTT AGTGATA GAAATTT CGTGAAA GGTGGGA TATTCTAG TCATA | AACGT (Used in this reporter construct) | Phospholip ase C[44] | TATGAAC ATTTGCA TA[45] |

**Supplementary Table 2. Peptide variants identified in the high-fluorescence screen.** Corresponding fold-changes in activation upon arabinose induction and additional plasmid mutations identified through sequencing are shown.

| Library position | Plasmid I.D. | Fold-change | Mutation | Additional mutations |
|---|---|---|---|---|
| 2 | pIAC234 | 8.3 | D2R | Mutation in ampR |
| 2 | pIAC235 | 8.5 | D2A | |
| 3 | pIAC236 | 8.9 | I3L | |
| 3 | pIAC237 | 9.0 | I3L | |
| 4 | pIAC238 | 8.5 | I4T | |
| 4 | pIAC239 | 7.9 | I4T | |
| 5 | pIAC240 | 1.0 | I5V | Mutation in araC |
| 5 | pIAC241 | 8.1 | I5L | |
| 5 | pIAC242 | 8.1 | I5L | Mutation in Rgg3 |
| 6 | pIAC243 | 9.1 | I6L | |
| 6 | pIAC244 | 8.8 | I6L | |
| 7 | pIAC245 | 8.9 | V7T | |
| 8 | pIAC246 | 8.7 | G8T | |
| 8 | pIAC247 | 9.8 | G8T | |
| 8 | pIAC248 | 9.8 | G8H | |
| 9 | pIAC249 | 9.4 | G9M | |
| 9 | pIAC250 | 9.4 | G9I | |

**Supplementary Table 3. Peptide variants identified in the moderate-fluorescence screen.** Corresponding fold-changes in activation upon arabinose induction and additional plasmid mutations identified through sequencing are shown.

| Library position | Plasmid I.D. | Fold-change | Mutation | Additional mutations |
|---|---|---|---|---|
| 2 | pIAC381 | 2.2 | D2* | |
| 2 | pIAC382 | 2.1 | D2* | |
| 2 | pIAC383 | 3.6 | D2L | Mutation in Rgg3 |
| 2 | pIAC384 | 2.6 | D2G | araC mutation |
| 3 | pIAC385 | 2.9 | I3R | |
| 3 | pIAC386 | 2.9 | I3R | |
| 3 | pIAC387 | 3.9 | I3S | |
| 4 | pIAC388 | 2.4 | I4D | |
| 4 | pIAC389 | 2.6 | I4D | |
| 4 | pIAC390 | 3.4 | I4E | |
| 5 | pIAC391 | 3.6 | I5F | |
| 5 | pIAC392 | 3.6 | I5F | |
| 5 | pIAC393 | 3.7 | I5F | |
| 5 | pIAC394 | 2.4 | I5A | |
| 6 | pIAC395 | 3.8 | I6T | |
| 6 | pIAC396 | 2.1 | I6T | |
| 6 | pIAC397 | 3.3 | I6F | |
| 7 | pIAC398 | 2.6 | V7S | |
| 7 | pIAC399 | 2.6 | V7M | |
| 7 | pIAC400 | 2.6 | V7M | |
| 8 | pIAC401 | 3.5 | G8W | |
| 8 | pIAC402 | 2.4 | Single nucleotide deletion | |
| 8 | pIAC403 | 3.1 | G8W | |
| 9 | pIAC404 | 4.3 | G9N | |
| 9 | pIAC405 | 4.2 | G9N | |
