## Supplementary material for "Design and characterization of peptide-responsive biosensors": Sequence Information

### Sequence Table 1. Summary of sequences used for in vitro experiments in this study.

#### Feature key

| Colour | Feature class | Annotations assigned to this class |
|---|---|---|
| | Point mutation (peptide variant) | D2A · G8A · G8T · G9A · G9M · I3A · I3L · I4A · I4T · I5A · I5V · I6A · I6L · V7A · V7T |
| | Signalling-peptide CDS | SHP2 Cognate Peptide · SHP3 Cognate Peptide · XIP-SI Peptide · XIP-ST Peptide |
| | Operator / TF binding site | Rgg binding site |
| | Core promoter element (−10, TSS) | -10 · Transcription start site |
| | Ribosome binding site | RBS · RBS (strong) |
| | Start codon (ATG) | Start codon (ATG) |
| | Promoter | ComR-SI promoter · ComR-ST promoter · J23119 promoter · PlcR promoter · Rgg2 promoter · Rgg3 promoter · T7 promoter |
| | Transcription factor CDS | ComR-SI CDS · ComR-ST CDS · PlcR CDS · Rgg2 CDS · Rgg3 CDS |
| | Reporter / fusion protein CDS | mScarlet-I · sfGFP |
| | mRNA stability hairpin / structural RNA | 5S rRNA · pHP14 mRNA stability hairpin |
| | Assembly scar / MCS | BamHI-BglII Scar · MCS (fragment) |
| | Transcriptional terminator | T7 terminator · rrnB T1 terminator · rrnB terminator (TrrnB) |
| | Vector backbone (ori, markers, lac) | AmpR · AmpR promoter · CAP binding site · KanR · SET1 (fragment) · bom (fragment) · lac operator · lac promoter · lacI (fragment) · lacZα (fragment) · ori · rop (fragment) |

| # | Type | Name and annotated features | Sequence (5′→3′) |
|---|---|---|---|
| **A. Transcription factor expression plasmids (pT7-aTF)** | | | |
| 1 | Plasmid | **pT7-Rgg2**<br>*Graham_Hendrickson_B6P_1_Rgg2_pLann · 2679 bp · circular*<br>Fig. 1; Suppl. Figs 2–4<br><br>**Annotated features**<br>T7 promoter 575–593<br>RBS 625–647<br>Rgg2 CDS 655–1509<br>SET1 (fragment) 1579–1628<br>T7 terminator 1636–1683<br>KanR 1701–2516<br>ori 2638–2679, 1–547 | 1 AAAAAAACCACCGCTACCAGCGGTGGTTTGTTTGCCGGATCAAGAGCTACCAACTCTTTTTCCGAAGGTAACTGGCTTCA<br>81 GCAGAGCGCAGATACCAAATACTGTTCTTCTAGTGTAGCCGTAGTTAGGCCACCACTTCAAGAACTCTGTAGCACCGCCT<br>161 ACATACCTCGCTCTGCTAATCCTGTTACCAGTGGCTGCTGCCAGTGGCGATAAGTCGTGTCTTACCGGGTTGGACTCAAG<br>241 ACGATAGTTATCGGATAAGGCGCAGCGGTCGGGCTGAACGGGGGGTTCGTGCACACAGCCCAGCTTGGAGCGAACGACCT<br>321 ACACCGAACTGAGATACCTACAGCGTGAGCTATGAGAAAGCGCCACGCTTCCCGAAGGGAGAAAGGCGGACAGGTATCCG<br>401 GTAAGCGGCAGGGTCGGAACAGGAGAGCGCACGAGGGAGCTTCCAGGGGGAAACGCCTGGTATCTTTATAGTCCTGTCGG<br>481 GTTTCGCCACCTCTGACTTGAGCGTCGATTTTTGTGATGCTCGTCAGGGGGGCGGAGCCTATGGAAAAACGCCAGCAACG<br>561 CGATCCCGCGAAATTAATACGACTCACTATAGGGTGTCCACAACGGTTTCCCTCTAGAAATAATTTTGTTTAACTTTAAG<br>641 AAGGAGATATACATATGGACAAAGAGCTTGGTAAGACCCTGCGCCGCCTGCGCCAAGGAAAACAAGTTAGTATATCCAGC<br>721 CTTGCGGACGAGCATTTATCGAAAAGTCAAATCTCGCGTTTTGAGCGAGGAGAGTCAGAAATTAGTTGTTCTCGGCTTTT<br>801 AAACTTATTGGACAAACTTAACATCACAATAGACGAGTTTGTATCTACGCACAGCAAAACCCACACGCACTTCTTTACGC<br>881 TCCTTTCCCGTGTCCGGAAGTACTATGCGGAAAAGAACGTGGCTAAGTTACTTAAGCTTCTGGAAGATTACGCCCACAAG<br>961 GATTACGAGAGTACTATGATCAAGGCGATTTTGTCTAGTATTGAACCGACAGTAGAGCCAAGCGAAGAGGAAGTCACTCG<br>1041 TTTAACCGACTATCTGTTTTCAGTGGAGCAATGGGGTTATTATGAAATTATCTTGCTTGGCAATTGTTCTCGATTCATTA<br>1121 ATTACAACACCCTGTTCTTACTGACAAAGGAGATGGTGACGAGTTTTGCATACAGTGAACAAAACAAGACGAACAAGACG<br>1201 CTGGTTACTCAACTCAGTATAAACTGTCTGATTATATCGATCGACTACAGTTACTTTGATCACTCCCACTACTTGATTGA<br>1281 AAAGATCGAATTCCTGCTGCGTGACGAGCTGAACTTTTACGAAAAGACGGTATTCCTCTACGTGCACGGATACTACAAGT<br>1361 TGAAACAGGGTCAAGTTTCGGGTAAGGACGATATGCGTCAGGCCTTACAAATTTTCAAGTACCTGGGCGAAGATGCGCTG<br>1441 TATTACTCCTATAAAGAGCACTACAGAAAGGAAGTGCTCGGTGACGAAGAGGATGAGGAAGAAGGCTGAATCCGGTAGCG<br>1521 GCGGTGGCAGCGGTGGCAGCGCGTGGAGCCATCCGCAGTTTGAAAAATAAGTCGACCGGCTGCTAACAAAGCCCGAAAGG<br>1601 AAGCTGAGTTGGCTGCTGCCACCGCTGAGCAATAACTAGCATAACCCCTTGGGGCCTCTAAACGGGTCTTGAGGGGTTTT<br>1681 TTGCTGAAAGCCAATTCTGATTAGAAAAACTCATCGAGCATCAAATGAAACTGCAATTTATTCATATCAGGATTATCAAT<br>1761 ACCATATTTTTGAAAAAGCCGTTTCTGTAATGAAGGAGAAAACTCACCGAGGCAGTTCCATAGGATGGCAAGATCCTGGT<br>1841 ATCGGTCTGCGATTCCGACTCGTCCAACATCAATACAACCTATTAATTTCCCCTCGTCAAAAATAAGGTTATCAAGTGAG<br>1921 AAATCACCATGAGTGACGACTGAATCCGGTGAGAATGGCAAAAGCTTATGCATTTCTTTCCAGACTTGTTCAACAGGCCA<br>2001 GCCATTACGCTCGTCATCAAAATCACTCGCATCAACCAAACCGTTATTCATTCGTGATTGCGCCTGAGCGAGACGAAATA |

| # | Type | Name and annotated features | Sequence (5′→3′) |
|---|---|---|---|
| | | | 2081 CGCGATCGCTGTTAAAAGGACAATTACAAACAGGAATCGAATGCAACCGGCGCAGGAACACTGCCAGCGCATCAACAATA<br>2161 TTTTCACCTGAATCAGGATATTCTTCTAATACCTGGAATGCTGTTTTCCCGGGGATCGCAGTGGTGAGTAACCATGCATC<br>2241 ATCAGGAGTACGGATAAAATGCTTGATGGTCGGAAGAGGCATAAATTCCGTCAGCCAGTTTAGTCTGACCATCTCATCTG<br>2321 TAACATCATTGGCAACGCTACCTTTGCCATGTTTCAGAAACAACTCTGGCGCATCGGGCTTCCCATACAATCGATAGATT<br>2401 GTCGCACCTGATTGCCCGACATTATCGCGAGCCCATTTATACCCATATAAATCAGCATCCATGTTGGAATTTAATCGCGG<br>2481 CTTCGAGCAAGACGTTTCCCGTTGAATATGGCTCATAACACCCCTTGTATTACTGTTTATGTAAGCAGACAGTTTTATTG<br>2561 TTCATGATGATATATTTTTATCTTGTGCAATGTAACATCAGAGATTTTGAGACACAACGTGAGATCAAAGGATCTTCTTG<br>2641 AGATCCTTTTTTTCTGCGCGTAATCTGCTGCTTGCAAAC |
| 2 | Plasmid | **pT7-Rgg3**<br>*Graham_Hendrickson_B6P_2_Rgg3_pLann · 2679 bp · circular*<br>Fig. 1–5; Suppl. Figs 2–4, 10–12<br><br>**Annotated features**<br>T7 promoter 575–593<br>RBS 625–647<br>Rgg3 CDS 655–1509<br>SET1 (fragment) 1579–1628<br>T7 terminator 1636–1683<br>KanR 1701–2516<br>ori 2638–2679, 1–547 | 1 AAAAAAACCACCGCTACCAGCGGTGGTTTGTTTGCCGGATCAAGAGCTACCAACTCTTTTTCCGAAGGTAACTGGCTTCA<br>81 GCAGAGCGCAGATACCAAATACTGTTCTTCTAGTGTAGCCGTAGTTAGGCCACCACTTCAAGAACTCTGTAGCACCGCCT<br>161 ACATACCTCGCTCTGCTAATCCTGTTACCAGTGGCTGCTGCCAGTGGCGATAAGTCGTGTCTTACCGGGTTGGACTCAAG<br>241 ACGATAGTTATCGGATAAGGCGCAGCGGTCGGGCTGAACGGGGGGTTCGTGCACACAGCCCAGCTTGGAGCGAACGACCT<br>321 ACACCGAACTGAGATACCTACAGCGTGAGCTATGAGAAAGCGCCACGCTTCCCGAAGGGAGAAAGGCGGACAGGTATCCG<br>401 GTAAGCGGCAGGGTCGGAACAGGAGAGCGCACGAGGGAGCTTCCAGGGGGAAACGCCTGGTATCTTTATAGTCCTGTCGG<br>481 GTTTCGCCACCTCTGACTTGAGCGTCGATTTTTGTGATGCTCGTCAGGGGGGCGGAGCCTATGGAAAAACGCCAGCAACG<br>561 CGATCCCGCGAAATTAATACGACTCACTATAGGGTGTCCACAACGGTTTCCCTCTAGAAATAATTTTGTTTAACTTTAAG<br>641 AAGGAGATATACATATGAAGTCTAAACTGGGGTCGACGCTTCGCAAAGTGAGAAACGGCAAGCAGATCTCTATCTGTTCC<br>721 GTTGCAGACGAGCACCTCTCTAAGTCCCAAATATCGAGATTTGAACGTGGAGAGAGCGAAATTAGCTGCATTCGGTTAAT<br>801 CAACATACTTGACAAACTTCATATAACGCTTGACGAATTCCTCATATTGCACGACGAGGATTACACAAAGACGGAAAGTT<br>881 TTGCCAACCTGGTGCAATACATCCGTAAGCAGTACTCACTGCAGAACATAAACAATATTCAATCATTATTATCCGATTCG<br>961 TCTAACTACACATTAGACCCGTTCGAAAAGACAATGGTGAAGTCCATCCTGCATACCATGGACTCGTCCATTATTCCATC<br>1041 AGACGACGAGTTGCTCCAGTTGGCAGATTATCTGTTCAAGGTGGAGAAGTGGGGCTATTATGAAATTATTCTGTTGGGCA<br>1121 ACTGCGTTCGGACTATTGATTACAACAGCGTGTTCTTGCTGACCAAGGAGATGTTAAACAACTATATTTATTCCTCTCTG<br>1201 AACAAAACCAACAAGCGGATAGTTACACAGCTGGCTATCAACTGCTTAATTTTGAGCATAGATATGGAAGAGTTCACTAA<br>1281 CTGTTTCTACCTGATAGACGAAATCAAAGCCTTACTTGACAACGAGCTTAACTTCTACGAGCAAACGGTATTCCTCTATG<br>1361 CCACCGGTTACTTTGAGTTCAAGCGGTGGCAATCGACTTCGGGCATTGAGAAGATGAAGCAAGCAATCCAAGTGTTAGAC<br>1441 ATATTGGGCGAAGATAACCTTAAGCTGCACTACACAATTCACTTTGACAAACTTATTAATAACAAGTGAATCCGGTAGCG<br>1521 GCGGTGGCAGCGGTGGCAGCGCGTGGAGCCATCCGCAGTTTGAAAAATAAGTCGACCGGCTGCTAACAAAGCCCGAAAGG<br>1601 AAGCTGAGTTGGCTGCTGCCACCGCTGAGCAATAACTAGCATAACCCCTTGGGGCCTCTAAACGGGTCTTGAGGGGTTTT<br>1681 TTGCTGAAAGCCAATTCTGATTAGAAAAACTCATCGAGCATCAAATGAAACTGCAATTTATTCATATCAGGATTATCAAT<br>1761 ACCATATTTTTGAAAAAGCCGTTTCTGTAATGAAGGAGAAAACTCACCGAGGCAGTTCCATAGGATGGCAAGATCCTGGT<br>1841 ATCGGTCTGCGATTCCGACTCGTCCAACATCAATACAACCTATTAATTTCCCCTCGTCAAAAATAAGGTTATCAAGTGAG<br>1921 AAATCACCATGAGTGACGACTGAATCCGGTGAGAATGGCAAAAGCTTATGCATTTCTTTCCAGACTTGTTCAACAGGCCA<br>2001 GCCATTACGCTCGTCATCAAAATCACTCGCATCAACCAAACCGTTATTCATTCGTGATTGCGCCTGAGCGAGACGAAATA<br>2081 CGCGATCGCTGTTAAAAGGACAATTACAAACAGGAATCGAATGCAACCGGCGCAGGAACACTGCCAGCGCATCAACAATA<br>2161 TTTTCACCTGAATCAGGATATTCTTCTAATACCTGGAATGCTGTTTTCCCGGGGATCGCAGTGGTGAGTAACCATGCATC<br>2241 ATCAGGAGTACGGATAAAATGCTTGATGGTCGGAAGAGGCATAAATTCCGTCAGCCAGTTTAGTCTGACCATCTCATCTG<br>2321 TAACATCATTGGCAACGCTACCTTTGCCATGTTTCAGAAACAACTCTGGCGCATCGGGCTTCCCATACAATCGATAGATT<br>2401 GTCGCACCTGATTGCCCGACATTATCGCGAGCCCATTTATACCCATATAAATCAGCATCCATGTTGGAATTTAATCGCGG<br>2481 CTTCGAGCAAGACGTTTCCCGTTGAATATGGCTCATAACACCCCTTGTATTACTGTTTATGTAAGCAGACAGTTTTATTG<br>2561 TTCATGATGATATATTTTTATCTTGTGCAATGTAACATCAGAGATTTTGAGACACAACGTGAGATCAAAGGATCTTCTTG<br>2641 AGATCCTTTTTTTCTGCGCGTAATCTGCTGCTTGCAAAC |
| 3 | Plasmid | **pT7-ComR-ST**<br>*Graham_Hendrickson_B6P_3_ComR_St_pLann · 2724 bp · circular*<br>Fig. 1; Suppl. Figs 2–5<br><br>**Annotated features**<br>T7 promoter 575–593<br>RBS 625–647<br>ComR-ST CDS 655–1554<br>SET1 (fragment) 1624–1673<br>T7 terminator 1681–1728<br>KanR 1746–2561<br>ori 2683–2724, 1–547 | 1 AAAAAAACCACCGCTACCAGCGGTGGTTTGTTTGCCGGATCAAGAGCTACCAACTCTTTTTCCGAAGGTAACTGGCTTCA<br>81 GCAGAGCGCAGATACCAAATACTGTTCTTCTAGTGTAGCCGTAGTTAGGCCACCACTTCAAGAACTCTGTAGCACCGCCT<br>161 ACATACCTCGCTCTGCTAATCCTGTTACCAGTGGCTGCTGCCAGTGGCGATAAGTCGTGTCTTACCGGGTTGGACTCAAG<br>241 ACGATAGTTATCGGATAAGGCGCAGCGGTCGGGCTGAACGGGGGGTTCGTGCACACAGCCCAGCTTGGAGCGAACGACCT<br>321 ACACCGAACTGAGATACCTACAGCGTGAGCTATGAGAAAGCGCCACGCTTCCCGAAGGGAGAAAGGCGGACAGGTATCCG<br>401 GTAAGCGGCAGGGTCGGAACAGGAGAGCGCACGAGGGAGCTTCCAGGGGGAAACGCCTGGTATCTTTATAGTCCTGTCGG<br>481 GTTTCGCCACCTCTGACTTGAGCGTCGATTTTTGTGATGCTCGTCAGGGGGGCGGAGCCTATGGAAAAACGCCAGCAACG<br>561 CGATCCCGCGAAATTAATACGACTCACTATAGGGTGTCCACAACGGTTTCCCTCTAGAAATAATTTTGTTTAACTTTAAG<br>641 AAGGAGATATACATATGAATCTGAAAGACAGTATAGGGTTACGCATTAAGACTGAGCGGGAACGGCAGCAAATGTCCCGC<br>721 GAGGTCTTGTGCCTTGACGGGGCTGAGTTAACGGTCCGGCAGCTGATTCGCATAGAGAAAGGGGAGTCCCTGCCGTCACT<br>801 GGATAGACTGTCGTACATCGCGAAGAGATTGGGCAAGTCGATGACCGAATTATTGGACCAAGACAACATTACAATCCCAG<br>881 ACGAGTACTACGAGATGAAGAATCGCCTGATCAAATTTCCGACTTACCGTAACCCCGATCGGATTAAAAGCAAGTTGACT<br>961 CTTATAGAGGAAGTTTACGAGAAGTTCTTTGACATCCTTCCGGAAGAGGAGTTACTTACATTAGACATACTGGAGAACAT<br>1041 CCTGTCATTCACGAGTTGGGAAGAATCGCCCAAAGTCGAGGAAATATATGAGGACTTATTCGAACAAGTAAAGCGCAAGC<br>1121 GTAAATTCAGCACAAATGATCTGCTTGTCATCGATTACTATTTCTTCCACCTGTATGGCCGTAAGCAGTATGACAAGAAG<br>1201 TTGTTCGAGCGTATCATTAAAAGAGTTTTGAACCAAGAGATATGGACCGACGACGTCTACAATATCGTGTTGTTTAACGA<br>1281 TTTAATGGCCATCGCGGCGCTCAAGATATTCCACAATAGTTTTAGCGACTTTCTGACGGTTGTTGACAAGGCGTTGGCCG<br>1361 TGATTGAAAAGTCTCAACTGTATTCCTATAAGCCTTCTGTGTTCGTTTTAAAAGCTAAGTACGAACTGCTGCACAAGGAA<br>1441 AACAAGAAGGAAGCCGCCGAAAACTATGACAAGGCTATAGTATTTGCGTCGGTGCTGGAAGACAGTGTTTTAGAGGAGTC<br>1521 TATTAAAGCGGGGAAGCTTGCAGACGGTTTATAGATCCGGTAGCGGCGGTGGCAGCGGTGGCAGCGCGTGGAGCCATCCG<br>1601 CAGTTTGAAAAATAAGTCGACCGGCTGCTAACAAAGCCCGAAAGGAAGCTGAGTTGGCTGCTGCCACCGCTGAGCAATAA<br>1681 CTAGCATAACCCCTTGGGGCCTCTAAACGGGTCTTGAGGGGTTTTTTGCTGAAAGCCAATTCTGATTAGAAAAACTCATC<br>1761 GAGCATCAAATGAAACTGCAATTTATTCATATCAGGATTATCAATACCATATTTTTGAAAAAGCCGTTTCTGTAATGAAG<br>1841 GAGAAAACTCACCGAGGCAGTTCCATAGGATGGCAAGATCCTGGTATCGGTCTGCGATTCCGACTCGTCCAACATCAATA<br>1921 CAACCTATTAATTTCCCCTCGTCAAAAATAAGGTTATCAAGTGAGAAATCACCATGAGTGACGACTGAATCCGGTGAGAA<br>2001 TGGCAAAAGCTTATGCATTTCTTTCCAGACTTGTTCAACAGGCCAGCCATTACGCTCGTCATCAAAATCACTCGCATCAA<br>2081 CCAAACCGTTATTCATTCGTGATTGCGCCTGAGCGAGACGAAATACGCGATCGCTGTTAAAAGGACAATTACAAACAGGA |

| # | Type | Name and annotated features | Sequence (5′→3′) |
|---|---|---|---|
| | | | 2161 ATCGAATGCAACCGGCGCAGGAACACTGCCAGCGCATCAACAATATTTTCACCTGAATCAGGATATTCTTCTAATACCTG<br>2241 GAATGCTGTTTTCCCGGGGATCGCAGTGGTGAGTAACCATGCATCATCAGGAGTACGGATAAAATGCTTGATGGTCGGAA<br>2321 GAGGCATAAATTCCGTCAGCCAGTTTAGTCTGACCATCTCATCTGTAACATCATTGGCAACGCTACCTTTGCCATGTTTC<br>2401 AGAAACAACTCTGGCGCATCGGGCTTCCCATACAATCGATAGATTGTCGCACCTGATTGCCCGACATTATCGCGAGCCCA<br>2481 TTTATACCCATATAAATCAGCATCCATGTTGGAATTTAATCGCGGCTTCGAGCAAGACGTTTCCCGTTGAATATGGCTCA<br>2561 TAACACCCCTTGTATTACTGTTTATGTAAGCAGACAGTTTTATTGTTCATGATGATATATTTTTATCTTGTGCAATGTAA<br>2641 CATCAGAGATTTTGAGACACAACGTGAGATCAAAGGATCTTCTTGAGATCCTTTTTTTCTGCGCGTAATCTGCTGCTTGC<br>2721 AAAC |
| 4 | Plasmid | **pT7-ComR-SI**<br>*Graham_Hendrickson_B6P_4_ComR_Si_pLann · 2733 bp · circular*<br>Fig. 1; Suppl. Figs 2–5<br><br>**Annotated features**<br>T7 promoter 575–593<br>RBS 625–647<br>ComR-SI CDS 655–1563<br>SET1 (fragment) 1633–1682<br>T7 terminator 1690–1737<br>KanR 1755–2570<br>ori 2692–2733, 1–547 | 1 AAAAAAACCACCGCTACCAGCGGTGGTTTGTTTGCCGGATCAAGAGCTACCAACTCTTTTTCCGAAGGTAACTGGCTTCA<br>81 GCAGAGCGCAGATACCAAATACTGTTCTTCTAGTGTAGCCGTAGTTAGGCCACCACTTCAAGAACTCTGTAGCACCGCCT<br>161 ACATACCTCGCTCTGCTAATCCTGTTACCAGTGGCTGCTGCCAGTGGCGATAAGTCGTGTCTTACCGGGTTGGACTCAAG<br>241 ACGATAGTTATCGGATAAGGCGCAGCGGTCGGGCTGAACGGGGGGTTCGTGCACACAGCCCAGCTTGGAGCGAACGACCT<br>321 ACACCGAACTGAGATACCTACAGCGTGAGCTATGAGAAAGCGCCACGCTTCCCGAAGGGAGAAAGGCGGACAGGTATCCG<br>401 GTAAGCGGCAGGGTCGGAACAGGAGAGCGCACGAGGGAGCTTCCAGGGGGAAACGCCTGGTATCTTTATAGTCCTGTCGG<br>481 GTTTCGCCACCTCTGACTTGAGCGTCGATTTTTGTGATGCTCGTCAGGGGGGCGGAGCCTATGGAAAAACGCCAGCAACG<br>561 CGATCCCGCGAAATTAATACGACTCACTATAGGGTGTCCACAACGGTTTCCCTCTAGAAATAATTTTGTTTAACTTTAAG<br>641 AAGGAGATATACATATGGAAAAGTTTGGAGAGAAAATTCGCCTGATGAGAGAAGAGAAGGGTATTTCTAGAGAAGAATTC<br>721 TGTGGGGACGAGACTGAACTGTCCGTTCGGCAGCTCGCACGAATCGAATTGAACCAGTCCATCCCAAACTTATCAAAAGC<br>801 CAGTTTTATAGCGAATCGGCTGGGCGTGAAGCTGGGGACGCTGACAGACGGGGACTCATTGGAGCTTCCTAAGCGTTACA<br>881 AGGAGCTGAAGTACCTTCTGCTTCGTACGCCAACCTACGGTGATCAAGTGAGACTGGATCGTAAGAACGACTATTTTGAT<br>961 GAAATCGCTGAAGTCTTCTACGATGTAATTCCAGAGGAAGAACGGCTGATAATAGACTGCCTTCAATCTAAGTTCGACGT<br>1041 ACATTTCTCCGAGGATGTGAATTTCGGCGAGGGTATCTTAAACGACTACTTCGGGCAAGTAAACCGGAAGAAAGTTTTCA<br>1121 CCATAAATGACTTAATCCTTATTGATTTGTATTTTGCATGCTTGAGTTCCGCCAAGAAGTTCGAGGGGATATACAGTCTG<br>1201 GACTTTTACGACGATCTTATGAAGAGATTAGTCAATCAAAAGCACGTTTCTCCCGAGACAGATCTTATCCTGAACAACGT<br>1281 TCTGTTAAATAACATAGACTTAGCATTCCAATTCAAACGAGAGTACTACATTGAACGTGTCATCACCATAAGCGAGAAGA<br>1361 TTATGACTGAGATCCACGACTTCCAGCGCCGCCCAATCCTGTCACTGGTCGAATGGAAGTATTACCTTAAGCTCAAGCAC<br>1441 GATTTTGCGCTGGCGGAACAAAGCTTCACAAATGCTACTCTTTTCGCACGACTGGTTGGAGACACATACTTGGAGAACAA<br>1521 GCTCAAGGAAGAATGGCAACTGGACATCGACGCGGTTGAGTGAATCCGGTAGCGGCGGTGGCAGCGGTGGCAGCGCGTGG<br>1601 AGCCATCCGCAGTTTGAAAAATAAGTCGACCGGCTGCTAACAAAGCCCGAAAGGAAGCTGAGTTGGCTGCTGCCACCGCT<br>1681 GAGCAATAACTAGCATAACCCCTTGGGGCCTCTAAACGGGTCTTGAGGGGTTTTTTGCTGAAAGCCAATTCTGATTAGAA<br>1761 AAACTCATCGAGCATCAAATGAAACTGCAATTTATTCATATCAGGATTATCAATACCATATTTTTGAAAAAGCCGTTTCT<br>1841 GTAATGAAGGAGAAAACTCACCGAGGCAGTTCCATAGGATGGCAAGATCCTGGTATCGGTCTGCGATTCCGACTCGTCCA<br>1921 ACATCAATACAACCTATTAATTTCCCCTCGTCAAAAATAAGGTTATCAAGTGAGAAATCACCATGAGTGACGACTGAATC<br>2001 CGGTGAGAATGGCAAAAGCTTATGCATTTCTTTCCAGACTTGTTCAACAGGCCAGCCATTACGCTCGTCATCAAAATCAC<br>2081 TCGCATCAACCAAACCGTTATTCATTCGTGATTGCGCCTGAGCGAGACGAAATACGCGATCGCTGTTAAAAGGACAATTA<br>2161 CAAACAGGAATCGAATGCAACCGGCGCAGGAACACTGCCAGCGCATCAACAATATTTTCACCTGAATCAGGATATTCTTC<br>2241 TAATACCTGGAATGCTGTTTTCCCGGGGATCGCAGTGGTGAGTAACCATGCATCATCAGGAGTACGGATAAAATGCTTGA<br>2321 TGGTCGGAAGAGGCATAAATTCCGTCAGCCAGTTTAGTCTGACCATCTCATCTGTAACATCATTGGCAACGCTACCTTTG<br>2401 CCATGTTTCAGAAACAACTCTGGCGCATCGGGCTTCCCATACAATCGATAGATTGTCGCACCTGATTGCCCGACATTATC<br>2481 GCGAGCCCATTTATACCCATATAAATCAGCATCCATGTTGGAATTTAATCGCGGCTTCGAGCAAGACGTTTCCCGTTGAA<br>2561 TATGGCTCATAACACCCCTTGTATTACTGTTTATGTAAGCAGACAGTTTTATTGTTCATGATGATATATTTTTATCTTGT<br>2641 GCAATGTAACATCAGAGATTTTGAGACACAACGTGAGATCAAAGGATCTTCTTGAGATCCTTTTTTTCTGCGCGTAATCT<br>2721 GCTGCTTGCAAAC |
| 5 | Plasmid | **pT7-PlcR**<br>*Graham_Hendrickson_B6P_6_PlcR_pLann · 2682 bp · circular*<br>Fig. 1; Suppl. Figs 2–4<br><br>**Annotated features**<br>T7 promoter 575–593<br>RBS 625–647<br>PlcR CDS 655–1512<br>SET1 (fragment) 1582–1631<br>T7 terminator 1639–1686<br>KanR 1704–2519<br>ori 2641–2682, 1–547 | 1 AAAAAAACCACCGCTACCAGCGGTGGTTTGTTTGCCGGATCAAGAGCTACCAACTCTTTTTCCGAAGGTAACTGGCTTCA<br>81 GCAGAGCGCAGATACCAAATACTGTTCTTCTAGTGTAGCCGTAGTTAGGCCACCACTTCAAGAACTCTGTAGCACCGCCT<br>161 ACATACCTCGCTCTGCTAATCCTGTTACCAGTGGCTGCTGCCAGTGGCGATAAGTCGTGTCTTACCGGGTTGGACTCAAG<br>241 ACGATAGTTATCGGATAAGGCGCAGCGGTCGGGCTGAACGGGGGGTTCGTGCACACAGCCCAGCTTGGAGCGAACGACCT<br>321 ACACCGAACTGAGATACCTACAGCGTGAGCTATGAGAAAGCGCCACGCTTCCCGAAGGGAGAAAGGCGGACAGGTATCCG<br>401 GTAAGCGGCAGGGTCGGAACAGGAGAGCGCACGAGGGAGCTTCCAGGGGGAAACGCCTGGTATCTTTATAGTCCTGTCGG<br>481 GTTTCGCCACCTCTGACTTGAGCGTCGATTTTTGTGATGCTCGTCAGGGGGGCGGAGCCTATGGAAAAACGCCAGCAACG<br>561 CGATCCCGCGAAATTAATACGACTCACTATAGGGTGTCCACAACGGTTTCCCTCTAGAAATAATTTTGTTTAACTTTAAG<br>641 AAGGAGATATACATATGCAAGCTGAGAAACTGGGATCCGAAATAAAGAAGATCCGTGTACTCAGAGGTCTTACGCAAAAG<br>721 CAACTTAGTGAGAATATATGTCATCAGTCGGAAGTGAGCCGTATCGAGAGTGGTGCCGTTTACCCATCCATGGACATTTT<br>801 ACAAGGTATCGCTGCGAAACTGCAAATTCCGATCATACATTTCTACGAGGTGCTTATCTACAGCGACATTGAGCGCAAGA<br>881 AGCAATTTAAGGACCAGGTCATAATGCTGTGCAAGCAAAAGCGTTATAAAGAGATATACAATAAGGTTTGGAACGAGCTT<br>961 AAGAAAGAGGAGTATCATCCTGAGTTCCAACAATTCTTGCAATGGCAGTACTACGTTGCGGCTTACGTTTTAAAGAAGGT<br>1041 GGACTATGAATACTGCATACTGGAATTAAAGAAGCTTTTAAACCAACAACTTACTGGCATAGACGTTTACCAAAACCTGT<br>1121 ACATCGAGAACGCAATAGCAAATATCTACGCCGAAAACGGATACCTGAAGAAGGGTATCGATCTCTTCGAGCAGATCCTG<br>1201 AAGCAATTAGAAGCCCTGCACGATAACGAGGAGTTCGACGTGAAGGTGCGCTACAATCACGCAAAGGCACTGTACCTGGA<br>1281 CAGCCGTTACGAAGAGTCTTTATACCAAGTAAACAAGGCGATCGAAATTAGCTGCAGAATAAACAGTATGGCGCTGATTG<br>1361 GCCAACTGTACTATCAACGGGGTGAGTGCCTTCGCAAGCTCGAGTATGAGGAAGCCGAAATAGAGGACGCGTACAAGAAG<br>1441 GCAAGCTTCTTCTTCGACATATTAGAGATGCACGCGTATAAGGAAGCGTTGGTAAACAAGATTAGTAGATAAATCCGGTA<br>1521 GCGGCGGTGGCAGCGGTGGCAGCGCGTGGAGCCATCCGCAGTTTGAAAAATAAGTCGACCGGCTGCTAACAAAGCCCGAA<br>1601 AGGAAGCTGAGTTGGCTGCTGCCACCGCTGAGCAATAACTAGCATAACCCCTTGGGGCCTCTAAACGGGTCTTGAGGGGT<br>1681 TTTTTGCTGAAAGCCAATTCTGATTAGAAAAACTCATCGAGCATCAAATGAAACTGCAATTTATTCATATCAGGATTATC<br>1761 AATACCATATTTTTGAAAAAGCCGTTTCTGTAATGAAGGAGAAAACTCACCGAGGCAGTTCCATAGGATGGCAAGATCCT<br>1841 GGTATCGGTCTGCGATTCCGACTCGTCCAACATCAATACAACCTATTAATTTCCCCTCGTCAAAAATAAGGTTATCAAGT<br>1921 GAGAAATCACCATGAGTGACGACTGAATCCGGTGAGAATGGCAAAAGCTTATGCATTTCTTTCCAGACTTGTTCAACAGG<br>2001 CCAGCCATTACGCTCGTCATCAAAATCACTCGCATCAACCAAACCGTTATTCATTCGTGATTGCGCCTGAGCGAGACGAA |

| # | Type | Name and annotated features | Sequence (5′→3′) |
|---|---|---|---|
| | | | 2081 ATACGCGATCGCTGTTAAAAGGACAATTACAAACAGGAATCGAATGCAACCGGCGCAGGAACACTGCCAGCGCATCAACA<br>2161 ATATTTTCACCTGAATCAGGATATTCTTCTAATACCTGGAATGCTGTTTTCCCGGGGATCGCAGTGGTGAGTAACCATGC<br>2241 ATCATCAGGAGTACGGATAAAATGCTTGATGGTCGGAAGAGGCATAAATTCCGTCAGCCAGTTTAGTCTGACCATCTCAT<br>2321 CTGTAACATCATTGGCAACGCTACCTTTGCCATGTTTCAGAAACAACTCTGGCGCATCGGGCTTCCCATACAATCGATAG<br>2401 ATTGTCGCACCTGATTGCCCGACATTATCGCGAGCCCATTTATACCCATATAAATCAGCATCCATGTTGGAATTTAATCG<br>2481 CGGCTTCGAGCAAGACGTTTCCCGTTGAATATGGCTCATAACACCCCTTGTATTACTGTTTATGTAAGCAGACAGTTTTA<br>2561 TTGTTCATGATGATATATTTTTATCTTGTGCAATGTAACATCAGAGATTTTGAGACACAACGTGAGATCAAAGGATCTTC<br>2641 TTGAGATCCTTTTTTTCTGCGCGTAATCTGCTGCTTGCAAAC |
| **B. Reporter plasmids (pRRNPP-sfGFP; template for reporter LETs)** | | | |
| 6 | Plasmid | **pRgg2-sfGFP**<br>*Graham_Hendrickson_Q6Y_1_Rgg2_prom_WT_pLann · 3983 bp · circular*<br>Fig. 1, 2; Suppl. Figs 2–4, 8–9, 11–12<br><br>**Annotated features**<br>AmpR 213–1073<br>AmpR promoter 1074–1178<br>rop (fragment) 1273–1308<br>bom (fragment) 1410–1509<br>lacZα (fragment) 1515–1668<br>Rgg2 promoter 1679–1787<br>sfGFP 1870–2583<br>5S rRNA 2777–2892<br>rrnB T1 terminator 2894–2940<br>MCS (fragment) 2989–3023<br>lac operator 3060–3076<br>lac promoter 3084–3114<br>CAP binding site 3129–3150<br>lacI (fragment) 3163–3255<br>ori 3437–3983, 1–42 | 1 GTTTGCAAGCAGCAGATTACGCGCAGAAAAAAAGGATCTCAAGAAGATCCTTTGATCTTTTCTACGGGGTCTGACGCTCA<br>81 GTGGAACGAAAACTCACGTTAAGGGATTTTGGTCATGAGATTATCAAAAAGGATCTTCACCTAGATCCTTTTAAATTAAA<br>161 AATGAAGTTTTAAATCAATCTAAAGTATATATGAGTAAACTTGGTCTGACAGTTACCAATGCTTAATCAGTGAGGCACCT<br>241 ATCTCAGCGATCTGTCTATTTCGTTCATCCATAGTTGCCTGACTCCCCGTCGTGTAGATAACTACGATACGGGAGGGCTT<br>321 ACCATCTGGCCCCAGTGCTGCAATGATACCGCGACTCCCACGCTCACCGGCTCCAGATTTATCAGCAATAAACCAGCCAG<br>401 CCGGAAGGGCCGAGCGCAGAAGTGGTCCTGCAACTTTATCCGCCTCCATCCAGTCTATTAATTGTTGCCGGGAAGCTAGA<br>481 GTAAGTAGTTCGCCAGTTAATAGTTTGCGCAACGTTGTTGCCATTGCTACAGGCATCGTGGTGTCACGCTCGTCGTTTGG<br>561 TATGGCTTCATTCAGCTCCGGTTCCCAACGATCAAGGCGAGTTACATGATCCCCCATGTTGTGCAAAAAAGCGGTTAGCT<br>641 CCTTCGGTCCTCCGATCGTTGTCAGAAGTAAGTTGGCCGCAGTGTTATCACTCATGGTTATGGCAGCACTGCATAATTCT<br>721 CTTACTGTCATGCCATCCGTAAGATGCTTTTCTGTGACTGGTGAGTACTCAACCAAGTCATTCTGAGAATAGTGTATGCG<br>801 GCGACCGAGTTGCTCTTGCCCGGCGTCAATACGGGATAATACCGCGCCACATAGCAGAACTTTAAAAGTGCTCATCATTG<br>881 GAAAACGTTCTTCGGGGCGAAAACTCTCAAGGATCTTACCGCTGTTGAGATCCAGTTCGATGTAACCCACTCGTGCACCC<br>961 AACTGATCTTCAGCATCTTTTACTTTCACCAGCGTTTCTGGGTGAGCAAAAACAGGAAGGCAAAATGCCGCAAAAAAGGG<br>1041 AATAAGGGCGACACGGAAATGTTGAATACTCATACTCTTCCTTTTTCAATATTATTGAAGCATTTATCAGGGTTATTGTC<br>1121 TCATGAGCGGATACATATTTGAATGTATTTAGAAAAATAAACAAATAGGGGTTCCGCGCACATTTCCCCGAAAAGTGCCA<br>1201 CCTGACGTCTAAGAAACCATTATTATCATGACATTAACCTATAAAAATAGGCGTATCACGAGGCCCTTTCGTCTCGCGCG<br>1281 TTTCGGTGATGACGGTGAAAACCTCTGACACATGCAGCTCCCGGAGACGGTCACAGCTTGTCTGTAAGCGGATGCCGGGA<br>1361 GCAGACAAGCCCGTCAGGGCGCGTCAGCGGGTGTTGGCGGGTGTCGGGGCTGGCTTAACTATGCGGCATCAGAGCAGATT<br>1441 GTACTGAGAGTGCACCATATGCGGTGTGAAATACCGCACAGATGCGTAAGGAGAAAATACCGCATCAGGCGCCATTCGCC<br>1521 ATTCAGGCTGCGCAACTGTTGGGAAGGGCGATCGGTGCGGGCCTCTTCGCTATTACGCCAGCTGGCGAAAGGGGGATGTG<br>1601 CTGCAAGGCGATTAAGTTGGGTAACGCCAGGGTTTTCCCAGTCACGACGTTGTAAAACGACGGCCAGTAGATCTTATGTG<br>1681 TTTTCCTTGGCGTAGTCGCCTTAAGGTTTACCGAGTTCTTTTCCATTTTCCCACTTTCACAACAAAAAATTCAAATC<br>1761 CGTCAAATCCATTATATAATACAAGTAACGTCGACTCTCGAGTGAGATTGTTGACGGTACCGTATTTTGGATCTGAGCAA<br>1841 AGCAATAACAGAACATAAGGAGGTACGTAATGAGCAAAGGAGAAGAACTTTTCACTGGAGTTGTCCCAATTCTTGTTGAA<br>1921 TTAGATGGTGATGTTAATGGGCACAAATTTTCTGTCCGTGGAGAGGGTGAAGGTGATGCTACAAACGGAAAACTCACCCT<br>2001 TAAATTTATTTGCACTACTGGAAAACTACCTGTTCCGTGGCCAACACTTGTCACTACTCTGACCTATGGTGTTCAATGCT<br>2081 TTTCCCGTTATCCGGATCACATGAAACGGCATGACTTTTTCAAGAGTGCAATGCCCGAAGGTTATGTACAGGAACGCACT<br>2161 ATATCTTTCAAAGATGACGGGACCTACAAGACGCGTGCTGAAGTCAAGTTTGAAGGTGATACCCTTGTTAATCGTATCGA<br>2241 GTTAAAGGGTATTGATTTTAAAGAAGATGGAAACATTCTTGGACACAAACTCGAGTACAACTTTAACTCACACAATGTAT<br>2321 ACATCACGGCAGACAAACAAAAGAATGGAATCAAAGCTAACTTCAAAATTCGCCACAACGTTGAAGATGGTTCCGTTCAA<br>2401 CTAGCAGACCATTATCAACAAAATACTCCAATTGGCGATGGCCCTGTCCTTTTACCAGACAACCATTACCTGTCGACACA<br>2481 ATCTGTCCTTTCGAAAGATCCCAACGAAAAGCGTGACCACATGGTCCTTCTTGAGTTTGTAACTGCTGCTGGGATTACAC<br>2561 ATGGCATGGATGAGCTCTACAAATAAGGATCTGAAGCTTGGGCCCGAACAAAAACTCATCTCAGAAGAGGATCTGAATAG<br>2641 CGCCGTCGACCATCATCATCATCATCATTGAGTTTAAACGGACACCAGCTTGGCTGTTTTGGCGGATGAGAGAAGATTTT<br>2721 CAGCCTGATACAGATTAAATCAGAACGCAGAAGCGGTCTGATAAAACAGAATTTGCCTGGCGGCAGTAGCGCGGTGGTCC<br>2801 CACCTGACCCCATGCCGAACTCAGAAGTGAAACGCCGTAGCGCCGATGGTAGTGTGGGGACACCCCATGCGAGAGTAGGG<br>2881 AACTGCCAGGCATCAAATAAAACGAAAGGCTCAGTCGAAAGACTGGGCCTTTCGTTTTATCTGTTGTTTGTCGGTGAACT<br>2961 CTGAAAGGAGGAACTATATCCGGATCGAGATCCTCTAGAGTCGACCTGCAGGCATGCAAGCTTGGCGTAATCATGGTCAT<br>3041 AGCTGTTTCCTGTGTGAAATTGTTATCCGCTCACAATTCCACACAACATACGAGCCGGAAGCATAAAGTGTAAAGCCTGG<br>3121 GGTGCCTAATGAGTGAGCTAACTCACATTAATTGCGTTGCGCTCACTGCCCGCTTTCCAGTCGGGAAACCTGTCGTGCCA<br>3201 GCTGCATTAATGAATCGGCCAACGCGCGGGGAGAGGCGGTTTGCGTATTGGGCGCTCTTCCGCTTCCTCGCTCACTGACT<br>3281 CGCTGCGCTCGGTCGTTCGGCTGCGGCGAGCGGTATCAGCTCACTCAAAGGCGGTAATACGGTTATCCACAGAATCAGGG<br>3361 GATAACGCAGGAAAGAACATGTGAGCAAAAGGCCAGCAAAAGGCCAGGAACCGTAAAAAGGCCGCGTTGCTGGCGTTTTTC<br>3441 CATAGGCTCCGCCCCCCTGACGAGCATCACAAAAATCGACGCTCAAGTCAGAGGTGGCGAAACCCGACAGGACTATAAAG<br>3521 ATACCAGGCGTTTCCCCCTGGAAGCTCCCTCGTGCGCTCTCCTGTTCCGACCCTGCCGCTTACCGGATACCTGTCCGCCT<br>3601 TTCTCCCTTCGGGAAGCGTGGCGCTTTCTCATAGCTCACGCTGTAGGTATCTCAGTTCGGTGTAGGTCGTTCGCTCCAAG<br>3681 CTGGGCTGTGTGCACGAACCCCCCGTTCAGCCCGACCGCTGCGCCTTATCCGGTAACTATCGTCTTGAGTCCAACCCGGT<br>3761 AAGACACGACTTATCGCCACTGGCAGCAGCCACTGGTAACAGGATTAGCAGAGCGAGGTATGTAGGCGGTGCTACAGAGT<br>3841 TCTTGAAGTGGTGGCCTAACTACGGCTACACTAGAAGAACAGTATTTGGTATCTGCGCTCTGCTGAAGCCAGTTACCTTC<br>3921 GGAAAAAGAGTTGGTAGCTCTTGATCCGGCAAACAAACCACCGCTGGTAGCGGTGGTTTTTTT |
| 7 | Plasmid | **pRgg3-sfGFP**<br>*Graham_Hendrickson_Q6Y_2_Rgg3_prom_WT_pLann · 3978 bp · circular*<br>Fig. 1–5; Suppl. Figs 2–4, 8–12<br><br>**Annotated features**<br>lacZα (fragment) 102–255 | 1 AAAAAAAAACGGAATTGATTTTTGTTGGGAAAGTGGGAAAATATGAAATCAAAACTTGGTGTCACACTTAGAAAAGTTCG<br>81 AAAAGGAAAGCCATAAGATCTACTGGCCGTCGTTTTACAACGTCGTGACTGGGAAAACCCTGGCGTTACCCAACTTAATC<br>161 GCCTTGCAGCACATCCCCCTTTCGCCAGCTGGCGTAATAGCGAAGAGGCCCGCACCGATCGCCCTTCCCAACAGTTGCGC<br>241 AGCCTGAATGGCGAATGGCGCCTGATGCGGTATTTTCTCCTTACGCATCTGTGCGGTATTTCACACCGCATATGGTGCAC<br>321 TCTCAGTACAATCTGCTCTGATGCCGCATAGTTAAGCCAGCCCCGACACCCGCCAACACCCGCTGACGCGCCCTGACGGG<br>401 CTTGTCTGCTCCCGGCATCCGCTTACAGACAAGCTGTGACCGTCTCCGGGAGCTGCATGTGTCAGAGGTTTTCACCGTCA<br>481 TCACCGAAACGCGCGAGACGAAAGGGCCTCGTGATACGCCTATTTTTATAGGTTAATGTCATGATAATAATGGTTTCTTA<br>561 GACGTCAGGTGGCACTTTTCGGGGAAATGTGCGCGGAACCCCTATTTGTTTATTTTTCTAAATACATTCAAATATGTATC<br>641 CGCTCATGAGACAATAACCCTGATAAATGCTTCAATAATATTGAAAAAGGAAGAGTATGAGTATTCAACATTTCCGTGTC |

| # | Type | Name and annotated features | Sequence (5′→3′) |
|---|---|---|---|
| | | bom (fragment) 261–360<br>rop (fragment) 462–497<br>AmpR promoter 592–696<br>AmpR 697–1557<br>ori 1728–2316<br>lacI (fragment) 2498–2590<br>CAP binding site 2603–2624<br>lac promoter 2639–2669<br>lac operator 2677–2693<br>MCS (fragment) 2730–2764<br>rrnB T1 terminator 2813–2859<br>5S rRNA 2861–2976<br>sfGFP 3170–3883<br>Rgg3 promoter 3963–3978, 1–95 (rev) | 721 GCCCTTATTCCCTTTTTTGCGGCATTTTGCCTTCCTGTTTTTGCTCACCCAGAAACGCTGGTGAAAGTAAAAGATGCTGA<br>801 AGATCAGTTGGGTGCACGAGTGGGTTACATCGAACTGGATCTCAACAGCGGTAAGATCCTTGAGAGTTTTCGCCCCGAAG<br>881 AACGTTTTCCAATGATGAGCACTTTTAAAGTTCTGCTATGTGGCGCGGTATTATCCCGTATTGACGCCGGGCAAGAGCAA<br>961 CTCGGTCGCCGCATACACTATTCTCAGAATGACTTGGTTGAGTACTCACCAGTCACAGAAAAGCATCTTACGGATGGCAT<br>1041 GACAGTAAGAGAATTATGCAGTGCTGCCATAACCATGAGTGATAACACTGCGGCCAACTTACTTCTGACAACGATCGGAG<br>1121 GACCGAAGGAGCTAACCGCTTTTTTGCACAACATGGGGGATCATGTAACTCGCCTTGATCGTTGGGAACCGGAGCTGAAT<br>1201 GAAGCCATACCAAACGACGAGCGTGACACCACGATGCCTGTAGCAATGGCAACAACGTTGCGCAAACTATTAACTGGCGA<br>1281 ACTACTTACTCTAGCTTCCCGGCAACAATTAATAGACTGGATGGAGGCGGATAAAGTTGCAGGACCACTTCTGCGCTCGG<br>1361 CCCTTCCGGCTGGCTGGTTTATTGCTGATAAATCTGGAGCCGGTGAGCGTGGGAGTCGCGGTATCATTGCAGCACTGGGG<br>1441 CCAGATGGTAAGCCCTCCCGTATCGTAGTTATCTACACGACGGGGAGTCAGGCAACTATGGATGAACGAAATAGACAGAT<br>1521 CGCTGAGATAGGTGCCTCACTGATTAAGCATTGGTAACTGTCAGACCAAGTTTACTCATATATACTTTAGATTGATTTAA<br>1601 AACTTCATTTTTAATTTAAAAGGATCTAGGTGAAGATCCTTTTTGATAATCTCATGACCAAAATCCCTTAACGTGAGTTT<br>1681 TCGTTCCACTGAGCGTCAGACCCCGTAGAAAAGATCAAAGGATCTTCTTGAGATCCTTTTTTTCTGCGCGTAATCTGCTG<br>1761 CTTGCAAACAAAAAAACCACCGCTACCAGCGGTGGTTTGTTTGCCGGATCAAGAGCTACCAACTCTTTTTCCGAAGGTAA<br>1841 CTGGCTTCAGCAGAGCGCAGATACCAAATACTGTTCTTCTAGTGTAGCCGTAGTTAGGCCACCACTTCAAGAACTCTGTA<br>1921 GCACCGCCTACATACCTCGCTCTGCTAATCCTGTTACCAGTGGCTGCTGCCAGTGGCGATAAGTCGTGTCTTACCGGGTT<br>2001 GGACTCAAGACGATAGTTACCGGATAAGGCGCAGCGGTCGGGCTGAACGGGGGGTTCGTGCACACAGCCCAGCTTGGAGC<br>2081 GAACGACCTACACCGAACTGAGATACCTACAGCGTGAGCTATGAGAAAGCGCCACGCTTCCCGAAGGGAGAAAGGCGGAC<br>2161 AGGTATCCGGTAAGCGGCAGGGTCGGAACAGGAGAGCGCACGAGGGAGCTTCCAGGGGGAAACGCCTGGTATCTTTATAG<br>2241 TCCTGTCGGGTTTCGCCACCTCTGACTTGAGCGTCGATTTTTGTGATGCTCGTCAGGGGGGCGGAGCCTATGGAAAAACG<br>2321 CCAGCAACGCGGCCTTTTTACGGTTCCTGGCCTTTTGCTGGCCTTTTGCTCACATGTTCTTTCCTGCGTTATCCCCTGATT<br>2401 CTGTGGATAACCGTATTACCGCCTTTGAGTGAGCTGATACCGCTCGCCGCAGCCGAACGACCGAGCGCAGCGAGTCAGTG<br>2481 AGCGAGGAAGCGGAAGAGCGCCCAATACGCAAACCGCCTCTCCCCGCGCGTTGGCCGATTCATTAATGCAGCTGGCACGA<br>2561 CAGGTTTCCCGACTGGAAAGCGGGCAGTGAGCGCAACGCAATTAATGTGAGTTAGCTCACTCATTAGGCACCCCAGGCTT<br>2641 TACACTTTATGCTTCCGGCTCGTATGTTGTGTGGAATTGTGAGCGGATAACAATTTCACACAGGAAACAGCTATGACCAT<br>2721 GATTACGCCAAGCTTGCATGCCTGCAGGTCGACTCTAGAGGATCTCGATCCGGATATAGTTCCTCCTTTCAGAGTTCACC<br>2801 GACAAACAACAGATAAAACGAAAGGCCCAGTCTTTCGACTGAGCCTTTCGTTTTATTTGATGCCTGGCAGTTCCCTACTC<br>2881 TCGCATGGGGTGTCCCCACACTACCATCGGCGCTACGGCGTTTCACTTCTGAGTTCGGCATGGGGTCAGGTGGGACCACC<br>2961 GCGCTACTGCCGCCAGGCAAATTCTGTTTTATCAGACCGCTTCTGCGTTCTGATTTAATCTGTATCAGGCTGAAAATCTT<br>3041 CTCTCATCCGCCAAAACAGCCAAGCTGGTGTCCGTTTAAACTCAATGATGATGATGATGATGGTCGACGGCGCTATTCAG<br>3121 ATCCTCTTCTGAGATGAGTTTTTGTTCGGGCCCAAGCTTCAGATCCTTATTTGTAGAGCTCATCCATGCCATGTGTAATC<br>3201 CCAGCAGCAGTTACAAACTCAAGAAGGACCATGTGGTCACGCTTTTCGTTGGGATCTTTCGAAAGGACAGATTGTGTCGA<br>3281 CAGGTAATGGTTGTCTGGTAAAAGGACAGGGCCATCGCCAATTGGAGTATTTTGTTGATAATGGTCTGCTAGTTGAACGG<br>3361 AACCATCTTCAACGTTGTGGCGAATTTTGAAGTTAGCTTTGATTCCATTCTTTTGTTTGTCTGCCGTGATGTATACATTG<br>3441 TGTGAGTTAAAGTTGTACTCGAGTTTGTGTCCAAGAATGTTTCCATCTTCTTTAAAATCAATACCCTTTAACTCGATACG<br>3521 ATTAACAAGGGTATCACCTTCAAACTTGACTTCAGCACGCGTCTTGTAGGTCCCGTCATCTTTGAAAGATATAGTGCGTT<br>3601 CCTGTACATAACCTTCGGGCATTGCACTCTTGAAAAAGTCATGCCGTTTCATGTGATCCGGATAACGGGAAAAGCATTGA<br>3681 ACACCATAGGTCAGAGTAGTGACAAGTGTTGGCCACGGAACAGGTAGTTTTCCAGTAGTGCAAATAAATTTAAGGGTGAG<br>3761 TTTTCCGTTTGTAGCATCACCTTCACCCTCTCCACGGACAGAAAATTTGTGCCCATTAACATCACCATCTAATTCAACAA<br>3841 GAATTGGGACAACTCCAGTGAAAAGTTCTTCTCCTTTGCTCATTACGTACCTCCTTATGTTCTGTTATTGCTTTGCTCAG<br>3921 ATCCAAAATACGGTACCGTCAACAATCTCACTCGAGAGTCGACGTTTTATTGTACAAT |
| 8 | Plasmid | **pComR-ST-sfGFP**<br>*Graham_Hendrickson_Q6Y_3_ComR_St_prom_WT_pLann · 3978 bp · circular*<br>Fig. 1; Suppl. Figs 2–5, 9<br><br>**Annotated features**<br>AmpR 213–1073<br>AmpR promoter 1074–1178<br>rop (fragment) 1273–1308<br>bom (fragment) 1410–1509<br>lacZα (fragment) 1515–1668<br>ComR-ST promoter 1679–1782<br>sfGFP 1865–2578<br>5S rRNA 2772–2887<br>rrnB T1 terminator 2889–2935<br>MCS (fragment) 2984–3018<br>lac operator 3055–3071<br>lac promoter 3079–3109<br>CAP binding site 3124–3145<br>lacI (fragment) 3158–3250<br>ori 3432–3978, 1–42 | 1 GTTTGCAAGCAGCAGATTACGCGCAGAAAAAAAGGATCTCAAGAAGATCCTTTGATCTTTTCTACGGGGTCTGACGCTCA<br>81 GTGGAACGAAAACTCACGTTAAGGGATTTTGGTCATGAGATTATCAAAAAGGATCTTCACCTAGATCCTTTTAAATTAAA<br>161 AATGAAGTTTTAAATCAATCTAAAGTATATATGAGTAAACTTGGTCTGACAGTTACCAATGCTTAATCAGTGAGGCACCT<br>241 ATCTCAGCGATCTGTCTATTTCGTTCATCCATAGTTGCCTGACTCCCCGTCGTGTAGATAACTACGATACGGGAGGGCTT<br>321 ACCATCTGGCCCCAGTGCTGCAATGATACCGCGACTCCCACGCTCACCGGCTCCAGATTTATCAGCAATAAACCAGCCAG<br>401 CCGGAAGGGCCGAGCGCAGAAGTGGTCCTGCAACTTTATCCGCCTCCATCCAGTCTATTAATTGTTGCCGGGAAGCTAGA<br>481 GTAAGTAGTTCGCCAGTTAATAGTTTGCGCAACGTTGTTGCCATTGCTACAGGCATCGTGGTGTCACGCTCGTCGTTTGG<br>561 TATGGCTTCATTCAGCTCCGGTTCCCAACGATCAAGGCGAGTTACATGATCCCCCATGTTGTGCAAAAAAGCGGTTAGCT<br>641 CCTTCGGTCCTCCGATCGTTGTCAGAAGTAAGTTGGCCGCAGTGTTATCACTCATGGTTATGGCAGCACTGCATAATTCT<br>721 CTTACTGTCATGCCATCCGTAAGATGCTTTTCTGTGACTGGTGAGTACTCAACCAAGTCATTCTGAGAATAGTGTATGCG<br>801 GCGACCGAGTTGCTCTTGCCCGGCGTCAATACGGGATAATACCGCGCCACATAGCAGAACTTTAAAAGTGCTCATCATTG<br>881 GAAAACGTTCTTCGGGGCGAAAACTCTCAAGGATCTTACCGCTGTTGAGATCCAGTTCGATGTAACCCACTCGTGCACCC<br>961 AACTGATCTTCAGCATCTTTTACTTTCACCAGCGTTTCTGGGTGAGCAAAAACAGGAAGGCAAAATGCCGCAAAAAAGGG<br>1041 AATAAGGGCGACACGGAAATGTTGAATACTCATACTCTTCCTTTTTCAATATTATTGAAGCATTTATCAGGGTTATTGTC<br>1121 TCATGAGCGGATACATATTTGAATGTATTTAGAAAAATAAACAAATAGGGGTTCCGCGCACATTTCCCCGAAAAGTGCCA<br>1201 CCTGACGTCTAAGAAACCATTATTATCATGACATTAACCTATAAAAATAGGCGTATCACGAGGCCCTTTCGTCTCGCGCG<br>1281 TTTCGGTGATGACGGTGAAAACCTCTGACACATGCAGCTCCCGGAGACGGTCACAGCTTGTCTGTAAGCGGATGCCGGGA<br>1361 GCAGACAAGCCCGTCAGGGCGCGTCAGCGGGTGTTGGCGGGTGTCGGGGCTGGCTTAACTATGCGGCATCAGAGCAGATT<br>1441 GTACTGAGAGTGCACCATATGCGGTGTGAAATACCGCACAGATGCGTAAGGAGAAAATACCGCATCAGGCGCCATTCGCC<br>1521 ATTCAGGCTGCGCAACTGTTGGGAAGGGCGATCGGTGCGGGCCTCTTCGCTATTACGCCAGCTGGCGAAAGGGGGATGTG<br>1601 CTGCAAGGCGATTAAGTTGGGTAACGCCAGGGTTTTCCCAGTCACGACGTTGTAAAACGACGGCCAGTAGATCTTATGGT<br>1681 GGTTTTGACTCTATCTCGCTTATTTAATTGTTTTAGTCAAGTTATTTATAGTGACATATATGTCTCTATTTTATTTTTCT<br>1761 ACATTCTTTTATAATAGGTCTAACGTCGACTCTCGAGTGAGATTGTTGACGGTACCGTATTTTGGATCTGAGCAAAGCAA<br>1841 TAACAGAACATAAGGAGGTACGTAATGAGCAAAGGAGAAGAACTTTTCACTGGAGTTGTCCCAATTCTTGTTGAATTAGA<br>1921 TGGTGATGTTAATGGGCACAAATTTTCTGTCCGTGGAGAGGGTGAAGGTGATGCTACAAACGGAAAACTCACCCTTAAAT<br>2001 TTATTTGCACTACTGGAAAACTACCTGTTCCGTGGCCAACACTTGTCACTACTCTGACCTATGGTGTTCAATGCTTTTCC<br>2081 CGTTATCCGGATCACATGAAACGGCATGACTTTTTCAAGAGTGCAATGCCCGAAGGTTATGTACAGGAACGCACTATATC<br>2161 TTTCAAAGATGACGGGACCTACAAGACGCGTGCTGAAGTCAAGTTTGAAGGTGATACCCTTGTTAATCGTATCGAGTTAA<br>2241 AGGGTATTGATTTTAAAGAAGATGGAAACATTCTTGGACACAAACTCGAGTACAACTTTAACTCACACAATGTATACATC |

| # | Type | Name and annotated features | Sequence (5′→3′) |
|---|---|---|---|
| | | | 2321 ACGGCAGACAAACAAAAGAATGGAATCAAAGCTAACTTCAAAATTCGCCACAACGTTGAAGATGGTTCCGTTCAACTAGC<br>2401 AGACCATTATCAACAAAATACTCCAATTGGCGATGGCCCTGTCCTTTTACCAGACAACCATTACCTGTCGACACAATCTG<br>2481 TCCTTTCGAAAGATCCCAACGAAAAGCGTGACCACATGGTCCTTCTTGAGTTTGTAACTGCTGCTGGGATTACACATGGC<br>2561 ATGGATGAGCTCTACAAATAAGGATCTGAAGCTTGGGCCCGAACAAAAACTCATCTCAGAAGAGGATCTGAATAGCGCCG<br>2641 TCGACCATCATCATCATCATCATTGAGTTTAAACGGACACCAGCTTGGCTGTTTTGGCGGATGAGAGAAGATTTTCAGCC<br>2721 TGATACAGATTAAATCAGAACGCAGAAGCGGTCTGATAAAACAGAATTTGCCTGGCGGCAGTAGCGCGGTGGTCCCACCT<br>2801 GACCCCATGCCGAACTCAGAAGTGAAACGCCGTAGCGCCGATGGTAGTGTGGGGACACCCCATGCGAGAGTAGGGAACTG<br>2881 CCAGGCATCAAATAAAACGAAAGGCTCAGTCGAAAGACTGGGCCTTTCGTTTTATCTGTTGTTTGTCGGTGAACTCTGAA<br>2961 AGGAGGAACTATATCCGGATCGAGATCCTCTAGAGTCGACCTGCAGGCATGCAAGCTTGGCGTAATCATGGTCATAGCTG<br>3041 TTTCCTGTGTGAAATTGTTATCCGCTCACAATTCCACACAACATACGAGCCGGAAGCATAAAGTGTAAAGCCTGGGGTGC<br>3121 CTAATGAGTGAGCTAACTCACATTAATTGCGTTGCGCTCACTGCCCGCTTTCCAGTCGGGAAACCTGTCGTGCCAGCTGC<br>3201 ATTAATGAATCGGCCAACGCGCGGGGAGAGGCGGTTTGCGTATTGGGCGCTCTTCCGCTTCCTCGCTCACTGACTCGCTG<br>3281 CGCTCGGTCGTTCGGCTGCGGCGAGCGGTATCAGCTCACTCAAAGGCGGTAATACGGTTATCCACAGAATCAGGGGATAA<br>3361 CGCAGGAAAGAACATGTGAGCAAAAGGCCAGCAAAGGCCAGGAACCGTAAAAAGGCCGCGTTGCTGGCGTTTTTCCATAG<br>3441 GCTCCGCCCCCCTGACGAGCATCACAAAAATCGACGCTCAAGTCAGAGGTGGCGAAACCCGACAGGACTATAAAGATACC<br>3521 AGGCGTTTCCCCCTGGAAGCTCCCTCGTGCGCTCTCCTGTTCCGACCCTGCCGCTTACCGGATACCTGTCCGCCTTTCTC<br>3601 CCTTCGGGAAGCGTGGCGCTTTCTCATAGCTCACGCTGTAGGTATCTCAGTTCGGTGTAGGTCGTTCGCTCCAAGCTGGG<br>3681 CTGTGTGCACGAACCCCCCGTTCAGCCCGACCGCTGCGCCTTATCCGGTAACTATCGTCTTGAGTCCAACCCGGTAAGAC<br>3761 ACGACTTATCGCCACTGGCAGCAGCCACTGGTAACAGGATTAGCAGAGCGAGGTATGTAGGCGGTGCTACAGAGTTCTTG<br>3841 AAGTGGTGGCCTAACTACGGCTACACTAGAAGAACAGTATTTGGTATCTGCGCTCTGCTGAAGCCAGTTACCTTCGGAAA<br>3921 AAGAGTTGGTAGCTCTTGATCCGGCAAACAAACCACCGCTGGTAGCGGTGGTTTTTTT |
| 9 | Plasmid | **pComR-SI-sfGFP**<br>*Graham_Hendrickson_Q6Y_4_ComR_Si_prom_WT_pLann · 3977 bp · circular*<br>Fig. 1; Suppl. Figs 2–5, 9<br><br>**Annotated features**<br>AmpR 213–1073<br>AmpR promoter 1074–1178<br>rop (fragment) 1273–1308<br>bom (fragment) 1410–1509<br>lacZα (fragment) 1515–1668<br>ComR-SI promoter 1679–1781<br>sfGFP 1864–2577<br>5S rRNA 2771–2886<br>rrnB T1 terminator 2888–2934<br>MCS (fragment) 2983–3017<br>lac operator 3054–3070<br>lac promoter 3078–3108<br>CAP binding site 3123–3144<br>lacI (fragment) 3157–3249<br>ori 3431–3977, 1–42 | 1 GTTTGCAAGCAGCAGATTACGCGCAGAAAAAAAGGATCTCAAGAAGATCCTTTGATCTTTTCTACGGGGTCTGACGCTCA<br>81 GTGGAACGAAAACTCACGTTAAGGGATTTTGGTCATGAGATTATCAAAAAGGATCTTCACCTAGATCCTTTTAAATTAAA<br>161 AATGAAGTTTTAAATCAATCTAAAGTATATATGAGTAAACTTGGTCTGACAGTTACCAATGCTTAATCAGTGAGGCACCT<br>241 ATCTCAGCGATCTGTCTATTTCGTTCATCCATAGTTGCCTGACTCCCCGTCGTGTAGATAACTACGATACGGGAGGGCTT<br>321 ACCATCTGGCCCCAGTGCTGCAATGATACCGCGACTCCCACGCTCACCGGCTCCAGATTTATCAGCAATAAACCAGCCAG<br>401 CCGGAAGGGCCGAGCGCAGAAGTGGTCCTGCAACTTTATCCGCCTCCATCCAGTCTATTAATTGTTGCCGGGAAGCTAGA<br>481 GTAAGTAGTTCGCCAGTTAATAGTTTGCGCAACGTTGTTGCCATTGCTACAGGCATCGTGGTGTCACGCTCGTCGTTTGG<br>561 TATGGCTTCATTCAGCTCCGGTTCCCAACGATCAAGGCGAGTTACATGATCCCCCATGTTGTGCAAAAAAGCGGTTAGCT<br>641 CCTTCGGTCCTCCGATCGTTGTCAGAAGTAAGTTGGCCGCAGTGTTATCACTCATGGTTATGGCAGCACTGCATAATTCT<br>721 CTTACTGTCATGCCATCCGTAAGATGCTTTTCTGTGACTGGTGAGTACTCAACCAAGTCATTCTGAGAATAGTGTATGCG<br>801 GCGACCGAGTTGCTCTTGCCCGGCGTCAATACGGGATAATACCGCGCCACATAGCAGAACTTTAAAAGTGCTCATCATTG<br>881 GAAAACGTTCTTCGGGGCGAAAACTCTCAAGGATCTTACCGCTGTTGAGATCCAGTTCGATGTAACCCACTCGTGCACCC<br>961 AACTGATCTTCAGCATCTTTTACTTTCACCAGCGTTTCTGGGTGAGCAAAAACAGGAAGGCAAAATGCCGCAAAAAAGGG<br>1041 AATAAGGGCGACACGGAAATGTTGAATACTCATACTCTTCCTTTTTCAATATTATTGAAGCATTTATCAGGGTTATTGTC<br>1121 TCATGAGCGGATACATATTTGAATGTATTTAGAAAAATAAACAAATAGGGGTTCCGCGCACATTTCCCCGAAAAGTGCCA<br>1201 CCTGACGTCTAAGAAACCATTATTATCATGACATTAACCTATAAAAATAGGCGTATCACGAGGCCCTTTCGTCTCGCGCG<br>1281 TTTCGGTGATGACGGTGAAAACCTCTGACACATGCAGCTCCCGGAGACGGTCACAGCTTGTCTGTAAGCGGATGCCGGGA<br>1361 GCAGACAAGCCCGTCAGGGCGCGTCAGCGGGTGTTGGCGGGTGTCGGGGCTGGCTTAACTATGCGGCATCAGAGCAGATT<br>1441 GTACTGAGAGTGCACCATATGCGGTGTGAAATACCGCACAGATGCGTAAGGAGAAAATACCGCATCAGGCGCCATTCGCC<br>1521 ATTCAGGCTGCGCAACTGTTGGGAAGGGCGATCGGTGCGGGCCTCTTCGCTATTACGCCAGCTGGCGAAAGGGGGATGTG<br>1601 CTGCAAGGCGATTAAGTTGGGTAACGCCAGGGTTTTCCCAGTCACGACGTTGTAAAACGACGGCCAGTAGATCTTATGAA<br>1681 GCGGTAGACGGAGAGCAACTCAGATGAGTTGCTTTTTTTGTGCATCAATTTGGACATTTTTGTCATGTTCCTTTTCCCTA<br>1761 ATCCTTTCTATAATAAGGGTAACGTCGACTCTCGAGTGAGATTGTTGACGGTACCGTATTTTGGATCTGAGCAAAGCAAT<br>1841 AACAGAACATAAGGAGGTACGTAATGAGCAAAGGAGAAGAACTTTTCACTGGAGTTGTCCCAATTCTTGTTGAATTAGAT<br>1921 GGTGATGTTAATGGGCACAAATTTTCTGTCCGTGGAGAGGGTGAAGGTGATGCTACAAACGGAAAACTCACCCTTAAATT<br>2001 TATTTGCACTACTGGAAAACTACCTGTTCCGTGGCCAACACTTGTCACTACTCTGACCTATGGTGTTCAATGCTTTTCCC<br>2081 GTTATCCGGATCACATGAAACGGCATGACTTTTTCAAGAGTGCAATGCCCGAAGGTTATGTACAGGAACGCACTATATCT<br>2161 TTCAAAGATGACGGGACCTACAAGACGCGTGCTGAAGTCAAGTTTGAAGGTGATACCCTTGTTAATCGTATCGAGTTAAA<br>2241 GGGTATTGATTTTAAAGAAGATGGAAACATTCTTGGACACAAACTCGAGTACAACTTTAACTCACACAATGTATACATCA<br>2321 CGGCAGACAAACAAAAGAATGGAATCAAAGCTAACTTCAAAATTCGCCACAACGTTGAAGATGGTTCCGTTCAACTAGCA<br>2401 GACCATTATCAACAAAATACTCCAATTGGCGATGGCCCTGTCCTTTTACCAGACAACCATTACCTGTCGACACAATCTGT<br>2481 CCTTTCGAAAGATCCCAACGAAAAGCGTGACCACATGGTCCTTCTTGAGTTTGTAACTGCTGCTGGGATTACACATGGCA<br>2561 TGGATGAGCTCTACAAATAAGGATCTGAAGCTTGGGCCCGAACAAAAACTCATCTCAGAAGAGGATCTGAATAGCGCCGT<br>2641 CGACCATCATCATCATCATCATTGAGTTTAAACGGACACCAGCTTGGCTGTTTTGGCGGATGAGAGAAGATTTTCAGCCT<br>2721 GATACAGATTAAATCAGAACGCAGAAGCGGTCTGATAAAACAGAATTTGCCTGGCGGCAGTAGCGCGGTGGTCCCACCTG<br>2801 ACCCCATGCCGAACTCAGAAGTGAAACGCCGTAGCGCCGATGGTAGTGTGGGGACACCCCATGCGAGAGTAGGGAACTGC<br>2881 CAGGCATCAAATAAAACGAAAGGCTCAGTCGAAAGACTGGGCCTTTCGTTTTATCTGTTGTTTGTCGGTGAACTCTGAAA<br>2961 GGAGGAACTATATCCGGATCGAGATCCTCTAGAGTCGACCTGCAGGCATGCAAGCTTGGCGTAATCATGGTCATAGCTGT<br>3041 TTCCTGTGTGAAATTGTTATCCGCTCACAATTCCACACAACATACGAGCCGGAAGCATAAAGTGTAAAGCCTGGGGTGCC<br>3121 TAATGAGTGAGCTAACTCACATTAATTGCGTTGCGCTCACTGCCCGCTTTCCAGTCGGGAAACCTGTCGTGCCAGCTGCA<br>3201 TTAATGAATCGGCCAACGCGCGGGGAGAGGCGGTTTGCGTATTGGGCGCTCTTCCGCTTCCTCGCTCACTGACTCGCTGC<br>3281 GCTCGGTCGTTCGGCTGCGGCGAGCGGTATCAGCTCACTCAAAGGCGGTAATACGGTTATCCACAGAATCAGGGGATAAC<br>3361 GCAGGAAAGAACATGTGAGCAAAAGGCCAGCAAAGGCCAGGAACCGTAAAAAGGCCGCGTTGCTGGCGTTTTTCCATAGG<br>3441 CTCCGCCCCCCTGACGAGCATCACAAAAATCGACGCTCAAGTCAGAGGTGGCGAAACCCGACAGGACTATAAAGATACCA<br>3521 GGCGTTTCCCCCTGGAAGCTCCCTCGTGCGCTCTCCTGTTCCGACCCTGCCGCTTACCGGATACCTGTCCGCCTTTCTCC<br>3601 CTTCGGGAAGCGTGGCGCTTTCTCATAGCTCACGCTGTAGGTATCTCAGTTCGGTGTAGGTCGTTCGCTCCAAGCTGGGC<br>3681 TGTGTGCACGAACCCCCCGTTCAGCCCGACCGCTGCGCCTTATCCGGTAACTATCGTCTTGAGTCCAACCCGGTAAGACA<br>3761 CGACTTATCGCCACTGGCAGCAGCCACTGGTAACAGGATTAGCAGAGCGAGGTATGTAGGCGGTGCTACAGAGTTCTTGA<br>3841 AGTGGTGGCCTAACTACGGCTACACTAGAAGAACAGTATTTGGTATCTGCGCTCTGCTGAAGCCAGTTACCTTCGGAAAA |

| # | Type | Name and annotated features | Sequence (5′→3′) |
|---|---|---|---|
| | | | 3921 AGAGTTGGTAGCTCTTGATCCGGCAAACAAACCACCGCTGGTAGCGGTGGTTTTTTT |
| 10 | Plasmid | **pPlcR-sfGFP**<br>*Graham_Hendrickson_Q6Y_5_PlcR_prom_WT_pLann · 3991 bp · circular*<br>Fig. 1; Suppl. Figs 2–4, 9<br><br>**Annotated features**<br>AmpR 213–1073<br>AmpR promoter 1074–1178<br>rop (fragment) 1273–1308<br>bom (fragment) 1410–1509<br>lacZα (fragment) 1515–1668<br>PlcR promoter 1679–1795<br>sfGFP 1878–2591<br>5S rRNA 2785–2900<br>rrnB T1 terminator 2902–2948<br>MCS (fragment) 2997–3031<br>lac operator 3068–3084<br>lac promoter 3092–3122<br>CAP binding site 3137–3158<br>lacI (fragment) 3171–3263<br>ori 3445–3991, 1–42 | 1 GTTTGCAAGCAGCAGATTACGCGCAGAAAAAAAGGATCTCAAGAAGATCCTTTGATCTTTTCTACGGGGTCTGACGCTCA<br>81 GTGGAACGAAAACTCACGTTAAGGGATTTTGGTCATGAGATTATCAAAAAGGATCTTCACCTAGATCCTTTTAAATTAAA<br>161 AATGAAGTTTTAAATCAATCTAAAGTATATATGAGTAAACTTGGTCTGACAGTTACCAATGCTTAATCAGTGAGGCACCT<br>241 ATCTCAGCGATCTGTCTATTTCGTTCATCCATAGTTGCCTGACTCCCCGTCGTGTAGATAACTACGATACGGGAGGGCTT<br>321 ACCATCTGGCCCCAGTGCTGCAATGATACCGCGACTCCCACGCTCACCGGCTCCAGATTTATCAGCAATAAACCAGCCAG<br>401 CCGGAAGGGCCGAGCGCAGAAGTGGTCCTGCAACTTTATCCGCCTCCATCCAGTCTATTAATTGTTGCCGGGAAGCTAGA<br>481 GTAAGTAGTTCGCCAGTTAATAGTTTGCGCAACGTTGTTGCCATTGCTACAGGCATCGTGGTGTCACGCTCGTCGTTTGG<br>561 TATGGCTTCATTCAGCTCCGGTTCCCAACGATCAAGGCGAGTTACATGATCCCCCATGTTGTGCAAAAAAGCGGTTAGCT<br>641 CCTTCGGTCCTCCGATCGTTGTCAGAAGTAAGTTGGCCGCAGTGTTATCACTCATGGTTATGGCAGCACTGCATAATTCT<br>721 CTTACTGTCATGCCATCCGTAAGATGCTTTTCTGTGACTGGTGAGTACTCAACCAAGTCATTCTGAGAATAGTGTATGCG<br>801 GCGACCGAGTTGCTCTTGCCCGGCGTCAATACGGGATAATACCGCGCCACATAGCAGAACTTTAAAAGTGCTCATCATTG<br>881 GAAAACGTTCTTCGGGGCGAAAACTCTCAAGGATCTTACCGCTGTTGAGATCCAGTTCGATGTAACCCACTCGTGCACCC<br>961 AACTGATCTTCAGCATCTTTTACTTTCACCAGCGTTTCTGGGTGAGCAAAAACAGGAAGGCAAAATGCCGCAAAAAAGGG<br>1041 AATAAGGGCGACACGGAAATGTTGAATACTCATACTCTTCCTTTTTCAATATTATTGAAGCATTTATCAGGGTTATTGTC<br>1121 TCATGAGCGGATACATATTTGAATGTATTTAGAAAAATAAACAAATAGGGGTTCCGCGCACATTTCCCCGAAAAGTGCCA<br>1201 CCTGACGTCTAAGAAACCATTATTATCATGACATTAACCTATAAAAATAGGCGTATCACGAGGCCCTTTCGTCTCGCGCG<br>1281 TTTCGGTGATGACGGTGAAAACCTCTGACACATGCAGCTCCCGGAGACGGTCACAGCTTGTCTGTAAGCGGATGCCGGGA<br>1361 GCAGACAAGCCCGTCAGGGCGCGTCAGCGGGTGTTGGCGGGTGTCGGGGCTGGCTTAACTATGCGGCATCAGAGCAGATT<br>1441 GTACTGAGAGTGCACCATATGCGGTGTGAAATACCGCACAGATGCGTAAGGAGAAAATACCGCATCAGGCGCCATTCGCC<br>1521 ATTCAGGCTGCGCAACTGTTGGGAAGGGCGATCGGTGCGGGCCTCTTCGCTATTACGCCAGCTGGCGAAAGGGGGATGTG<br>1601 CTGCAAGGCGATTAAGTTGGGTAACGCCAGGGTTTTCCCAGTCACGACGTTGTAAAACGACGGCCAGTAGATCTTATGAA<br>1681 ATGAAAGTGTATTCATTCATTATATTCACTGTGTATAAAGTTATAATGATATGAACATTTGCATATTTTAATTTAGTGAT<br>1761 AGAAATTTCGTGAAAGGTGGGATATTCTAGTCATAACGTCGACTCTCGAGTGAGATTGTTGACGGTACCGTATTTTGGAT<br>1841 CTGAGCAAAGCAATAACAGAACATAAGGAGGTACGTAATGAGCAAAGGAGAAGAACTTTTCACTGGAGTTGTCCCAATTC<br>1921 TTGTTGAATTAGATGGTGATGTTAATGGGCACAAATTTTCTGTCCGTGGAGAGGGTGAAGGTGATGCTACAAACGGAAAA<br>2001 CTCACCCTTAAATTTATTTGCACTACTGGAAAACTACCTGTTCCGTGGCCAACACTTGTCACTACTCTGACCTATGGTGT<br>2081 TCAATGCTTTTCCCGTTATCCGGATCACATGAAACGGCATGACTTTTTCAAGAGTGCAATGCCCGAAGGTTATGTACAGG<br>2161 AACGCACTATATCTTTCAAAGATGACGGGACCTACAAGACGCGTGCTGAAGTCAAGTTTGAAGGTGATACCCTTGTTAAT<br>2241 CGTATCGAGTTAAAGGGTATTGATTTTAAAGAAGATGGAAACATTCTTGGACACAAACTCGAGTACAACTTTAACTCACA<br>2321 CAATGTATACATCACGGCAGACAAACAAAAGAATGGAATCAAAGCTAACTTCAAAATTCGCCACAACGTTGAAGATGGTT<br>2401 CCGTTCAACTAGCAGACCATTATCAACAAAATACTCCAATTGGCGATGGCCCTGTCCTTTTACCAGACAACCATTACCTG<br>2481 TCGACACAATCTGTCCTTTCGAAAGATCCCAACGAAAAGCGTGACCACATGGTCCTTCTTGAGTTTGTAACTGCTGCTGG<br>2561 GATTACACATGGCATGGATGAGCTCTACAAATAAGGATCTGAAGCTTGGGCCCGAACAAAAACTCATCTCAGAAGAGGAT<br>2641 CTGAATAGCGCCGTCGACCATCATCATCATCATCATTGAGTTTAAACGGACACCAGCTTGGCTGTTTTGGCGGATGAGAG<br>2721 AAGATTTTCAGCCTGATACAGATTAAATCAGAACGCAGAAGCGGTCTGATAAAACAGAATTTGCCTGGCGGCAGTAGCGC<br>2801 GGTGGTCCCACCTGACCCCATGCCGAACTCAGAAGTGAAACGCCGTAGCGCCGATGGTAGTGTGGGGACACCCCATGCGA<br>2881 GAGTAGGGAACTGCCAGGCATCAAATAAAACGAAAGGCTCAGTCGAAAGACTGGGCCTTTCGTTTTATCTGTTGTTTGTC<br>2961 GGTGAACTCTGAAAGGAGGAACTATATCCGGATCGAGATCCTCTAGAGTCGACCTGCAGGCATGCAAGCTTGGCGTAATC<br>3041 ATGGTCATAGCTGTTTCCTGTGTGAAATTGTTATCCGCTCACAATTCCACACAACATACGAGCCGGAAGCATAAAGTGTA<br>3121 AAGCCTGGGGTGCCTAATGAGTGAGCTAACTCACATTAATTGCGTTGCGCTCACTGCCCGCTTTCCAGTCGGGAAACCTG<br>3201 TCGTGCCAGCTGCATTAATGAATCGGCCAACGCGCGGGGAGAGGCGGTTTGCGTATTGGGCGCTCTTCCGCTTCCTCGCT<br>3281 CACTGACTCGCTGCGCTCGGTCGTTCGGCTGCGGCGAGCGGTATCAGCTCACTCAAAGGCGGTAATACGGTTATCCACAG<br>3361 AATCAGGGGATAACGCAGGAAAGAACATGTGAGCAAAAGGCCAGCAAAAGGCCAGGAACCGTAAAAAGGCCGCGTTGCTGG<br>3441 CGTTTTTCCATAGGCTCCGCCCCCCTGACGAGCATCACAAAAATCGACGCTCAAGTCAGAGGTGGCGAAACCCGACAGGA<br>3521 CTATAAAGATACCAGGCGTTTCCCCCTGGAAGCTCCCTCGTGCGCTCTCCTGTTCCGACCCTGCCGCTTACCGGATACCT<br>3601 GTCCGCCTTTCTCCCTTCGGGAAGCGTGGCGCTTTCTCATAGCTCACGCTGTAGGTATCTCAGTTCGGTGTAGGTCGTTC<br>3681 GCTCCAAGCTGGGCTGTGTGCACGAACCCCCCGTTCAGCCCGACCGCTGCGCCTTATCCGGTAACTATCGTCTTGAGTCC<br>3761 AACCCGGTAAGACACGACTTATCGCCACTGGCAGCAGCCACTGGTAACAGGATTAGCAGAGCGAGGTATGTAGGCGGTGC<br>3841 TACAGAGTTCTTGAAGTGGTGGCCTAACTACGGCTACACTAGAAGAACAGTATTTGGTATCTGCGCTCTGCTGAAGCCAG<br>3921 TTACCTTCGGAAAAAGAGTTGGTAGCTCTTGATCCGGCAAACAAACCACCGCTGGTAGCGGTGGTTTTTTT |

**C. Constitutive peptide expression templates (pJ23119-peptide)**

| # | Type | Name and annotated features | Sequence (5′→3′) |
|---|---|---|---|
| 11 | LET | **pJ23119-SHP2**<br>*A_J23119_SHP2 · 960 bp · linear*<br>Fig. 2a–b<br>Encoded peptide: **DILIIVGG**<br><br>**Annotated features**<br>lacZα (fragment) 67–135<br>J23119 promoter 251–285<br>RBS (strong) 293–306<br>SHP2 Cognate Peptide 310–333<br>BamHI-BglII Scar 337–342<br>rrnB terminator (TrrnB) 343–710<br>lac operator 808–830 | 1 GAGAGTGCACCATATGCGGTGTGAAATACCGCACAGATGCGTAAGGAGAAAATACCGCATCAGGCGCCATTCGCCATTCA<br>81 GGCTGCGCAACTGTTGGGAAGGGCGATCGGTGCGGGCCTCTTCGCTATTACGCCAGCTGGCGAAAGGGGGATGTGCTGCA<br>161 AGGCGATTAAGTTGGGTAACGCCAGGGTTTTCCCAGTCACGACGTTGTAAAACGACGGCCAGTAAGATCTCGGCTGAATT<br>241 CTAAAGATCTTTGACAGCTAGCTCAGTCCTAGGTATAATACTAGTATAATTAAGGAGGAAGGATCTATGGATATCCTGAT<br>321 TATCGTGGGTGGTTAAGGATCTGAAGCTTGGGCCCGAACAAAAACTCATCTCAGAAGAGGATCTGAATAGCGCCGTCGAC<br>401 CATCATCATCATCATCATTGAGTTTAAACGGTCTCCAGCTTGGCTGTTTTGGCGGATGAGAGAAGATTTTCAGCCTGATA<br>481 CAGATTAAATCAGAACGCAGAAGCGGTCTGATAAAACAGAATTTGCCTGGCGGCAGTAGCGCGGTGGTCCCACCTGACCC<br>561 CATGCCGAACTCAGAAGTGAAACGCCGTAGCGCCGATGGTAGTGTGGGGTCTCCCCATGCGAGAGTAGGGAACTGCCAGG<br>641 CATCAAATAAAACGAAAGGCTCAGTCGAAAGACTGGGCCTTTCGTTTTATCTGTTGTTTGTCGGTGAACTCTGAAAGGAG<br>721 GAACTATATCCGGATCGAGATCCTCTAGAGTCGACCTGCAGGCATGCAAGCTTGGCGTAATCATGGTCATAGCTGTTTCC<br>801 TGTGTGAAATTGTTATCCGCTCACAATTCCACACAACATACGAGCCGGAAGCATAAAGTGTAAAGCCTGGGGTGCCTAAT<br>881 GAGTGAGCTAACTCACATTAATTGCGTTGCGCTCACTGCCCGCTTTCCAGTCGGGAAACCTGTCGTGCCAGCTGCATTAA |
| 12 | LET | **pJ23119-SHP3**<br>*A_J23119_SHP3 · 960 bp · linear*<br>Fig. 2a, 2c<br>Encoded peptide: **DIIIIVGG**<br><br>**Annotated features**<br>lacZα (fragment) 67–135<br>J23119 promoter 251–285<br>RBS (strong) 293–306<br>SHP3 Cognate Peptide 310–333<br>BamHI-BglII Scar 337–342<br>rrnB terminator (TrrnB) 343–710<br>lac operator 808–830 | 1 GAGAGTGCACCATATGCGGTGTGAAATACCGCACAGATGCGTAAGGAGAAAATACCGCATCAGGCGCCATTCGCCATTCA<br>81 GGCTGCGCAACTGTTGGGAAGGGCGATCGGTGCGGGCCTCTTCGCTATTACGCCAGCTGGCGAAAGGGGGATGTGCTGCA<br>161 AGGCGATTAAGTTGGGTAACGCCAGGGTTTTCCCAGTCACGACGTTGTAAAACGACGGCCAGTAAGATCTCGGCTGAATT<br>241 CTAAAGATCTTTGACAGCTAGCTCAGTCCTAGGTATAATACTAGTATAATTAAGGAGGAAGGATCTATGGATATCATTAT<br>321 TATCGTGGGTGGTTAAGGATCTGAAGCTTGGGCCCGAACAAAAACTCATCTCAGAAGAGGATCTGAATAGCGCCGTCGAC<br>401 CATCATCATCATCATCATTGAGTTTAAACGGTCTCCAGCTTGGCTGTTTTGGCGGATGAGAGAAGATTTTCAGCCTGATA<br>481 CAGATTAAATCAGAACGCAGAAGCGGTCTGATAAAACAGAATTTGCCTGGCGGCAGTAGCGCGGTGGTCCCACCTGACCC<br>561 CATGCCGAACTCAGAAGTGAAACGCCGTAGCGCCGATGGTAGTGTGGGGTCTCCCCATGCGAGAGTAGGGAACTGCCAGG<br>641 CATCAAATAAAACGAAAGGCTCAGTCGAAAGACTGGGCCTTTCGTTTTATCTGTTGTTTGTCGGTGAACTCTGAAAGGAG<br>721 GAACTATATCCGGATCGAGATCCTCTAGAGTCGACCTGCAGGCATGCAAGCTTGGCGTAATCATGGTCATAGCTGTTTCC<br>801 TGTGTGAAATTGTTATCCGCTCACAATTCCACACAACATACGAGCCGGAAGCATAAAGTGTAAAGCCTGGGGTGCCTAAT<br>881 GAGTGAGCTAACTCACATTAATTGCGTTGCGCTCACTGCCCGCTTTCCAGTCGGGAAACCTGTCGTGCCAGCTGCATTAA |
| 13 | LET | **pJ23119-XIP-ST**<br>*A_J23119_XIPST · 957 bp · linear*<br>Suppl. Fig. 5a–b<br>Encoded peptide: **PYFAGCL**<br><br>**Annotated features**<br>lacZα (fragment) 67–135<br>J23119 promoter 251–285<br>RBS (strong) 293–306<br>XIP-ST Peptide 310–330<br>BamHI-BglII Scar 334–339<br>rrnB terminator (TrrnB) 340–707<br>lac operator 805–827 | 1 GAGAGTGCACCATATGCGGTGTGAAATACCGCACAGATGCGTAAGGAGAAAATACCGCATCAGGCGCCATTCGCCATTCA<br>81 GGCTGCGCAACTGTTGGGAAGGGCGATCGGTGCGGGCCTCTTCGCTATTACGCCAGCTGGCGAAAGGGGGATGTGCTGCA<br>161 AGGCGATTAAGTTGGGTAACGCCAGGGTTTTCCCAGTCACGACGTTGTAAAACGACGGCCAGTAAGATCTCGGCTGAATT<br>241 CTAAAGATCTTTGACAGCTAGCTCAGTCCTAGGTATAATACTAGTATAATTAAGGAGGAAGGATCTATGCCATACTTCGC<br>321 CGGCTGTCTGTAAGGATCTGAAGCTTGGGCCCGAACAAAAACTCATCTCAGAAGAGGATCTGAATAGCGCCGTCGACCAT<br>401 CATCATCATCATCATTGAGTTTAAACGGTCTCCAGCTTGGCTGTTTTGGCGGATGAGAGAAGATTTTCAGCCTGATACAG<br>481 ATTAAATCAGAACGCAGAAGCGGTCTGATAAAACAGAATTTGCCTGGCGGCAGTAGCGCGGTGGTCCCACCTGACCCCAT<br>561 GCCGAACTCAGAAGTGAAACGCCGTAGCGCCGATGGTAGTGTGGGGTCTCCCCATGCGAGAGTAGGGAACTGCCAGGCAT<br>641 CAAATAAAACGAAAGGCTCAGTCGAAAGACTGGGCCTTTCGTTTTATCTGTTGTTTGTCGGTGAACTCTGAAAGGAGGAA<br>721 CTATATCCGGATCGAGATCCTCTAGAGTCGACCTGCAGGCATGCAAGCTTGGCGTAATCATGGTCATAGCTGTTTCCTGT<br>801 GTGAAATTGTTATCCGCTCACAATTCCACACAACATACGAGCCGGAAGCATAAAGTGTAAAGCCTGGGGTGCCTAATGAG<br>881 TGAGCTAACTCACATTAATTGCGTTGCGCTCACTGCCCGCTTTCCAGTCGGGAAACCTGTCGTGCCAGCTGCATTAA |
| 14 | LET | **pJ23119-XIP-SI**<br>*A_J23119_XIPSI · 960 bp · linear*<br>Suppl. Fig. 5a, 5c<br>Encoded peptide: **LLTAWWGL**<br><br>**Annotated features**<br>lacZα (fragment) 67–135<br>J23119 promoter 251–285<br>RBS (strong) 293–306<br>XIP-SI Peptide 310–333<br>BamHI-BglII Scar 337–342<br>rrnB terminator (TrrnB) 343–710<br>lac operator 808–830 | 1 GAGAGTGCACCATATGCGGTGTGAAATACCGCACAGATGCGTAAGGAGAAAATACCGCATCAGGCGCCATTCGCCATTCA<br>81 GGCTGCGCAACTGTTGGGAAGGGCGATCGGTGCGGGCCTCTTCGCTATTACGCCAGCTGGCGAAAGGGGGATGTGCTGCA<br>161 AGGCGATTAAGTTGGGTAACGCCAGGGTTTTCCCAGTCACGACGTTGTAAAACGACGGCCAGTAAGATCTCGGCTGAATT<br>241 CTAAAGATCTTTGACAGCTAGCTCAGTCCTAGGTATAATACTAGTATAATTAAGGAGGAAGGATCTATGCTCCTCACAGC<br>321 GTGGTGGGGCCTATAAGGATCTGAAGCTTGGGCCCGAACAAAAACTCATCTCAGAAGAGGATCTGAATAGCGCCGTCGAC<br>401 CATCATCATCATCATCATTGAGTTTAAACGGTCTCCAGCTTGGCTGTTTTGGCGGATGAGAGAAGATTTTCAGCCTGATA<br>481 CAGATTAAATCAGAACGCAGAAGCGGTCTGATAAAACAGAATTTGCCTGGCGGCAGTAGCGCGGTGGTCCCACCTGACCC<br>561 CATGCCGAACTCAGAAGTGAAACGCCGTAGCGCCGATGGTAGTGTGGGGTCTCCCCATGCGAGAGTAGGGAACTGCCAGG<br>641 CATCAAATAAAACGAAAGGCTCAGTCGAAAGACTGGGCCTTTCGTTTTATCTGTTGTTTGTCGGTGAACTCTGAAAGGAG<br>721 GAACTATATCCGGATCGAGATCCTCTAGAGTCGACCTGCAGGCATGCAAGCTTGGCGTAATCATGGTCATAGCTGTTTCC<br>801 TGTGTGAAATTGTTATCCGCTCACAATTCCACACAACATACGAGCCGGAAGCATAAAGTGTAAAGCCTGGGGTGCCTAAT<br>881 GAGTGAGCTAACTCACATTAATTGCGTTGCGCTCACTGCCCGCTTTCCAGTCGGGAAACCTGTCGTGCCAGCTGCATTAA |

| # | Type | Name and annotated features | Sequence (5′→3′) |
|---|---|---|---|
| **D. Self-activating peptide expression templates (pRRNPP-peptide)** | | | |
| 15 | LET | **pRgg2-SHP2**<br>*Rgg2O_SHP2_Peptide_Expression · 1078 bp · linear*<br>Fig. 2d–e<br>Encoded peptide: **DILIIVGG**<br><br>**Annotated features**<br>lacZα (fragment) 67–135<br>Rgg2 promoter 251–359<br>pHP14 mRNA stability hairpin 364–404<br>BamHI-BglII Scar 405–410<br>RBS (strong) 411–424<br>Start codon (ATG) 425–427<br>SHP2 Cognate Peptide 428–451<br>rrnB terminator (TrrnB) 461–828<br>lac operator 926–948 | 1 GAGAGTGCACCATATGCGGTGTGAAATACCGCACAGATGCGTAAGGAGAAAATACCGCATCAGGCGCCATTCGCCATTCA<br>81 GGCTGCGCAACTGTTGGGAAGGGCGATCGGTGCGGGCCTCTTCGCTATTACGCCAGCTGGCGAAAGGGGGATGTGCTGCA<br>161 AGGCGATTAAGTTGGGTAACGCCAGGGTTTTCCCAGTCACGACGTTGTAAAACGACGGCCAGTAAGATCTCGGCTGAATT<br>241 CTAAAGATCTTGTTTTCCTTGGCGTAGTCGCCTTAAGGTTTTACCGAGTTCTTTTTCCATTTTTCCCACTTTCACAACAA<br>321 AAAATTCAAATCCGTCAAATCCATTATATAATACAAGTAGGCTACGTCGACTCTCGAGTGAGATTGTTGACGGTACCGTA<br>401 TTTTGGATCTAGGAGGAAGGATCTATGGATATCCTGATTATCGTGGGTGGTTAAGGATCTGAAGCTTGGGCCCGAACAAA<br>481 AACTCATCTCAGAAGAGGATCTGAATAGCGCCGTCGACCATCATCATCATCATCATTGAGTTTAAACGGTCTCCAGCTTG<br>561 GCTGTTTTGGCGGATGAGAGAAGATTTTCAGCCTGATACAGATTAAATCAGAACGCAGAAGCGGTCTGATAAAACAGAAT<br>641 TTGCCTGGCGGCAGTAGCGCGGTGGTCCCACCTGACCCCATGCCGAACTCAGAAGTGAAACGCCGTAGCGCCGATGGTAG<br>721 TGTGGGGTCTCCCCATGCGAGAGTAGGGAACTGCCAGGCATCAAATAAAACGAAAGGCTCAGTCGAAAGACTGGGCCTTT<br>801 CGTTTTATCTGTTGTTTGTCGGTGAACTCTGAAAGGAGGAACTATATCCGGATCGAGATCCTCTAGAGTCGACCTGCAGG<br>881 CATGCAAGCTTGGCGTAATCATGGTCATAGCTGTTTCCTGTGTGAAATTGTTATCCGCTCACAATTCCACACAACATACG<br>961 AGCCGGAAGCATAAAGTGTAAAGCCTGGGGTGCCTAATGAGTGAGCTAACTCACATTAATTGCGTTGCGCTCACTGCCCG<br>1041 CTTTCCAGTCGGGAAACCTGTCGTGCCAGCTGCATTAA |
| 16 | LET | **pRgg3-SHP3**<br>*Rgg3O_SHP3_Peptide_Expression · 1080 bp · linear*<br>Fig. 2d, 2f; Suppl. Fig. 12<br>Encoded peptide: **DIIIIVGG**<br><br>**Annotated features**<br>lacZα (fragment) 67–135<br>Rgg3 promoter 251–361<br>Rgg binding site 304–328<br>-10 345–350<br>Transcription start site 358–358<br>pHP14 mRNA stability hairpin 366–406<br>BamHI-BglII Scar 407–412<br>RBS (strong) 413–426<br>Start codon (ATG) 427–429<br>SHP3 Cognate Peptide 430–453<br>rrnB terminator (TrrnB) 463–830<br>lac operator 928–950 | 1 GAGAGTGCACCATATGCGGTGTGAAATACCGCACAGATGCGTAAGGAGAAAATACCGCATCAGGCGCCATTCGCCATTCA<br>81 GGCTGCGCAACTGTTGGGAAGGGCGATCGGTGCGGGCCTCTTCGCTATTACGCCAGCTGGCGAAAGGGGGATGTGCTGCA<br>161 AGGCGATTAAGTTGGGTAACGCCAGGGTTTTCCCAGTCACGACGTTGTAAAACGACGGCCAGTAAGATCTCGGCTGAATT<br>241 CTAAAGATCTTATGGCTTTCCTTTTCGAACTTTTCTAAGTGTGACACCAAGTTTTGATTTCATATTTTCCCACTTTCCCA<br>321 ACAAAAATCAATTCCGTTTTTTTTTATTGTACAATAAAACGGGCTACGTCGACTCTCGAGTGAGATTGTTGACGGTACCG<br>401 TATTTTGGATCTAGGAGGAAGGATCTATGGATATCATTATTATCGTGGGTGGTTAAGGATCTGAAGCTTGGGCCCGAACA<br>481 AAAACTCATCTCAGAAGAGGATCTGAATAGCGCCGTCGACCATCATCATCATCATCATTGAGTTTAAACGGTCTCCAGCT<br>561 TGGCTGTTTTGGCGGATGAGAGAAGATTTTCAGCCTGATACAGATTAAATCAGAACGCAGAAGCGGTCTGATAAAACAGA<br>641 ATTTGCCTGGCGGCAGTAGCGCGGTGGTCCCACCTGACCCCATGCCGAACTCAGAAGTGAAACGCCGTAGCGCCGATGGT<br>721 AGTGTGGGGTCTCCCCATGCGAGAGTAGGGAACTGCCAGGCATCAAATAAAACGAAAGGCTCAGTCGAAAGACTGGGCCT<br>801 TTCGTTTTATCTGTTGTTTGTCGGTGAACTCTGAAAGGAGGAACTATATCCGGATCGAGATCCTCTAGAGTCGACCTGCA<br>881 GGCATGCAAGCTTGGCGTAATCATGGTCATAGCTGTTTCCTGTGTGAAATTGTTATCCGCTCACAATTCCACACAACATA<br>961 CGAGCCGGAAGCATAAAGTGTAAAGCCTGGGGTGCCTAATGAGTGAGCTAACTCACATTAATTGCGTTGCGCTCACTGCC<br>1041 CGCTTTCCAGTCGGGAAACCTGTCGTGCCAGCTGCATTAA |
| 17 | LET | **pComR-ST-XIP-ST**<br>*ComRSTO_XIPST_Peptide_Expression · 1070 bp · linear*<br>Suppl. Fig. 5d–e<br>Encoded peptide: **PYFAGCL**<br><br>**Annotated features**<br>lacZα (fragment) 67–135<br>ComR-ST promoter 251–354<br>pHP14 mRNA stability hairpin 359–399<br>BamHI-BglII Scar 400–405<br>RBS (strong) 406–419<br>Start codon (ATG) 420–422<br>XIP-ST Peptide 423–443<br>rrnB terminator (TrrnB) 453–820<br>lac operator 918–940 | 1 GAGAGTGCACCATATGCGGTGTGAAATACCGCACAGATGCGTAAGGAGAAAATACCGCATCAGGCGCCATTCGCCATTCA<br>81 GGCTGCGCAACTGTTGGGAAGGGCGATCGGTGCGGGCCTCTTCGCTATTACGCCAGCTGGCGAAAGGGGGATGTGCTGCA<br>161 AGGCGATTAAGTTGGGTAACGCCAGGGTTTTCCCAGTCACGACGTTGTAAAACGACGGCCAGTAAGATCTCGGCTGAATT<br>241 CTAAAGATCTGTGGTTTTGACTCTATCTCGCTTATTTAATTGTTTTAGTCAAGTTATTTATAGTGACATATATGTCTCTA<br>321 TTTTATTTTTCTACATTCTTTTATAATAGGTCTAGGCTACGTCGACTCTCGAGTGAGATTGTTGACGGTACCGTATTTTG<br>401 GATCTAGGAGGAAGGATCTATGCCATACTTCGCCGGCTGTCTGTAAGGATCTGAAGCTTGGGCCCGAACAAAAACTCATC<br>481 TCAGAAGAGGATCTGAATAGCGCCGTCGACCATCATCATCATCATCATTGAGTTTAAACGGTCTCCAGCTTGGCTGTTTT<br>561 GGCGGATGAGAGAAGATTTTCAGCCTGATACAGATTAAATCAGAACGCAGAAGCGGTCTGATAAAACAGAATTTGCCTGG<br>641 CGGCAGTAGCGCGGTGGTCCCACCTGACCCCATGCCGAACTCAGAAGTGAAACGCCGTAGCGCCGATGGTAGTGTGGGGT<br>721 CTCCCCATGCGAGAGTAGGGAACTGCCAGGCATCAAATAAAACGAAAGGCTCAGTCGAAAGACTGGGCCTTTCGTTTTAT<br>801 CTGTTGTTTGTCGGTGAACTCTGAAAGGAGGAACTATATCCGGATCGAGATCCTCTAGAGTCGACCTGCAGGCATGCAAG<br>881 CTTGGCGTAATCATGGTCATAGCTGTTTCCTGTGTGAAATTGTTATCCGCTCACAATTCCACACAACATACGAGCCGGAA<br>961 GCATAAAGTGTAAAGCCTGGGGTGCCTAATGAGTGAGCTAACTCACATTAATTGCGTTGCGCTCACTGCCCGCTTTCCAG<br>1041 TCGGGAAACCTGTCGTGCCAGCTGCATTAA |

| # | Type | Name and annotated features | Sequence (5′→3′) |
|---|---|---|---|
| 18 | LET | **pComR-SI-XIP-SI**<br>*ComRSTI_XIPSI_Peptide_Expression · 1072 bp · linear*<br>Suppl. Fig. 5d, 5f<br>Encoded peptide: **LLTAWWGL**<br><br>**Annotated features**<br>lacZα (fragment) 67–135<br>ComR-SI promoter 251–353<br>pHP14 mRNA stability hairpin 358–398<br>BamHI-BglII Scar 399–404<br>RBS (strong) 405–418<br>Start codon (ATG) 419–421<br>XIP-SI Peptide 422–445<br>rrnB terminator (TrrnB) 455–822<br>lac operator 920–942 | 1 GAGAGTGCACCATATGCGGTGTGAAATACCGCACAGATGCGTAAGGAGAAAATACCGCATCAGGCGCCATTCGCCATTCA<br>81 GGCTGCGCAACTGTTGGGAAGGGCGATCGGTGCGGGCCTCTTCGCTATTACGCCAGCTGGCGAAAGGGGGATGTGCTGCA<br>161 AGGCGATTAAGTTGGGTAACGCCAGGGTTTTCCCAGTCACGACGTTGTAAAACGACGGCCAGTAAGATCTCGGCTGAATT<br>241 CTAAAGATCTAAGCGGTAGACGGAGAGCAACTCAGATGAGTTGCTTTTTTTGTGCATCAATTTGGACATTTTTGTCATGT<br>321 TCCTTTTCCCTAATCCTTTCTATAATAAGGGTAGGCTACGTCGACTCTCGAGTGAGATTGTTGACGGTACCGTATTTTGG<br>401 ATCTAGGAGGAAGGATCTATGCTCCTCACAGCGTGGTGGGGCCTATAAGGATCTGAAGCTTGGGCCCGAACAAAAACTCA<br>481 TCTCAGAAGAGGATCTGAATAGCGCCGTCGACCATCATCATCATCATCATTGAGTTTAAACGGTCTCCAGCTTGGCTGTT<br>561 TTGGCGGATGAGAGAAGATTTTCAGCCTGATACAGATTAAATCAGAACGCAGAAGCGGTCTGATAAAACAGAATTTGCCT<br>641 GGCGGCAGTAGCGCGGTGGTCCCACCTGACCCCATGCCGAACTCAGAAGTGAAACGCCGTAGCGCCGATGGTAGTGTGGG<br>721 GTCTCCCCATGCGAGAGTAGGGAACTGCCAGGCATCAAATAAAACGAAAGGCTCAGTCGAAAGACTGGGCCTTTCGTTTT<br>801 ATCTGTTGTTTGTCGGTGAACTCTGAAAGGAGGAACTATATCCGGATCGAGATCCTCTAGAGTCGACCTGCAGGCATGCA<br>881 AGCTTGGCGTAATCATGGTCATAGCTGTTTCCTGTGTGAAATTGTTATCCGCTCACAATTCCACACAACATACGAGCCGG<br>961 AAGCATAAAGTGTAAAGCCTGGGGTGCCTAATGAGTGAGCTAACTCACATTAATTGCGTTGCGCTCACTGCCCGCTTTCC<br>1041 AGTCGGGAAACCTGTCGTGCCAGCTGCATTAA |
| **E. SHP3 alanine-scan peptide expression templates (pRgg3-SHP3 variants)** | | | |
| 19 | LET | **pRgg3-SHP3(D2A)**<br>*Ala1_Rgg3O_SHP3_Peptide_Expression · 1080 bp · linear*<br>Fig. 4a; Suppl. Figs 10–11<br>Encoded peptide: **AIIIVGG**<br><br>**Annotated features**<br>lacZα (fragment) 67–135<br>Rgg3 promoter 251–361<br>Rgg binding site 304–328<br>-10 345–350<br>Transcription start site 358–358<br>pHP14 mRNA stability hairpin 366–406<br>BamHI-BglII Scar 407–412<br>RBS (strong) 413–426<br>Start codon (ATG) 427–429<br>D2A 430–432<br>SHP3 Cognate Peptide 430–453<br>rrnB terminator (TrrnB) 463–830<br>lac operator 928–950 | 1 GAGAGTGCACCATATGCGGTGTGAAATACCGCACAGATGCGTAAGGAGAAAATACCGCATCAGGCGCCATTCGCCATTCA<br>81 GGCTGCGCAACTGTTGGGAAGGGCGATCGGTGCGGGCCTCTTCGCTATTACGCCAGCTGGCGAAAGGGGGATGTGCTGCA<br>161 AGGCGATTAAGTTGGGTAACGCCAGGGTTTTCCCAGTCACGACGTTGTAAAACGACGGCCAGTAAGATCTCGGCTGAATT<br>241 CTAAAGATCTTATGGCTTTCCTTTTCGAACTTTTCTAAGTGTGACACCAAGTTTTGATTTCATATTTTCCCACTTTCCCA<br>321 ACAAAAATCAATTCCGTTTTTTTTTATTGTACAATAAAACGGGCTACGTCGACTCTCGAGTGAGATTGTTGACGGTACCG<br>401 TATTTTGGATCTAGGAGGAAGGATCTATGGCGATCATTATTATCGTGGGTGGTTAAGGATCTGAAGCTTGGGCCCGAACA<br>481 AAAACTCATCTCAGAAGAGGATCTGAATAGCGCCGTCGACCATCATCATCATCATCATTGAGTTTAAACGGTCTCCAGCT<br>561 TGGCTGTTTTGGCGGATGAGAGAAGATTTTCAGCCTGATACAGATTAAATCAGAACGCAGAAGCGGTCTGATAAAACAGA<br>641 ATTTGCCTGGCGGCAGTAGCGCGGTGGTCCCACCTGACCCCATGCCGAACTCAGAAGTGAAACGCCGTAGCGCCGATGGT<br>721 AGTGTGGGGTCTCCCCATGCGAGAGTAGGGAACTGCCAGGCATCAAATAAAACGAAAGGCTCAGTCGAAAGACTGGGCCT<br>801 TTCGTTTTATCTGTTGTTTGTCGGTGAACTCTGAAAGGAGGAACTATATCCGGATCGAGATCCTCTAGAGTCGACCTGCA<br>881 GGCATGCAAGCTTGGCGTAATCATGGTCATAGCTGTTTCCTGTGTGAAATTGTTATCCGCTCACAATTCCACACAACATA<br>961 CGAGCCGGAAGCATAAAGTGTAAAGCCTGGGGTGCCTAATGAGTGAGCTAACTCACATTAATTGCGTTGCGCTCACTGCC<br>1041 CGCTTTCCAGTCGGGAAACCTGTCGTGCCAGCTGCATTAA |

| # | Type | Name and annotated features | Sequence (5′→3′) |
|---|---|---|---|
| 20 | LET | **pRgg3-SHP3(I3A)**<br>*Ala2_Rgg3O_SHP3_Peptide_Expression · 1080 bp · linear*<br>Fig. 4a; Suppl. Figs 10–11<br>Encoded peptide: **DAIIIVGG**<br><br>**Annotated features**<br>lacZα (fragment) 67–135<br>Rgg3 promoter 251–361<br>Rgg binding site 304–328<br>-10 345–350<br>Transcription start site 358–358<br>pHP14 mRNA stability hairpin 366–406<br>BamHI-BglII Scar 407–412<br>RBS (strong) 413–426<br>Start codon (ATG) 427–429<br>SHP3 Cognate Peptide 430–453<br>I3A 433–435<br>rrnB terminator (TrrnB) 463–830<br>lac operator 928–950 | 1 GAGAGTGCACCATATGCGGTGTGAAATACCGCACAGATGCGTAAGGAGAAAATACCGCATCAGGCGCCATTCGCCATTCA<br>81 GGCTGCGCAACTGTTGGGAAGGGCGATCGGTGCGGGCCTCTTCGCTATTACGCCAGCTGGCGAAAGGGGGATGTGCTGCA<br>161 AGGCGATTAAGTTGGGTAACGCCAGGGTTTTCCCAGTCACGACGTTGTAAAACGACGGCCAGTAAGATCTCGGCTGAATT<br>241 CTAAAGATCTTATGGCTTTCCTTTTCGAACTTTTCTAAGTGTGACACCAAGTTTTGATTTCATATTTTCCCACTTTCCCA<br>321 ACAAAAATCAATTCCGTTTTTTTTTATTGTACAATAAAACGGGCTACGTCGACTCTCGAGTGAGATTGTTGACGGTACCG<br>401 TATTTTGGATCTAGGAGGAAGGATCTATGGATGCGATTATTATCGTGGGTGGTTAAGGATCTGAAGCTTGGGCCCGAACA<br>481 AAAACTCATCTCAGAAGAGGATCTGAATAGCGCCGTCGACCATCATCATCATCATCATTGAGTTTAAACGGTCTCCAGCT<br>561 TGGCTGTTTTGGCGGATGAGAGAAGATTTTCAGCCTGATACAGATTAAATCAGAACGCAGAAGCGGTCTGATAAAACAGA<br>641 ATTTGCCTGGCGGCAGTAGCGCGGTGGTCCCACCTGACCCCATGCCGAACTCAGAAGTGAAACGCCGTAGCGCCGATGGT<br>721 AGTGTGGGGTCTCCCCATGCGAGAGTAGGGAACTGCCAGGCATCAAATAAAACGAAAGGCTCAGTCGAAAGACTGGGCCT<br>801 TTCGTTTTATCTGTTGTTTGTCGGTGAACTCTGAAAGGAGGAACTATATCCGGATCGAGATCCTCTAGAGTCGACCTGCA<br>881 GGCATGCAAGCTTGGCGTAATCATGGTCATAGCTGTTTCCTGTGTGAAATTGTTATCCGCTCACAATTCCACACAACATA<br>961 CGAGCCGGAAGCATAAAGTGTAAAGCCTGGGGTGCCTAATGAGTGAGCTAACTCACATTAATTGCGTTGCGCTCACTGCC<br>1041 CGCTTTCCAGTCGGGAAACCTGTCGTGCCAGCTGCATTAA |
| 21 | LET | **pRgg3-SHP3(I4A)**<br>*Ala3_Rgg3O_SHP3_Peptide_Expression · 1080 bp · linear*<br>Fig. 4a; Suppl. Figs 10–11<br>Encoded peptide: **DIAIIVGG**<br><br>**Annotated features**<br>lacZα (fragment) 67–135<br>Rgg3 promoter 251–361<br>Rgg binding site 304–328<br>-10 345–350<br>Transcription start site 358–358<br>pHP14 mRNA stability hairpin 366–406<br>BamHI-BglII Scar 407–412<br>RBS (strong) 413–426<br>Start codon (ATG) 427–429<br>SHP3 Cognate Peptide 430–453<br>I4A 436–438<br>rrnB terminator (TrrnB) 463–830<br>lac operator 928–950 | 1 GAGAGTGCACCATATGCGGTGTGAAATACCGCACAGATGCGTAAGGAGAAAATACCGCATCAGGCGCCATTCGCCATTCA<br>81 GGCTGCGCAACTGTTGGGAAGGGCGATCGGTGCGGGCCTCTTCGCTATTACGCCAGCTGGCGAAAGGGGGATGTGCTGCA<br>161 AGGCGATTAAGTTGGGTAACGCCAGGGTTTTCCCAGTCACGACGTTGTAAAACGACGGCCAGTAAGATCTCGGCTGAATT<br>241 CTAAAGATCTTATGGCTTTCCTTTTCGAACTTTTCTAAGTGTGACACCAAGTTTTGATTTCATATTTTCCCACTTTCCCA<br>321 ACAAAAATCAATTCCGTTTTTTTTTATTGTACAATAAAACGGGCTACGTCGACTCTCGAGTGAGATTGTTGACGGTACCG<br>401 TATTTTGGATCTAGGAGGAAGGATCTATGGATATCGCGATTATCGTGGGTGGTTAAGGATCTGAAGCTTGGGCCCGAACA<br>481 AAAACTCATCTCAGAAGAGGATCTGAATAGCGCCGTCGACCATCATCATCATCATCATTGAGTTTAAACGGTCTCCAGCT<br>561 TGGCTGTTTTGGCGGATGAGAGAAGATTTTCAGCCTGATACAGATTAAATCAGAACGCAGAAGCGGTCTGATAAAACAGA<br>641 ATTTGCCTGGCGGCAGTAGCGCGGTGGTCCCACCTGACCCCATGCCGAACTCAGAAGTGAAACGCCGTAGCGCCGATGGT<br>721 AGTGTGGGGTCTCCCCATGCGAGAGTAGGGAACTGCCAGGCATCAAATAAAACGAAAGGCTCAGTCGAAAGACTGGGCCT<br>801 TTCGTTTTATCTGTTGTTTGTCGGTGAACTCTGAAAGGAGGAACTATATCCGGATCGAGATCCTCTAGAGTCGACCTGCA<br>881 GGCATGCAAGCTTGGCGTAATCATGGTCATAGCTGTTTCCTGTGTGAAATTGTTATCCGCTCACAATTCCACACAACATA<br>961 CGAGCCGGAAGCATAAAGTGTAAAGCCTGGGGTGCCTAATGAGTGAGCTAACTCACATTAATTGCGTTGCGCTCACTGCC<br>1041 CGCTTTCCAGTCGGGAAACCTGTCGTGCCAGCTGCATTAA |

| # | Type | Name and annotated features | Sequence (5′→3′) |
|---|---|---|---|
| 22 | LET | **pRgg3-SHP3(I5A)**<br>*Ala4_Rgg3O_SHP3_Peptide_Expression · 1080 bp · linear*<br>Fig. 4a; Suppl. Figs 10–11<br>Encoded peptide: **DIIAIVGG**<br><br>**Annotated features**<br>lacZα (fragment) 67–135<br>Rgg3 promoter 251–361<br>Rgg binding site 304–328<br>-10 345–350<br>Transcription start site 358–358<br>pHP14 mRNA stability hairpin 366–406<br>BamHI-BglII Scar 407–412<br>RBS (strong) 413–426<br>Start codon (ATG) 427–429<br>SHP3 Cognate Peptide 430–453<br>I5A 439–441<br>rrnB terminator (TrrnB) 463–830<br>lac operator 928–950 | 1 GAGAGTGCACCATATGCGGTGTGAAATACCGCACAGATGCGTAAGGAGAAAATACCGCATCAGGCGCCATTCGCCATTCA<br>81 GGCTGCGCAACTGTTGGGAAGGGCGATCGGTGCGGGCCTCTTCGCTATTACGCCAGCTGGCGAAAGGGGGATGTGCTGCA<br>161 AGGCGATTAAGTTGGGTAACGCCAGGGTTTTCCCAGTCACGACGTTGTAAAACGACGGCCAGTAAGATCTCGGCTGAATT<br>241 CTAAAGATCTTATGGCTTTCCTTTTCGAACTTTTCTAAGTGTGACACCAAGTTTTGATTTCATATTTTCCCACTTTCCCA<br>321 ACAAAAATCAATTCCGTTTTTTTTTATTGTACAATAAAACGGGCTACGTCGACTCTCGAGTGAGATTGTTGACGGTACCG<br>401 TATTTTGGATCTAGGAGGAAGGATCTATGGATATCATTGCGATCGTGGGTGGTTAAGGATCTGAAGCTTGGGCCCGAACA<br>481 AAAACTCATCTCAGAAGAGGATCTGAATAGCGCCGTCGACCATCATCATCATCATCATTGAGTTTAAACGGTCTCCAGCT<br>561 TGGCTGTTTTGGCGGATGAGAGAAGATTTTCAGCCTGATACAGATTAAATCAGAACGCAGAAGCGGTCTGATAAAACAGA<br>641 ATTTGCCTGGCGGCAGTAGCGCGGTGGTCCCACCTGACCCCATGCCGAACTCAGAAGTGAAACGCCGTAGCGCCGATGGT<br>721 AGTGTGGGGTCTCCCCATGCGAGAGTAGGGAACTGCCAGGCATCAAATAAAACGAAAGGCTCAGTCGAAAGACTGGGCCT<br>801 TTCGTTTTATCTGTTGTTTGTCGGTGAACTCTGAAAGGAGGAACTATATCCGGATCGAGATCCTCTAGAGTCGACCTGCA<br>881 GGCATGCAAGCTTGGCGTAATCATGGTCATAGCTGTTTCCTGTGTGAAATTGTTATCCGCTCACAATTCCACACAACATA<br>961 CGAGCCGGAAGCATAAAGTGTAAAGCCTGGGGTGCCTAATGAGTGAGCTAACTCACATTAATTGCGTTGCGCTCACTGCC<br>1041 CGCTTTCCAGTCGGGAAACCTGTCGTGCCAGCTGCATTAA |
| 23 | LET | **pRgg3-SHP3(I6A)**<br>*Ala5_Rgg3O_SHP3_Peptide_Expression · 1080 bp · linear*<br>Fig. 4a; Suppl. Figs 10–11<br>Encoded peptide: **DIIIAVGG**<br><br>**Annotated features**<br>lacZα (fragment) 67–135<br>Rgg3 promoter 251–361<br>Rgg binding site 304–328<br>-10 345–350<br>Transcription start site 358–358<br>pHP14 mRNA stability hairpin 366–406<br>BamHI-BglII Scar 407–412<br>RBS (strong) 413–426<br>Start codon (ATG) 427–429<br>SHP3 Cognate Peptide 430–453<br>I6A 442–444<br>rrnB terminator (TrrnB) 463–830<br>lac operator 928–950 | 1 GAGAGTGCACCATATGCGGTGTGAAATACCGCACAGATGCGTAAGGAGAAAATACCGCATCAGGCGCCATTCGCCATTCA<br>81 GGCTGCGCAACTGTTGGGAAGGGCGATCGGTGCGGGCCTCTTCGCTATTACGCCAGCTGGCGAAAGGGGGATGTGCTGCA<br>161 AGGCGATTAAGTTGGGTAACGCCAGGGTTTTCCCAGTCACGACGTTGTAAAACGACGGCCAGTAAGATCTCGGCTGAATT<br>241 CTAAAGATCTTATGGCTTTCCTTTTCGAACTTTTCTAAGTGTGACACCAAGTTTTGATTTCATATTTTCCCACTTTCCCA<br>321 ACAAAAATCAATTCCGTTTTTTTTTATTGTACAATAAAACGGGCTACGTCGACTCTCGAGTGAGATTGTTGACGGTACCG<br>401 TATTTTGGATCTAGGAGGAAGGATCTATGGATATCATTATTGCGGTGGGTGGTTAAGGATCTGAAGCTTGGGCCCGAACA<br>481 AAAACTCATCTCAGAAGAGGATCTGAATAGCGCCGTCGACCATCATCATCATCATCATTGAGTTTAAACGGTCTCCAGCT<br>561 TGGCTGTTTTGGCGGATGAGAGAAGATTTTCAGCCTGATACAGATTAAATCAGAACGCAGAAGCGGTCTGATAAAACAGA<br>641 ATTTGCCTGGCGGCAGTAGCGCGGTGGTCCCACCTGACCCCATGCCGAACTCAGAAGTGAAACGCCGTAGCGCCGATGGT<br>721 AGTGTGGGGTCTCCCCATGCGAGAGTAGGGAACTGCCAGGCATCAAATAAAACGAAAGGCTCAGTCGAAAGACTGGGCCT<br>801 TTCGTTTTATCTGTTGTTTGTCGGTGAACTCTGAAAGGAGGAACTATATCCGGATCGAGATCCTCTAGAGTCGACCTGCA<br>881 GGCATGCAAGCTTGGCGTAATCATGGTCATAGCTGTTTCCTGTGTGAAATTGTTATCCGCTCACAATTCCACACAACATA<br>961 CGAGCCGGAAGCATAAAGTGTAAAGCCTGGGGTGCCTAATGAGTGAGCTAACTCACATTAATTGCGTTGCGCTCACTGCC<br>1041 CGCTTTCCAGTCGGGAAACCTGTCGTGCCAGCTGCATTAA |

| # | Type | Name and annotated features | Sequence (5′→3′) |
|---|---|---|---|
| 24 | LET | **pRgg3-SHP3(V7A)**<br>*Ala6_Rgg3O_SHP3_Peptide_Expression · 1080 bp · linear*<br>Fig. 4a; Suppl. Figs 10–11<br>Encoded peptide: **DIIIIAGG**<br><br>**Annotated features**<br>lacZα (fragment) 67–135<br>Rgg3 promoter 251–361<br>Rgg binding site 304–328<br>-10 345–350<br>Transcription start site 358–358<br>pHP14 mRNA stability hairpin 366–406<br>BamHI-BglII Scar 407–412<br>RBS (strong) 413–426<br>Start codon (ATG) 427–429<br>SHP3 Cognate Peptide 430–453<br>V7A 445–447<br>rrnB terminator (TrrnB) 463–830<br>lac operator 928–950 | 1 GAGAGTGCACCATATGCGGTGTGAAATACCGCACAGATGCGTAAGGAGAAAATACCGCATCAGGCGCCATTCGCCATTCA<br>81 GGCTGCGCAACTGTTGGGAAGGGCGATCGGTGCGGGCCTCTTCGCTATTACGCCAGCTGGCGAAAGGGGGATGTGCTGCA<br>161 AGGCGATTAAGTTGGGTAACGCCAGGGTTTTCCCAGTCACGACGTTGTAAAACGACGGCCAGTAAGATCTCGGCTGAATT<br>241 CTAAAGATCTTATGGCTTTCCTTTTCGAACTTTTCTAAGTGTGACACCAAGTTTTGATTTCATATTTTCCCACTTTCCCA<br>321 ACAAAAATCAATTCCGTTTTTTTTTATTGTACAATAAAACGGGCTACGTCGACTCTCGAGTGAGATTGTTGACGGTACCG<br>401 TATTTTGGATCTAGGAGGAAGGATCTATGGATATCATTATTATCGCGGGTGGTTAAGGATCTGAAGCTTGGGCCCGAACA<br>481 AAAACTCATCTCAGAAGAGGATCTGAATAGCGCCGTCGACCATCATCATCATCATCATTGAGTTTAAACGGTCTCCAGCT<br>561 TGGCTGTTTTGGCGGATGAGAGAAGATTTTCAGCCTGATACAGATTAAATCAGAACGCAGAAGCGGTCTGATAAAACAGA<br>641 ATTTGCCTGGCGGCAGTAGCGCGGTGGTCCCACCTGACCCCATGCCGAACTCAGAAGTGAAACGCCGTAGCGCCGATGGT<br>721 AGTGTGGGGTCTCCCCATGCGAGAGTAGGGAACTGCCAGGCATCAAATAAAACGAAAGGCTCAGTCGAAAGACTGGGCCT<br>801 TTCGTTTTATCTGTTGTTTGTCGGTGAACTCTGAAAGGAGGAACTATATCCGGATCGAGATCCTCTAGAGTCGACCTGCA<br>881 GGCATGCAAGCTTGGCGTAATCATGGTCATAGCTGTTTCCTGTGTGAAATTGTTATCCGCTCACAATTCCACACAACATA<br>961 CGAGCCGGAAGCATAAAGTGTAAAGCCTGGGGTGCCTAATGAGTGAGCTAACTCACATTAATTGCGTTGCGCTCACTGCC<br>1041 CGCTTTCCAGTCGGGAAACCTGTCGTGCCAGCTGCATTAA |
| 25 | LET | **pRgg3-SHP3(G8A)**<br>*Ala7_Rgg3O_SHP3_Peptide_Expression · 1080 bp · linear*<br>Fig. 4a; Suppl. Figs 10–11<br>Encoded peptide: **DIIIIVAG**<br><br>**Annotated features**<br>lacZα (fragment) 67–135<br>Rgg3 promoter 251–361<br>Rgg binding site 304–328<br>-10 345–350<br>Transcription start site 358–358<br>pHP14 mRNA stability hairpin 366–406<br>BamHI-BglII Scar 407–412<br>RBS (strong) 413–426<br>Start codon (ATG) 427–429<br>SHP3 Cognate Peptide 430–453<br>G8A 448–450<br>rrnB terminator (TrrnB) 463–830<br>lac operator 928–950 | 1 GAGAGTGCACCATATGCGGTGTGAAATACCGCACAGATGCGTAAGGAGAAAATACCGCATCAGGCGCCATTCGCCATTCA<br>81 GGCTGCGCAACTGTTGGGAAGGGCGATCGGTGCGGGCCTCTTCGCTATTACGCCAGCTGGCGAAAGGGGGATGTGCTGCA<br>161 AGGCGATTAAGTTGGGTAACGCCAGGGTTTTCCCAGTCACGACGTTGTAAAACGACGGCCAGTAAGATCTCGGCTGAATT<br>241 CTAAAGATCTTATGGCTTTCCTTTTCGAACTTTTCTAAGTGTGACACCAAGTTTTGATTTCATATTTTCCCACTTTCCCA<br>321 ACAAAAATCAATTCCGTTTTTTTTTATTGTACAATAAAACGGGCTACGTCGACTCTCGAGTGAGATTGTTGACGGTACCG<br>401 TATTTTGGATCTAGGAGGAAGGATCTATGGATATCATTATTATCGTGGCGGGTTAAGGATCTGAAGCTTGGGCCCGAACA<br>481 AAAACTCATCTCAGAAGAGGATCTGAATAGCGCCGTCGACCATCATCATCATCATCATTGAGTTTAAACGGTCTCCAGCT<br>561 TGGCTGTTTTGGCGGATGAGAGAAGATTTTCAGCCTGATACAGATTAAATCAGAACGCAGAAGCGGTCTGATAAAACAGA<br>641 ATTTGCCTGGCGGCAGTAGCGCGGTGGTCCCACCTGACCCCATGCCGAACTCAGAAGTGAAACGCCGTAGCGCCGATGGT<br>721 AGTGTGGGGTCTCCCCATGCGAGAGTAGGGAACTGCCAGGCATCAAATAAAACGAAAGGCTCAGTCGAAAGACTGGGCCT<br>801 TTCGTTTTATCTGTTGTTTGTCGGTGAACTCTGAAAGGAGGAACTATATCCGGATCGAGATCCTCTAGAGTCGACCTGCA<br>881 GGCATGCAAGCTTGGCGTAATCATGGTCATAGCTGTTTCCTGTGTGAAATTGTTATCCGCTCACAATTCCACACAACATA<br>961 CGAGCCGGAAGCATAAAGTGTAAAGCCTGGGGTGCCTAATGAGTGAGCTAACTCACATTAATTGCGTTGCGCTCACTGCC<br>1041 CGCTTTCCAGTCGGGAAACCTGTCGTGCCAGCTGCATTAA |

| # | Type | Name and annotated features | Sequence (5′→3′) |
|---|---|---|---|
| 26 | LET | **pRgg3-SHP3(G9A)**<br>*Ala8_Rgg3O_SHP3_Peptide_Expression · 1080 bp · linear*<br>Fig. 4a; Suppl. Figs 10–11<br>Encoded peptide: **DIIIIVGA**<br><br>**Annotated features**<br>lacZα (fragment) 67–135<br>Rgg3 promoter 251–361<br>Rgg binding site 304–328<br>-10 345–350<br>Transcription start site 358–358<br>pHP14 mRNA stability hairpin 366–406<br>BamHI-BglII Scar 407–412<br>RBS (strong) 413–426<br>Start codon (ATG) 427–429<br>SHP3 Cognate Peptide 430–453<br>G9A 451–453<br>rrnB terminator (TrrnB) 463–830<br>lac operator 928–950 | 1 GAGAGTGCACCATATGCGGTGTGAAATACCGCACAGATGCGTAAGGAGAAAATACCGCATCAGGCGCCATTCGCCATTCA<br>81 GGCTGCGCAACTGTTGGGAAGGGCGATCGGTGCGGGCCTCTTCGCTATTACGCCAGCTGGCGAAAGGGGGATGTGCTGCA<br>161 AGGCGATTAAGTTGGGTAACGCCAGGGTTTTCCCAGTCACGACGTTGTAAAACGACGGCCAGTAAGATCTCGGCTGAATT<br>241 CTAAAGATCTTATGGCTTTCCTTTTCGAACTTTTCTAAGTGTGACACCAAGTTTTGATTTCATATTTTCCCACTTTCCCA<br>321 ACAAAAATCAATTCCGTTTTTTTTTATTGTACAATAAAACGGGCTACGTCGACTCTCGAGTGAGATTGTTGACGGTACCG<br>401 TATTTTGGATCTAGGAGGAAGGATCTATGGATATCATTATTATCGTGGGTGCGTAAGGATCTGAAGCTTGGGCCCGAACA<br>481 AAAACTCATCTCAGAAGAGGATCTGAATAGCGCCGTCGACCATCATCATCATCATCATTGAGTTTAAACGGTCTCCAGCT<br>561 TGGCTGTTTTGGCGGATGAGAGAAGATTTTCAGCCTGATACAGATTAAATCAGAACGCAGAAGCGGTCTGATAAAACAGA<br>641 ATTTGCCTGGCGGCAGTAGCGCGGTGGTCCCACCTGACCCCATGCCGAACTCAGAAGTGAAACGCCGTAGCGCCGATGGT<br>721 AGTGTGGGGTCTCCCCATGCGAGAGTAGGGAACTGCCAGGCATCAAATAAAACGAAAGGCTCAGTCGAAAGACTGGGCCT<br>801 TTCGTTTTATCTGTTGTTTGTCGGTGAACTCTGAAAGGAGGAACTATATCCGGATCGAGATCCTCTAGAGTCGACCTGCA<br>881 GGCATGCAAGCTTGGCGTAATCATGGTCATAGCTGTTTCCTGTGTGAAATTGTTATCCGCTCACAATTCCACACAACATA<br>961 CGAGCCGGAAGCATAAAGTGTAAAGCCTGGGGTGCCTAATGAGTGAGCTAACTCACATTAATTGCGTTGCGCTCACTGCC<br>1041 CGCTTTCCAGTCGGGAAACCTGTCGTGCCAGCTGCATTAA |
| **F. SHP3 site-saturation variant peptide expression templates (pRgg3-SHP3 variants)** | | | |
| 27 | LET | **pRgg3-SHP3(I3L)**<br>*1_SHP3_I3L · 1080 bp · linear*<br>Fig. 5a; Suppl. Fig. 12<br>Encoded peptide: **DLIIIVGG**<br><br>**Annotated features**<br>lacZα (fragment) 67–135<br>Rgg3 promoter 251–361<br>Rgg binding site 304–328<br>-10 345–350<br>Transcription start site 358–358<br>pHP14 mRNA stability hairpin 366–406<br>BamHI-BglII Scar 407–412<br>RBS (strong) 413–426<br>Start codon (ATG) 427–429<br>SHP3 Cognate Peptide 430–453<br>I3L 433–435<br>rrnB terminator (TrrnB) 463–830<br>lac operator 928–950 | 1 GAGAGTGCACCATATGCGGTGTGAAATACCGCACAGATGCGTAAGGAGAAAATACCGCATCAGGCGCCATTCGCCATTCA<br>81 GGCTGCGCAACTGTTGGGAAGGGCGATCGGTGCGGGCCTCTTCGCTATTACGCCAGCTGGCGAAAGGGGGATGTGCTGCA<br>161 AGGCGATTAAGTTGGGTAACGCCAGGGTTTTCCCAGTCACGACGTTGTAAAACGACGGCCAGTAAGATCTCGGCTGAATT<br>241 CTAAAGATCTTATGGCTTTCCTTTTCGAACTTTTCTAAGTGTGACACCAAGTTTTGATTTCATATTTTCCCACTTTCCCA<br>321 ACAAAAATCAATTCCGTTTTTTTTTATTGTACAATAAAACGGGCTACGTCGACTCTCGAGTGAGATTGTTGACGGTACCG<br>401 TATTTTGGATCTAGGAGGAAGGATCTATGGATTTAATTATTATCGTGGGTGGTTAAGGATCTGAAGCTTGGGCCCGAACA<br>481 AAAACTCATCTCAGAAGAGGATCTGAATAGCGCCGTCGACCATCATCATCATCATCATTGAGTTTAAACGGTCTCCAGCT<br>561 TGGCTGTTTTGGCGGATGAGAGAAGATTTTCAGCCTGATACAGATTAAATCAGAACGCAGAAGCGGTCTGATAAAACAGA<br>641 ATTTGCCTGGCGGCAGTAGCGCGGTGGTCCCACCTGACCCCATGCCGAACTCAGAAGTGAAACGCCGTAGCGCCGATGGT<br>721 AGTGTGGGGTCTCCCCATGCGAGAGTAGGGAACTGCCAGGCATCAAATAAAACGAAAGGCTCAGTCGAAAGACTGGGCCT<br>801 TTCGTTTTATCTGTTGTTTGTCGGTGAACTCTGAAAGGAGGAACTATATCCGGATCGAGATCCTCTAGAGTCGACCTGCA<br>881 GGCATGCAAGCTTGGCGTAATCATGGTCATAGCTGTTTCCTGTGTGAAATTGTTATCCGCTCACAATTCCACACAACATA<br>961 CGAGCCGGAAGCATAAAGTGTAAAGCCTGGGGTGCCTAATGAGTGAGCTAACTCACATTAATTGCGTTGCGCTCACTGCC<br>1041 CGCTTTCCAGTCGGGAAACCTGTCGTGCCAGCTGCATTAA |

| # | Type | Name and annotated features | Sequence (5′→3′) |
|---|---|---|---|
| 28 | LET | **pRgg3-SHP3(I4T)**<br>*2_SHP3_I4T · 1080 bp · linear*<br>Fig. 5a; Suppl. Fig. 12<br>Encoded peptide: **DITIIVGG**<br><br>**Annotated features**<br>lacZα (fragment) 67–135<br>Rgg3 promoter 251–361<br>Rgg binding site 304–328<br>-10 345–350<br>Transcription start site 358–358<br>pHP14 mRNA stability hairpin 366–406<br>BamHI-BglII Scar 407–412<br>RBS (strong) 413–426<br>Start codon (ATG) 427–429<br>SHP3 Cognate Peptide 430–453<br>I4T 436–438<br>rrnB terminator (TrrnB) 463–830<br>lac operator 928–950 | 1 GAGAGTGCACCATATGCGGTGTGAAATACCGCACAGATGCGTAAGGAGAAAATACCGCATCAGGCGCCATTCGCCATTCA<br>81 GGCTGCGCAACTGTTGGGAAGGGCGATCGGTGCGGGCCTCTTCGCTATTACGCCAGCTGGCGAAAGGGGGATGTGCTGCA<br>161 AGGCGATTAAGTTGGGTAACGCCAGGGTTTTCCCAGTCACGACGTTGTAAAACGACGGCCAGTAAGATCTCGGCTGAATT<br>241 CTAAAGATCTTATGGCTTTCCTTTTCGAACTTTTCTAAGTGTGACACCAAGTTTTGATTTCATATTTTCCCACTTTCCCA<br>321 ACAAAAATCAATTCCGTTTTTTTTTATTGTACAATAAAACGGGCTACGTCGACTCTCGAGTGAGATTGTTGACGGTACCG<br>401 TATTTTGGATCTAGGAGGAAGGATCTATGGATATCACCATTATCGTGGGTGGTTAAGGATCTGAAGCTTGGGCCCGAACA<br>481 AAAACTCATCTCAGAAGAGGATCTGAATAGCGCCGTCGACCATCATCATCATCATCATTGAGTTTAAACGGTCTCCAGCT<br>561 TGGCTGTTTTGGCGGATGAGAGAAGATTTTCAGCCTGATACAGATTAAATCAGAACGCAGAAGCGGTCTGATAAAACAGA<br>641 ATTTGCCTGGCGGCAGTAGCGCGGTGGTCCCACCTGACCCCATGCCGAACTCAGAAGTGAAACGCCGTAGCGCCGATGGT<br>721 AGTGTGGGGTCTCCCCATGCGAGAGTAGGGAACTGCCAGGCATCAAATAAAACGAAAGGCTCAGTCGAAAGACTGGGCCT<br>801 TTCGTTTTATCTGTTGTTTGTCGGTGAACTCTGAAAGGAGGAACTATATCCGGATCGAGATCCTCTAGAGTCGACCTGCA<br>881 GGCATGCAAGCTTGGCGTAATCATGGTCATAGCTGTTTCCTGTGTGAAATTGTTATCCGCTCACAATTCCACACAACATA<br>961 CGAGCCGGAAGCATAAAGTGTAAAGCCTGGGGTGCCTAATGAGTGAGCTAACTCACATTAATTGCGTTGCGCTCACTGCC<br>1041 CGCTTTCCAGTCGGGAAACCTGTCGTGCCAGCTGCATTAA |
| 29 | LET | **pRgg3-SHP3(I5V)**<br>*3_SHP3_I5V · 1080 bp · linear*<br>Fig. 5a; Suppl. Fig. 12<br>Encoded peptide: **DIIVIVGG**<br><br>**Annotated features**<br>lacZα (fragment) 67–135<br>Rgg3 promoter 251–361<br>Rgg binding site 304–328<br>-10 345–350<br>Transcription start site 358–358<br>pHP14 mRNA stability hairpin 366–406<br>BamHI-BglII Scar 407–412<br>RBS (strong) 413–426<br>Start codon (ATG) 427–429<br>SHP3 Cognate Peptide 430–453<br>I5V 439–441<br>rrnB terminator (TrrnB) 463–830<br>lac operator 928–950 | 1 GAGAGTGCACCATATGCGGTGTGAAATACCGCACAGATGCGTAAGGAGAAAATACCGCATCAGGCGCCATTCGCCATTCA<br>81 GGCTGCGCAACTGTTGGGAAGGGCGATCGGTGCGGGCCTCTTCGCTATTACGCCAGCTGGCGAAAGGGGGATGTGCTGCA<br>161 AGGCGATTAAGTTGGGTAACGCCAGGGTTTTCCCAGTCACGACGTTGTAAAACGACGGCCAGTAAGATCTCGGCTGAATT<br>241 CTAAAGATCTTATGGCTTTCCTTTTCGAACTTTTCTAAGTGTGACACCAAGTTTTGATTTCATATTTTCCCACTTTCCCA<br>321 ACAAAAATCAATTCCGTTTTTTTTTATTGTACAATAAAACGGGCTACGTCGACTCTCGAGTGAGATTGTTGACGGTACCG<br>401 TATTTTGGATCTAGGAGGAAGGATCTATGGATATCATTGTTATCGTGGGTGGTTAAGGATCTGAAGCTTGGGCCCGAACA<br>481 AAAACTCATCTCAGAAGAGGATCTGAATAGCGCCGTCGACCATCATCATCATCATCATTGAGTTTAAACGGTCTCCAGCT<br>561 TGGCTGTTTTGGCGGATGAGAGAAGATTTTCAGCCTGATACAGATTAAATCAGAACGCAGAAGCGGTCTGATAAAACAGA<br>641 ATTTGCCTGGCGGCAGTAGCGCGGTGGTCCCACCTGACCCCATGCCGAACTCAGAAGTGAAACGCCGTAGCGCCGATGGT<br>721 AGTGTGGGGTCTCCCCATGCGAGAGTAGGGAACTGCCAGGCATCAAATAAAACGAAAGGCTCAGTCGAAAGACTGGGCCT<br>801 TTCGTTTTATCTGTTGTTTGTCGGTGAACTCTGAAAGGAGGAACTATATCCGGATCGAGATCCTCTAGAGTCGACCTGCA<br>881 GGCATGCAAGCTTGGCGTAATCATGGTCATAGCTGTTTCCTGTGTGAAATTGTTATCCGCTCACAATTCCACACAACATA<br>961 CGAGCCGGAAGCATAAAGTGTAAAGCCTGGGGTGCCTAATGAGTGAGCTAACTCACATTAATTGCGTTGCGCTCACTGCC<br>1041 CGCTTTCCAGTCGGGAAACCTGTCGTGCCAGCTGCATTAA |

| # | Type | Name and annotated features | Sequence (5′→3′) |
|---|---|---|---|
| 30 | LET | **pRgg3-SHP3(I6L)**<br>*4_SHP3_I6L · 1080 bp · linear*<br>Fig. 5a; Suppl. Fig. 12<br>Encoded peptide: **DIIILVGG**<br><br>**Annotated features**<br>lacZα (fragment) 67–135<br>Rgg3 promoter 251–361<br>Rgg binding site 304–328<br>-10 345–350<br>Transcription start site 358–358<br>pHP14 mRNA stability hairpin 366–406<br>BamHI-BglII Scar 407–412<br>RBS (strong) 413–426<br>Start codon (ATG) 427–429<br>SHP3 Cognate Peptide 430–453<br>I6L 442–444<br>rrnB terminator (TrrnB) 463–830<br>lac operator 928–950 | 1 GAGAGTGCACCATATGCGGTGTGAAATACCGCACAGATGCGTAAGGAGAAAATACCGCATCAGGCGCCATTCGCCATTCA<br>81 GGCTGCGCAACTGTTGGGAAGGGCGATCGGTGCGGGCCTCTTCGCTATTACGCCAGCTGGCGAAAGGGGGATGTGCTGCA<br>161 AGGCGATTAAGTTGGGTAACGCCAGGGTTTTCCCAGTCACGACGTTGTAAAACGACGGCCAGTAAGATCTCGGCTGAATT<br>241 CTAAAGATCTTATGGCTTTCCTTTTCGAACTTTTCTAAGTGTGACACCAAGTTTTGATTTCATATTTTCCCACTTTCCCA<br>321 ACAAAAATCAATTCCGTTTTTTTTTATTGTACAATAAAACGGGCTACGTCGACTCTCGAGTGAGATTGTTGACGGTACCG<br>401 TATTTTGGATCTAGGAGGAAGGATCTATGGATATCATTATTTTAGTGGGTGGTTAAGGATCTGAAGCTTGGGCCCGAACA<br>481 AAAACTCATCTCAGAAGAGGATCTGAATAGCGCCGTCGACCATCATCATCATCATCATTGAGTTTAAACGGTCTCCAGCT<br>561 TGGCTGTTTTGGCGGATGAGAGAAGATTTTCAGCCTGATACAGATTAAATCAGAACGCAGAAGCGGTCTGATAAAACAGA<br>641 ATTTGCCTGGCGGCAGTAGCGCGGTGGTCCCACCTGACCCCATGCCGAACTCAGAAGTGAAACGCCGTAGCGCCGATGGT<br>721 AGTGTGGGGTCTCCCCATGCGAGAGTAGGGAACTGCCAGGCATCAAATAAAACGAAAGGCTCAGTCGAAAGACTGGGCCT<br>801 TTCGTTTTATCTGTTGTTTGTCGGTGAACTCTGAAAGGAGGAACTATATCCGGATCGAGATCCTCTAGAGTCGACCTGCA<br>881 GGCATGCAAGCTTGGCGTAATCATGGTCATAGCTGTTTCCTGTGTGAAATTGTTATCCGCTCACAATTCCACACAACATA<br>961 CGAGCCGGAAGCATAAAGTGTAAAGCCTGGGGTGCCTAATGAGTGAGCTAACTCACATTAATTGCGTTGCGCTCACTGCC<br>1041 CGCTTTCCAGTCGGGAAACCTGTCGTGCCAGCTGCATTAA |
| 31 | LET | **pRgg3-SHP3(V7T)**<br>*5_SHP3_V7T · 1080 bp · linear*<br>Fig. 5a; Suppl. Fig. 12<br>Encoded peptide: **DIIIITGG**<br><br>**Annotated features**<br>lacZα (fragment) 67–135<br>Rgg3 promoter 251–361<br>Rgg binding site 304–328<br>-10 345–350<br>Transcription start site 358–358<br>pHP14 mRNA stability hairpin 366–406<br>BamHI-BglII Scar 407–412<br>RBS (strong) 413–426<br>Start codon (ATG) 427–429<br>SHP3 Cognate Peptide 430–453<br>V7T 445–447<br>rrnB terminator (TrrnB) 463–830<br>lac operator 928–950 | 1 GAGAGTGCACCATATGCGGTGTGAAATACCGCACAGATGCGTAAGGAGAAAATACCGCATCAGGCGCCATTCGCCATTCA<br>81 GGCTGCGCAACTGTTGGGAAGGGCGATCGGTGCGGGCCTCTTCGCTATTACGCCAGCTGGCGAAAGGGGGATGTGCTGCA<br>161 AGGCGATTAAGTTGGGTAACGCCAGGGTTTTCCCAGTCACGACGTTGTAAAACGACGGCCAGTAAGATCTCGGCTGAATT<br>241 CTAAAGATCTTATGGCTTTCCTTTTCGAACTTTTCTAAGTGTGACACCAAGTTTTGATTTCATATTTTCCCACTTTCCCA<br>321 ACAAAAATCAATTCCGTTTTTTTTTATTGTACAATAAAACGGGCTACGTCGACTCTCGAGTGAGATTGTTGACGGTACCG<br>401 TATTTTGGATCTAGGAGGAAGGATCTATGGATATCATTATTATCACCGGTGGTTAAGGATCTGAAGCTTGGGCCCGAACA<br>481 AAAACTCATCTCAGAAGAGGATCTGAATAGCGCCGTCGACCATCATCATCATCATCATTGAGTTTAAACGGTCTCCAGCT<br>561 TGGCTGTTTTGGCGGATGAGAGAAGATTTTCAGCCTGATACAGATTAAATCAGAACGCAGAAGCGGTCTGATAAAACAGA<br>641 ATTTGCCTGGCGGCAGTAGCGCGGTGGTCCCACCTGACCCCATGCCGAACTCAGAAGTGAAACGCCGTAGCGCCGATGGT<br>721 AGTGTGGGGTCTCCCCATGCGAGAGTAGGGAACTGCCAGGCATCAAATAAAACGAAAGGCTCAGTCGAAAGACTGGGCCT<br>801 TTCGTTTTATCTGTTGTTTGTCGGTGAACTCTGAAAGGAGGAACTATATCCGGATCGAGATCCTCTAGAGTCGACCTGCA<br>881 GGCATGCAAGCTTGGCGTAATCATGGTCATAGCTGTTTCCTGTGTGAAATTGTTATCCGCTCACAATTCCACACAACATA<br>961 CGAGCCGGAAGCATAAAGTGTAAAGCCTGGGGTGCCTAATGAGTGAGCTAACTCACATTAATTGCGTTGCGCTCACTGCC<br>1041 CGCTTTCCAGTCGGGAAACCTGTCGTGCCAGCTGCATTAA |

| # | Type | Name and annotated features | Sequence (5′→3′) |
|---|---|---|---|
| 32 | LET | **pRgg3-SHP3(G8T)**<br>*6_SHP3_G8T · 1080 bp · linear*<br>Fig. 5a; Suppl. Fig. 12<br>Encoded peptide: **DIIIIVTG**<br><br>**Annotated features**<br>lacZα (fragment) 67–135<br>Rgg3 promoter 251–361<br>Rgg binding site 304–328<br>-10 345–350<br>Transcription start site 358–358<br>pHP14 mRNA stability hairpin 366–406<br>BamHI-BglII Scar 407–412<br>RBS (strong) 413–426<br>Start codon (ATG) 427–429<br>SHP3 Cognate Peptide 430–453<br>G8T 448–450<br>rrnB terminator (TrrnB) 463–830<br>lac operator 928–950 | 1 GAGAGTGCACCATATGCGGTGTGAAATACCGCACAGATGCGTAAGGAGAAAATACCGCATCAGGCGCCATTCGCCATTCA<br>81 GGCTGCGCAACTGTTGGGAAGGGCGATCGGTGCGGGCCTCTTCGCTATTACGCCAGCTGGCGAAAGGGGGATGTGCTGCA<br>161 AGGCGATTAAGTTGGGTAACGCCAGGGTTTTCCCAGTCACGACGTTGTAAAACGACGGCCAGTAAGATCTCGGCTGAATT<br>241 CTAAAGATCTTATGGCTTTCCTTTTCGAACTTTTCTAAGTGTGACACCAAGTTTTGATTTCATATTTTCCCACTTTCCCA<br>321 ACAAAAATCAATTCCGTTTTTTTTTATTGTACAATAAAACGGGCTACGTCGACTCTCGAGTGAGATTGTTGACGGTACCG<br>401 TATTTTGGATCTAGGAGGAAGGATCTATGGATATCATTATTATCGTGACCGGTTAAGGATCTGAAGCTTGGGCCCGAACA<br>481 AAAACTCATCTCAGAAGAGGATCTGAATAGCGCCGTCGACCATCATCATCATCATCATTGAGTTTAAACGGTCTCCAGCT<br>561 TGGCTGTTTTGGCGGATGAGAGAAGATTTTCAGCCTGATACAGATTAAATCAGAACGCAGAAGCGGTCTGATAAAACAGA<br>641 ATTTGCCTGGCGGCAGTAGCGCGGTGGTCCCACCTGACCCCATGCCGAACTCAGAAGTGAAACGCCGTAGCGCCGATGGT<br>721 AGTGTGGGGTCTCCCCATGCGAGAGTAGGGAACTGCCAGGCATCAAATAAAACGAAAGGCTCAGTCGAAAGACTGGGCCT<br>801 TTCGTTTTATCTGTTGTTTGTCGGTGAACTCTGAAAGGAGGAACTATATCCGGATCGAGATCCTCTAGAGTCGACCTGCA<br>881 GGCATGCAAGCTTGGCGTAATCATGGTCATAGCTGTTTCCTGTGTGAAATTGTTATCCGCTCACAATTCCACACAACATA<br>961 CGAGCCGGAAGCATAAAGTGTAAAGCCTGGGGTGCCTAATGAGTGAGCTAACTCACATTAATTGCGTTGCGCTCACTGCC<br>1041 CGCTTTCCAGTCGGGAAACCTGTCGTGCCAGCTGCATTAA |
| 33 | LET | **pRgg3-SHP3(G9M)**<br>*7_SHP3_G9M · 1080 bp · linear*<br>Fig. 5a; Suppl. Fig. 12<br>Encoded peptide: **DIIIIVGM**<br><br>**Annotated features**<br>lacZα (fragment) 67–135<br>Rgg3 promoter 251–361<br>Rgg binding site 304–328<br>-10 345–350<br>Transcription start site 358–358<br>pHP14 mRNA stability hairpin 366–406<br>BamHI-BglII Scar 407–412<br>RBS (strong) 413–426<br>Start codon (ATG) 427–429<br>SHP3 Cognate Peptide 430–453<br>G9M 451–453<br>rrnB terminator (TrrnB) 463–830<br>lac operator 928–950 | 1 GAGAGTGCACCATATGCGGTGTGAAATACCGCACAGATGCGTAAGGAGAAAATACCGCATCAGGCGCCATTCGCCATTCA<br>81 GGCTGCGCAACTGTTGGGAAGGGCGATCGGTGCGGGCCTCTTCGCTATTACGCCAGCTGGCGAAAGGGGGATGTGCTGCA<br>161 AGGCGATTAAGTTGGGTAACGCCAGGGTTTTCCCAGTCACGACGTTGTAAAACGACGGCCAGTAAGATCTCGGCTGAATT<br>241 CTAAAGATCTTATGGCTTTCCTTTTCGAACTTTTCTAAGTGTGACACCAAGTTTTGATTTCATATTTTCCCACTTTCCCA<br>321 ACAAAAATCAATTCCGTTTTTTTTTATTGTACAATAAAACGGGCTACGTCGACTCTCGAGTGAGATTGTTGACGGTACCG<br>401 TATTTTGGATCTAGGAGGAAGGATCTATGGATATCATTATTATCGTGGGTATGTAAGGATCTGAAGCTTGGGCCCGAACA<br>481 AAAACTCATCTCAGAAGAGGATCTGAATAGCGCCGTCGACCATCATCATCATCATCATTGAGTTTAAACGGTCTCCAGCT<br>561 TGGCTGTTTTGGCGGATGAGAGAAGATTTTCAGCCTGATACAGATTAAATCAGAACGCAGAAGCGGTCTGATAAAACAGA<br>641 ATTTGCCTGGCGGCAGTAGCGCGGTGGTCCCACCTGACCCCATGCCGAACTCAGAAGTGAAACGCCGTAGCGCCGATGGT<br>721 AGTGTGGGGTCTCCCCATGCGAGAGTAGGGAACTGCCAGGCATCAAATAAAACGAAAGGCTCAGTCGAAAGACTGGGCCT<br>801 TTCGTTTTATCTGTTGTTTGTCGGTGAACTCTGAAAGGAGGAACTATATCCGGATCGAGATCCTCTAGAGTCGACCTGCA<br>881 GGCATGCAAGCTTGGCGTAATCATGGTCATAGCTGTTTCCTGTGTGAAATTGTTATCCGCTCACAATTCCACACAACATA<br>961 CGAGCCGGAAGCATAAAGTGTAAAGCCTGGGGTGCCTAATGAGTGAGCTAACTCACATTAATTGCGTTGCGCTCACTGCC<br>1041 CGCTTTCCAGTCGGGAAACCTGTCGTGCCAGCTGCATTAA |
| **G. mScarlet–SHP3 fusion expression templates** | | | |
| 34 | LET | **pT7-SHP3-mScarlet (N-terminal tag)**<br>*T7_mScarletSHP3_N · 1325 bp · linear*<br>Fig. 3a–d<br>Encoded peptide: **DIIIIVGG**<br><br>**Annotated features**<br>lacZα (fragment) 67–135<br>T7 promoter 251–269<br>RBS 303–325<br>SHP3 Cognate Peptide 335–358<br>mScarlet-I 335–358<br>lac operator 1173–1195 | 1 GAGAGTGCACCATATGCGGTGTGAAATACCGCACAGATGCGTAAGGAGAAAATACCGCATCAGGCGCCATTCGCCATTCA<br>81 GGCTGCGCAACTGTTGGGAAGGGCGATCGGTGCGGGCCTCTTCGCTATTACGCCAGCTGGCGAAAGGGGGATGTGCTGCA<br>161 AGGCGATTAAGTTGGGTAACGCCAGGGTTTTCCCAGTCACGACGTTGTAAAACGACGGCCAGTAAGATCTCGGCTGAATT<br>241 CTAAAGATCTTAATACGACTCACTATAGGGAGACCACAACGGTTTCCCCCTCTAGAAATAATTTTGTTTAACTTTAAGAA<br>321 GGAGATATACCATGGATATCATTATTATCGTGGGTGGTGTGAGCAAGGGCGAGGCAGTGATCAAGGAGTTCATGCGGTTC<br>401 AAGGTGCACATGGAGGGCTCCATGAACGGCCACGAGTTCGAGATCGAGGGCGAGGGCGAGGGCCGCCCCTACGAGGGCAC<br>481 CCAGACCGCCAAGCTGAAGGTGACCAAGGGTGGCCCCCTGCCCTTCTCCTGGGACATCCTGTCCCCTCAGTTCATGTACG<br>561 GCTCCAGGGCCTTCATCAAGCACCCCGCCGACATCCCCGACTACTATAAGCAGTCCTTCCCCGAGGGCTTCAAGTGGGAG<br>641 CGCGTGATGAACTTCGAGGACGGCGGCGCCGTGACCGTGACCCAGGACACCTCCCTGGAGGACGGCACCCTGATCTACAA<br>721 GGTGAAGCTCCGCGGCACCAACTTCCCTCCTGACGGCCCCGTAATGCAGAAGAAGACAATGGGCTGGGAAGCGTCCACCG<br>801 AGCGGTTGTACCCCGAGGACGGCGTGCTGAAGGGCGACATTAAGATGGCCCTGCGCCTGAAGGACGGCGGCCGCTACCTG<br>881 GCGGACTTCAAGACCACCTACAAGGCCAAGAAGCCCGTGCAGATGCCCGGCGCCTACAACGTCGACCGCAAGTTGGACAT<br>961 CACCTCCCACAACGAGGACTACACCGTGGTGGAACAGTACGAACGCTCCGAGGGCCGCCACTCCACCGGCGGCATGGACG<br>1041 AGCTGTACAAGTAATAAGCTGAAAGCCAATTCTGACTGAAAGGAGGAACTATATCCGGATCGAGATCCTCTAGAGTCGAC<br>1121 CTGCAGGCATGCAAGCTTGGCGTAATCATGGTCATAGCTGTTTCCTGTGTGAAATTGTTATCCGCTCACAATTCCACACA<br>1201 ACATACGAGCCGGAAGCATAAAGTGTAAAGCCTGGGGTGCCTAATGAGTGAGCTAACTCACATTAATTGCGTTGCGCTCA<br>1281 CTGCCCGCTTTCCAGTCGGGAAACCTGTCGTGCCAGCTGCATTAA |

| # | Type | Name and annotated features | Sequence (5′→3′) |
|---|---|---|---|
| 35 | LET | **pT7-mScarlet-SHP3 (C-terminal tag)**<br>*T7_mScarSHP3_C · 1325 bp · linear*<br>Fig. 3a–d<br>Encoded peptide: **DIIIIVGG**<br><br>**Annotated features**<br>lacZα (fragment) 67–135<br>T7 promoter 251–269<br>RBS 303–325<br>mScarlet-I 335–1054<br>SHP3 Cognate Peptide 1028–1051<br>lac operator 1173–1195 | 1 GAGAGTGCACCATATGCGGTGTGAAATACCGCACAGATGCGTAAGGAGAAAATACCGCATCAGGCGCCATTCGCCATTCA<br>81 GGCTGCGCAACTGTTGGGAAGGGCGATCGGTGCGGGCCTCTTCGCTATTACGCCAGCTGGCGAAAGGGGGATGTGCTGCA<br>161 AGGCGATTAAGTTGGGTAACGCCAGGGTTTTCCCAGTCACGACGTTGTAAAACGACGGCCAGTAAGATCTCGGCTGAATT<br>241 CTAAAGATCTTAATACGACTCACTATAGGGAGACCACAACGGTTTCCCCCTCTAGAAATAATTTTGTTTAACTTTAAGAA<br>321 GGAGATATACCATGGTGAGCAAGGGCGAGGCAGTGATCAAGGAGTTCATGCGGTTCAAGGTGCACATGGAGGGCTCCATG<br>401 AACGGCCACGAGTTCGAGATCGAGGGCGAGGGCGAGGGCCGCCCCTACGAGGGCACCCAGACCGCCAAGCTGAAGGTGAC<br>481 CAAGGGTGGCCCCCTGCCCTTCTCCTGGGACATCCTGTCCCCTCAGTTCATGTACGGCTCCAGGGCCTTCATCAAGCACC<br>561 CCGCCGACATCCCCGACTACTATAAGCAGTCCTTCCCCGAGGGCTTCAAGTGGGAGCGCGTGATGAACTTCGAGGACGGC<br>641 GGCGCCGTGACCGTGACCCAGGACACCTCCCTGGAGGACGGCACCCTGATCTACAAGGTGAAGCTCCGCGGCACCAACTT<br>721 CCCTCCTGACGGCCCCGTAATGCAGAAGAAGACAATGGGCTGGGAAGCGTCCACCGAGCGGTTGTACCCCGAGGACGGCG<br>801 TGCTGAAGGGCGACATTAAGATGGCCCTGCGCCTGAAGGACGGCGGCCGCTACCTGGCGGACTTCAAGACCACCTACAAG<br>881 GCCAAGAAGCCCGTGCAGATGCCCGGCGCCTACAACGTCGACCGCAAGTTGGACATCACCTCCCACAACGAGGACTACAC<br>961 CGTGGTGGAACAGTACGAACGCTCCGAGGGCCGCCACTCCACCGGCGGCATGGACGAGCTGTACAAGGATATCATTATTA<br>1041 TCGTGGGTGGTTAATAAGCTGAAAGCCAATTCTGACTGAAAGGAGGAACTATATCCGGATCGAGATCCTCTAGAGTCGAC<br>1121 CTGCAGGCATGCAAGCTTGGCGTAATCATGGTCATAGCTGTTTCCTGTGTGAAATTGTTATCCGCTCACAATTCCACACA<br>1201 ACATACGAGCCGGAAGCATAAAGTGTAAAGCCTGGGGTGCCTAATGAGTGAGCTAACTCACATTAATTGCGTTGCGCTCA<br>1281 CTGCCCGCTTTCCAGTCGGGAAACCTGTCGTGCCAGCTGCATTAA |

#### Sequence Table 2. Primers used for LET generation for study DNA templates

| Primer Type | Forward Sequence (5'-3') | Reverse Sequence (5'-3') |
|---|---|---|
| **RRNPP-aTF Primers**<br>for use on generating aTFs from pT7-TF plasmids | cgaactgagatacctacagcg | tcgtcactcatggtgatttctc |
| **pRRNPP-sfGFP Reporter Primers**<br>for use generating reporter LETs from pRRNPP-sfGFP Plasmids | AGAAAATACCGCATCAGGCG | GGTTTCCCGACTGGAAAG |
| **LET Primers**<br>for use amplifying all LET sequences off gBlock starting material | gagagtgcaccatatgcg | ttaatgcagctggcacg |

**Sequence Table 3. Rgg3 plasmids used for cell reactions in the study.** pREG denotes regulator plasmid for expressing transcription factor and peptide. pREP denotes the reporter plasmid containing the GFP reporter under control of the transcription factor-regulated promoter. For constructing the additional plasmids used in this study, the Rgg3 transcription factor sequence and the SHP3 peptide sequence were swapped for the desired DNA part found in Sequence Table 4.

| Type | Name | Sequence (5′ to 3′) |
|---|---|---|
| Plasmid | **pREG+Rgg3+SHP3**<br>colE1<br>ECK12_lpd term<br>P150 promoter<br>riboJ<br>BCD4.6k<br>RBS<br>Rgg3<br>AmpR promoter<br>AmpR CDS<br>Term BBa_B0011<br>AraC<br>araBad promoter<br>pHP14 mRNA stability hairpin<br>RBS<br>SHP3 cognate peptide | ACATGAATGGTCATCGGTTTCCGTGTTTCGTAAAGTCTGGAAACGCGGAAGTCAGCGCCCTGCACCATTATGTTCCGGATCTGCATCGCAGGATGCTGCTGGCTAC<br>CCTGTGGAACACCTACATCTGTATTAACGAAGCGCTGGCATTGACCCTGAGTGATTTTTCTCTGGTCCCGCCGCATCCATACCGCCAGTTGTTTACCCTCACAACGT<br>TCCAGTAACCGGGCATGTTCATCATCAGTAACCCGTATCGTGAGCATCCTCTCTCGTTTCATCGGTATCATTACCCCCATGAACAGAAATCCCCCTTACACGGAGGC<br>ATCAGTGACCAAACAGGAAAAAACCGCCCTTAACATGGCCCGCTTTATCAGAAGCCAGACATTAACGCTTCTGGAGAAACTCAACGAGCTGGATGCGGATGAACAG<br>GCAGACATCTGTGAATCGCTTCACGACCACGCTGATGAGCTTTACCGCAGCTGCCTCGCGCGTTTCGGTGATGACGGTGAAAACCTCTGACACATGCAGCTCCCG<br>CAGACGGTCACAGCTTGTCTGTAAGCGGATGCCGGGAGCAGACAAGCCCGTCAGGGCGCGTCAGCGGGTGTTGGCGGGTGTCGGGGCGCAGCCATGACCCAGT<br>CACGTAGCGATAGCGGAGTGTATACTGGCTTAACTATGCGGCATCAGAGCAGATTGTACTGAGAGTGCACCGGTGTGAAATACCGCACAGATGCGTAAGGAGAAA<br>ATACCGCATCAGGCGCTCTTCCGCTTCCTCGCTCACTGACTCGCTGCGCTCGGTCGTTCGGCTGCGGCGAGCGGTATCAGCTCACTCAAAGGCGGTAATACGGTT<br>ATCCACAGAATCAGGGGATAACGCAGGAAAGAACATGTGAGCAAAAGGCCAGCAAAAGGCCAGGAACCGTAAAAAGGCCGCGTTGCTGGCGTTTTTCCATAGGCT<br>CCGCCCCCCTGACGAGCATCACAAAAATTGACGCTCAAGTCAGAGGTGGCGAAACCCGACAGGACTATAAAGATACCAGGCGTTTCCCCCTGGAAGCTCCCTCGT<br>GCGCTCTCCTGTTCCGACCCTGCCGCTTACCGGATACCTGTCCGCCTTTCTCCCTTCGGGAAGCGTGGCGCTTTCTCATAGCTCACGCTGTAGGTATCTCAGTTCG<br>GTGTAGGTCGTTCGCTCCAAGCTGGGCTGTGTGCACGAACCCCCCGTTCAGCCCGACCGCTGCGCCTTATCCGGTAACTATCGTCTTGAGTCCAACCCGGTAAGA<br>CACGACTTATCGCCACTGGCAGCAGCCACTGGTAACAGGATTAGCAGAGCGAGGTATGTAGGCGGTGCTACAGAGTTCTTGAAGTGGTGGCCTAACTACGGCTAC<br>ACTAGAAGGACAGTATTTGGTATCTGCGCTCTGCTGAAGCCAGTTACCTTCGGAAAAAGAGTTGGTAGCTCTTGATCCGGCAAACAAACCACCGCTGGTAGCGGTG<br>GTTTTTTTGTTTGCAAGCAGCAGATTACGCGCAGAAAAAAGGATCTCAAGAAGATCCTTTGATCTTTTCTACGGGGTCTGACGCTCAGTGGAACGAAAACTCACGTT<br>AAGGCCCTCTCCAAGACCGAGCCATCAACAAAGCGTCTCGCTGAGGTTTCATGGAGCCTCTGGTTCATCTCCGGCAATTAAAAAAGCGGCTAACCACGCCGCTTTT<br>TTTACGTCTGCAGGAACGGGCTGTCGACCTTTGAAAGTTCGTTTACCGCTAGCTCAGTCCTAGGTACAATTACAGCCATCGTAGAAGACACTGGCAGCTGTCACC<br>GGATGTGCTTTCCGGTCTGATGAGTCCGTGAGGACGAAACAGCCTCTACAAATAATTTTGTTTAAGGGCCCAAGTTCACTTAAAAAGGAGATCAACAATGAAAGCAA<br>TTTTCGTACTGAAACATCTTAATCATGCGCCGGTGTTTTAAATATGAAGTCTAAACTGGGGTCGACGCTTCGCAAAGTGAGAAACGGCAAGCAGATCTCTATCTGTTC<br>CGTTGCAGACGAGCACCTCTCTAAGTCCCAAATATCGAGATTTGAACGTGGAGAGAGCGAAATTAGCTGCATTCGGTTAATCAACATACTTGACAAACTTCATATAA<br>CGCTTGACGAATTCCTCATATTGCACGACGAGGATTACACAAAGACGGAAAGTTTTGCCAACCTGGTGCAATACATCCGTAAGCAGTACTCACTGCAGAACATAAAC<br>AATATTCAATCATTATTATCCGATTCGTCTAACTACACATTAGACCCGTTCGAAAAGACAATGGTGAAGTCCATCCTGCATACCATGGACTCGTCCATTATTCCATCAG<br>ACGACGAGTTGCTCCAGTTGGCAGATTATCTGTTCAAGGTGGAGAAGTGGGGCTATTATGAAATTATTCTGTTGGGCAACTGCGTTCGGACTATTGATTACAACAGC<br>GTGTTCTTGCTGACCAAGGAGATGTTAAACAACTATATTTATTCCTCTCTGAACAAAACCAACAAGCGGATAGTTACACAGCTGGCTATCAACTGCTTAATTTTGAGC<br>ATAGATATGGAAGAGTTCACTAACTGTTTCTACCTGATAGACGAAATCAAAGCCTTACTTGACAACGAGCTTAACTTCTACGAGCAAACGGTATTCCTCTATGCCACC |

| Type | Name | Sequence (5′ to 3′) |
| --- | --- | --- |
| | Term BBa_B1002 | GGTTACTTTGAGTTCAAGCGGTGGCAATCGACTTCGGGCATTGAGAAGATGAAGCAAGCAATCCAAGTGTTAGACATATTGGGCGAAGATAACCTTAAGCTGCACT<br>ACACAATTCACTTTGACAAACTTATTAATAACAAGTGATAATCCTAGTCTTCTCGCCACTTTCAGCCAAAAAACTTAAGACCGCCGGTCTTGTCCACTACCTTGCAGTA<br>ATGCGGTGGACAGGATCGGCGGTTTTCTTTTCTCTTCTCAACACCCTTCGCGTCAACACTTTTCCGCCAAGGAGACGGTTGGTCAGGTTTTCGGGAGGTGTGGCTG<br>GAAGTTCCTATACTTTCTAGAGAATAGGAACTTCTTTCTAAATACATTCAAATATGTATCCGCTCATGAGACAATAACCCTGATAAATGCTTCAATAATATTGAAAAAG<br>GAAGAGTATGAGTATTCAACATTTCCGTGTCGCCCTTATTCCCTTTTTTGCGGCATTTTGCCTTCCTGTTTTTGCTCACCCAGAAACGCTGGTGAAAGTAAAAGATGC<br>TGAAGATCAGTTGGGTGCACGAGTGGGTTACATCGAACTGGATCTCAACAGCGGTAAGATCCTTGAGAGTTTTCGCCCCGAAGAACGTTTTCCAATGATGAGCACT<br>TTTAAAGTTCTGCTATGTGGCGCGGTATTATCCCGTGTTGACGCCGGGCAAGAGCAACTCGGTCGCCGCATACACTATTCTCAGAATGACTTGGTTGAGTACTCAC<br>CAGTCACAGAAAAGCATCTTACGGATGGCATGACAGTAAGAGAATTATGCAGTGCTGCCATAACCATGAGTGATAACACTGCGGCCAACTTACTTCTGACAACGATC<br>GGAGGACCGAAGGAGCTAACCGCTTTTTTGCACAACATGGGGGATCATGTAACTCGCCTTGATCGTTGGGAACCGGAGCTGAATGAAGCCATACCAAACGACGAG<br>CGTGACACCACGATGCCTGCAGCAATGGCAACAACGTTGCGCAAACTATTAACTGGCGAACTACTTACTCTAGCTTCCCGGCAACAATTAATAGACTGGATGGAGG<br>CGGATAAAGTTGCAGGACCACTTCTGCGCTCGGCCCTTCCGGCTGGCTGGTTTATTGCTGATAAATCTGGAGCCGGTGAGCGTGGATCGCGCGGTATCATTGCAG<br>CACTGGGGCCAGATGGTAAGCCCTCCCGTATCGTAGTTATCTACACGACGGGGAGTCAGGCAACTATGGATGAACGAAATAGACAGATCGCTGAGATAGGTGCCT<br>CACTGATTAAGCATTGGTAAGTTGTGATGGCGGTAGGAATGTAATCGTTAATCCGCAAATAACGTAAAAACCCGCTTCGGCGGGTTTTTTTATGGGGGGAGTTTAGG<br>GAAAGAGCATTTGTCATCCCGTTGAATATGGCTCCCTTAACGTGAGGAAGTTCCTATACTTTCTAGAGAATAGGAACTTCTACAGATGGACTTGGGTTGGCGGCCTA<br>GACGTGTGATAGACTCAGTCGGCTTCGATGTTTGAGTTGTGTGTTTTTACATCATATGACATCAGGAGTCTGCAAAACGTCTGCGACCTGAGCAACAAGAGAATATA<br>AAAAGCCAGATTATTAATCCGGCTTTTTTATTATTTGGAAAATCAGTTATGACAACTTGACGGCTACATCATTCACTTTTTCTTCACAACCGGCACGGAACTCGCTCGG<br>GCTGGCCCCGGTGCATTTTTTAAATACCCGCGAGAAATAGAGTTGATCGTCAAAACCAACATTGCGACCGACGGTGGCGATAGGCATCCGGGTGGTGCTCAAAAG<br>CAGCTTCGCCTGGCTGATACGTTGGTCCTCGCGCCAGCTTAAGACGCTAATCCCTAACTGCTGGCGGAAAAGATGTGACAGACGCGACGGCGACAAGCAAACATG<br>CTGTGCGACGCTGGCGATATCAAAATTGCTGTCTGCCAGGTGATCGCTGATGTACTGACAAGCCTCGCGTACCCGATTATCCATCGGTGGATGGAGCGACTCGTTA<br>ATCGCTTCCATGCGCCGCAGTAACAATTGCTCAAGCAGATTTATCGCCAGCAGCTCCGAATAGCGCCCTTCCCCTTGCCCGGCGTTAATGATTTGCCCAAACAGGT<br>CGCTGAAATGCGGCTGGTGCGCTTCATCCGGGCGAAAGAACCCCGTATTGGCAAATATTGACGGCCAGTTAAGCCATTCATGCCAGTAGGCGCGCGGACGAAAGT<br>AAACCCACTGGTGATACCATTCGCGAGCCTCCGGATGACGACCGTAGTGATGAATCTCTCCTGGCGGGAACAGCAAAATATCACCCGGTCGGCAAACAAATTCTCG<br>TCCCTGATTTTTCACCACCCCCTGACCGCGAATGGTGAGATTGAGAATATAACCTTTCATTCCCAGCGGTCGGTCGATAAAAAAATCGAGATAACCGTTGGCCTCAA<br>TCGGCGTTAAACCCGCCACCAGATGGGCATTAAACGAGTATCCCGGCAGCAGGGGATCATTTTGCGCTTCAGCCATACTTTTCATACTCCCGCCATTCAGAGAAGA<br>AACCAATTGTCCATATTGCATCAGACATTGCCGTCACTGCGTCTTTTACTGGCTCTTCTCGCTAACCAAACCGGTAACCCCGCTTATTAAAAGCATTCTGTAACAAAG<br>CGGGACCAAAGCCATGACAAAAACGCGTAACAAAAGTGTCTATAATCACGGCAGAAAAGTCCACATTGATTATTTGCACGGCGTCACACTTTGCTATGCCATAGCAT<br>TTTTATCCATAAGATTAGCGGATCCTACCTGACGCTTTTTATCGCAACTCTCTACTGTTTCTCCATGGCTACGTCGACTCTCGAGTGAGATTGTTGACGGTACCGTAT<br>TTTGGATCTAGGAGGAAGGATCTATGGATATCATTATTATCGTGGGTGGTTAACGCAAAAAACCCCGCTTCGGCGGGGTTTTTTCGC |

| Type | Name | Sequence (5′ to 3′) |
| --- | --- | --- |
| Plasmid | **pREP_Rgg3**<br>p15a<br>CAT<br>LuxI terminator<br>EZeo CDS<br>RBS<br>Lambda promoter<br>Bidirectional terminator<br>Rgg3 promoter<br>RBS<br>sfGFP<br>rrnb.TE Tandem Terminator<br>LacI | TTTGAAGAGATAAATTGCACTGAAATCTAGAAATATTTTATCTGATTAATAAGATGATCTTCTTGAGATCGTTTTGGTCTGCGCGTAATCTCTTGCTCTGAAAACGAAA<br>AAACCGCCTTGCAGGGCGGTTTTTCGAAGGTTCTCTGAGCTACCAACTCTTTGAACCGAGGTAACTGGCTTGGAGGAGCGCAGTCACCAAAACTTGTCCTTTCAGT<br>TTAGCCTTAACCGGCGCATGACTTCAAGACTAACTCCTCTAAATCAATTACCAGTGGCTGCTGCCAGTGGTGCTTTTGCATGTCTTTCCGGGTTGGACTCAAGACGA<br>TAGTTACCGGATAAGGCGCAGCGGTCGGACTGAACGGGGGGTTCGTGCATACAGTCCAGCTTGGAGCGAACTGCCTACCCGGAACTGAGTGTTAGGCGTGGAAT<br>GAGACAAACGCGGCCATAACAGCGGAATGACACCGGTAAACCGAAAGGCAGGAACAGGAGAGCGCACGAGGGAGCCACCAGGGGGAAACGCCTGGTATCTTTAT<br>AGTCCTGTCGGGTTTCGCCACCACTGATTTGAGCGTCAGATTTCGTGATGCTTGTCAGGGGGGCGGAGCCTATGGAAAAACGGCTTTGCCGCGGCCCTCTCACTT<br>CCCTGTTAAGTATCTTCCTGGCATCTTCCAGGAAATCTCCGCCCCGTTCGTAAGCCATTTCCGCTCGCCGCAGTCGAACGACCGAGCGTAGCGAGTCAGTGAGCG<br>AGGAAGCGGAATATATCCTGTATCACATATTCTGCTGACGCACCGGTGCAGCCTTTTTTCTCCTGCCACATGAAGCACTTCACTGACACCCTCATCAGTGCCAACAT<br>AGTAAGCCAGTATACACTCCGCTAGCGCTGATGTCCGGCGGTGCTTTTGCCGTTACGCACCACCCCGTCAGTAGCTGAACAGGAGGGACAGCTGATAGAAACAGA<br>AGCCACTGGAGCACCTCAAAAACACCATCATACACTAAATCAGTAAGTTGGCAGCATCACCCGACGCACTTTGCGCCGAATAAATACCTGTGACGGAAGATCACTT<br>CGCAGAATAAATAAATCCTGGTGTCCCTGTTGATACCGGGAAGCCCTGGGCCAACTTTTGGCGAAAATGAGACGTTGATCGGCACGTAAGAGGTTCCAACTTTCAC<br>CATAATGAAATAAGATCACTACCGGGCGTATTTTTTGAGTTATCGAGATTTTCAGGAGCTAAGGAAGCTAAAATGGAGAAAAAAATCACTGGATATACCACCGTTGAT<br>ATATCCCGATGGCATCGTAAAGAACATTTTGAGGCATTTCAGTCAGTTGCTCAATGTACCTATAACCAGACCGTTCAGCTGGATATTACGGCCTTTTTAAAGACCGTA<br>AAGAAAAATAAGCACAAGTTTTATCCGGCCTTTATTCACATTCTTGCCCGCCTGATGAATGCTCATCCGGAGTTCCGTATGGCAATGAAAGACGGTGAGCTGGTGAT<br>ATGGGATAGTGTTCACCCTTGTTACACCGTTTTCCATGAGCAAACTGAAACGTTTTCATCGCTCTGGAGTGAATACCACGACGATTTCCGGCAGTTTCTACACATATA<br>TTCGCAAGATGTGGCGTGTTACGGTGAAAACCTGGCCTATTTCCCTAAAGGGTTTATTGAGAATATGTTTTTCGTCTCAGCCAATCCCTGGGTGAGTTTCACCAGTTT<br>TGATTTAAACGTGGCCAATATGGACAACTTCTTCGCCCCCGTTTTCACTATGGGCAAATATTATACGCAAGGCGACAAGGTGCTGATGCCGCTGGCGATTCAGGTTC<br>ATCATGCCGTTTGTGATGGCTTCCATGTCGGCAGAATGCTTAATGAATTACAACAGTACTGCGATGAGTGGCAGGGCGGGGCGTAATTTTTTTAAGGCAGTTATTGG<br>TGCCCTTAAACGCCTGGTTGCTACGCCTGAATAAGTGATAATAAGCGGATGAATGGCAGAAATTCGAAAGCAAATTCGACCCGGTCGTCGGTTCAGGGCAGGGTCG<br>TTAAATAGCCGCTTATGTCTATTGCTGGTTTACCGGTTTATTGACTACCGGAAGCAGTGTGACCGTGTGCTTCTCAAATGCCTGAGGTTTCAGAAATAATAAAAAAGC<br>CGGATTAATAATCTGGCTTTTTATATTCTCTCTTTATTAATCTTGTTCTTCAGCAACAAAATGAACGCAATTACCAGCTGGATCTCTCAAAGCAAATTCTCTACCCCAAG<br>GTTGTTCACCAATTTCGGTCATAGCTGGACCAGAAGCATCTCTAAAATTGGTAGAAACAACTTCGGACCATTCGGCATATAATTCATCCAAACCTCTAACCCAAACCC<br>AAGCCAAAGTATTATCTGGAACAACTTGATCTTGAACGGCGGAAATGAACAAGGTAACATCATCTCTAACAACACCAGCGAAATCATCTTCAACGAAATCTCTGGAG<br>AAACCCAATCTATCAGTCCAAAATTCAACAGCACCAGCAACATCTCTAGCGGTCAAAACTGGAACAGCAGATGTCAATTTAGACATATGTCAACCTCCTCCAATAACC<br>GTCACTGCTGACGACTCTACCATTATCACTTGCAACCATTATCACCGCCAGAGGTAAAATAGTCAACACGCACGGTGTTAGTCTTTTCGCCGGTGTATTTCTTGCAAA<br>AAAAAACCCCGCCGAAGCGGGGTTTTTTTTTACAGGGCTATAAGGTAGATCCGGGAGCTGTATGGCTTTCCTTTTCGAACTTTTCTAAGTGTGACACCAAGTTTTGA<br>TTTCATATTTTCCCACTTTCCCAACAAAAATCAATTCCGTTTTTTTTTATTGTACAATAAAACGGGGTTTGATTTAATAAGGAGGTATACATATGGCATCCAAGGGCGA<br>GGAGCTCTTTACTGGCGTAGTACCAATTCTCGTAGAGCTCGATGGCGATGTAAATGGCCATAAGTTTTCCGTACGCGGCGAGGGCGAGGGCGATGCAACTAACGG<br>CAAGCTCACTCTCAAGTTTATTTGTACTACTGGCAAGCTCCCAGTACCATGGCCAACTCTCGTAACTACTCTGACCTATGGCGTACAATGTTTTTCCCGCTATCCAGA |

| Type | Name | Sequence (5′ to 3′) |
|---|---|---|
| | | TCACATGAAGCAACATGATTTTTTTAAGTCCGCAATGCCAGAGGGCTATGTACAAGAGCGCACTATTAGCTTTAAGGATGATGGCACCTATAAGACTCGCGCAGAGGTAAAGTTTGAGGGCGATACTCTCGTAAATCGCATTGAGCTCAAGGGCATTGATTTTAAGGAGGATGGCAATATTCTCGGCCATAAGCTGGAGTATAATTTCAATTCCCATAATGTATATATTACCGCAGATAAGCAAAAGAATGGCATTAAGGCGAATTTTAAGATTCGCCATAATGTGGAGGATGGCTCCGTACAACTCGCAGATCATTATCAACAAAATACTCCAATTGGCGATGGCCCAGTACTCCTCCCAGATAATCATTATCTCTCCACTCAATCCGTGCTCTCCAAAGATCCAAATGAGAAGCGCGATCACATGGTACTCCTGGAGTTTGTAACTGCAGCAGGCATTACTCATGGCATGGATGAGCTCTATAAGCTCGAGCACCACCACCACCACCACTAATAACCAGGCATCAAATAAAACGAAAGGCTCAGTCGAAAGACTGGGCCTTTCGTTTTATCTGTTGTTTGTCGGTGAACGCTCTCTACTAGAGTCACACTGGCTCACCTTCGGGTGGGCCTTTCTGCGTTTATAATTTCCTAATGCAGGAGTCGCATAAGGGAGAGCGTCGAGATCCCGGACACCATCGAATGGCGCAAAACCTTTCGCGGTATGGCATGATAGCGCCCGGAAGAGAGTCAATTCAGGGTGGTGAATGTGAAACCAGTAACGTTATACGATGTCGCAGAGTATGCCGGTGTCTCTTATCAGACCGTTTCCCGCGTGGTGAACCAGGCCAGCCACGTTTCTGCGAAAACGCGGGAAAAAGCGGAAGCGGCGATGGCGGAGCTGAATTACATTCCCAACCGCGTGGCACAACAACTGGCGGGCAAACAGTCGTTGCTGATTGGCGTTGCCACCTCCAGTCTGGCCCTGCACGCGCCGTCGCAAATTGTCGCGGCGATTAAATCTCGCGCCGATCAACTGGGTGCCAGCGTGGTGGTGTCGATGGTAGAACGAAGCGGCGTCGAAGCCTGTAAAGCGGCGGTGCACAATCTTCTCGCGCAACGCGTCAGTGGGCTGATCATTAACTATCCGCTGGATGACCAGGATGCCATTGCTGTGGAAGCTGCCTGCACTAATGTTCCGGCGTTATTTCTTGATGTCTCTGACCAGACACCCATCAACAGTATTATTTTCTCCCATGAAGACGGTACGCGACTGGGCGTGGAGCATCTGGTCGCATTGGGTCACCAGCAAATCGCGCTGTTAGCGGGCCTATTAAGTTCTGTCTCGGCGCGTCTGCGTCTGGCTGGCTGGCATAAATATCTCACTCGCAATCAAATTCAGCCGATAGCGGAACGGGAAGGCGACTGGAGTGCCATGTCCGGTTTTCAACAAACCATGCAAATGCTGAATGAGGGCATCGTTCCCACTGCGATGCTGGTTGCCAACGATCAGATGGCGCTGGGCGCAATGCGCGCCATTACCGAGTCCGGGCTGCGCGTTGGTGCGGACATCTCGGTAGTGGGATACGACGATACCGAAGACAGCTCATGTTATATCCCGCCGTTAACCACCATCAAACAGGATTTTCGCCTGCTGGGGCAAACCAGCGTGGACCACTTGCTGCAACTCTCTCAGGGCCAGGCGGTGAAGGGCAATCAGCTGTTGCCCGTCTCACTGGTGAAAAGAAAAACCACCCTGGCGCCCAATACGCAAACCGCCTCTCCCCGCGCGTTGGCCGATTCATTAATGCAGCTGGCACGACAGGTTTCCCGACTGGAAAGCGGGCAGTGAGCGCAACGCAATTAATGTAAGTTAGCTCACTCATTAGGCACCGGGATCTCGACCGATGCCCTTGAGAGCCTTCAACCCAGTCAGCTCCTTCCGGTGGGCGCGGGGCATGACTAACATGAGAATTACAACTTATATCGTATGGGGCTGACTTCAGGTGCTACA |

#### Sequence Table 4. DNA sequences used for cloning plasmids for cell reactions in this study.

| Type | Name | Sequence (5′ to 3′) |
|---|---|---|
| Transcription factor | **Rgg3** | ATGAAGTCTAAACTGGGGTCGACGCTTCGCAAAGTGAGAAACGGCAAGCAGATCTCTATCTGTTCCGTTGCAGACGAGCACCTCTCTAAGTCCCAAATATCGAGATTTGAACGTGGAGAGAGCGAAATTAGCTGCATTCGGTTAATCAACATACTTGACAAACTTCATATAACGCTTGACGAATTCCTCATATTGCACGACGAGGATTACACAAAGACGGAAAGTTTTGCCAACCTGGTGCAATACATCCGTAAGCAGTACTCACTGCAGAACATAAACAATATTCAATCATTATTATCCGATTCGTCTAACTACACATTAGACCCGTTCGAAAAGACAATGGTGAAGTCCATCCTGCATACCATGGACTCGTCCATTATTCCATCAGACGACGAGTTGCTCCAGTTGGCAGATTATCTGTTCAAGGTGGAGAAGTGGGGCTATTATGAAATTATTCTGTTGGGC |

| Type | Name | Sequence (5′ to 3′) |
| --- | --- | --- |
| | | AACTGCGTTCGGACTATTGATTACAACAGCGTGTTCTTGCTGACCAAGGAGATGTTAAACAACTATATTTATTCCTCTCTGAACAAAACCAACAAGCGGATAGTTACACAGCTGGCTATCAACTGCTTAATTTTGAGCATAGATATGGAAGAGTTCACTAACTGTTTCTACCTGATAGACGAAATCAAAGCCTTACTTGACAACGAGCTTAACTTCTACGAGCAAACGGTATTCCTCTATGCCACCGGTTACTTTGAGTTCAAGCGGTGGCAATCGACTTCGGGCATTGAGAAGATGAAGCAAGCAATCCAAGTGTTAGACATATTGGGCGAAGATAACCTTAAGCTGCACTACACAATTCACTTTGACAAACTTATTAATAACAAG |
| Transcription factor | **Rgg2** | ATGGACAAAGAGCTTGGTAAGACCCTGCGCCGCCTGCGCCAAGGAAAACAAGTTAGTATATCCAGCCTTGCGGACGAGCATTTATCGAAAAGTCAAATCTCGCGTTTTGAGCGAGGAGAGTCAGAAATTAGTTGTTCTCGGCTTTTAAACTTATTGGACAAACTTAACATCACAATAGACGAGTTTGTATCTACGCACAGCAAAACCCACACGCACTTCTTTACGCTCCTTTCCCGTGTCCGGAAGTACTATGCGGAAAAGAACGTGGCTAAGTTACTTAAGCTTCTGGAAGATTACGCCCACAAGGATTACGAGAGTACTATGATCAAGGCGATTTTGTCTAGTATTGAACCGACAGTAGAGCCAAGCGAAGAGGAAGTCACTCGTTTAACCGACTATCTGTTTTCAGTGGAGCAATGGGGTTATTATGAAATTATCTTGCTTGGCAATTGTTCTCGATTCATTAATTACAACACCCTGTTCTTACTGACAAAGGAGATGGTGACGAGTTTTGCATACAGTGAACAAAACAAGACGAACAAGACGCTGGTTACTCAACTCAGTATAAACTGTCTGATTATATCGATCGACTACAGTTACTTTGATCACTCCCACTACTTGATTGAAAAGATCGAATTCCTGCTGCGTGACGAGCTGAACTTTTACGAAAAGACGGTATTCCTCTACGTGCACGGATACTACAAGTTGAAACAGGGTCAAGTTTCGGGTAAGGACGATATGCGTCAGGCCTTACAAATTTTCAAGTACCTGGGCGAAGATGCGCTGTATTACTCCTATAAAGAGCACTACAGAAAGGAAGTGCTCGGTGACGAAGAGGATGAGGAAGAAGGC |
| Transcription factor | **ComR_ST** | ATGAATCTGAAAGACAGTATAGGGTTACGCATTAAGACTGAGCGGGAACGGCAGCAAATGTCCCGCGAGGTCTTGTGCCTTGACGGGGCTGAGTTAACGGTCCGGCAGCTGATTCGCATAGAGAAAGGGGAGTCCCTGCCGTCACTGGATAGACTGTCGTACATCGCGAAGAGATTGGGCAAGTCGATGACCGAATTATTGGACCAAGACAACATTACAATCCCAGACGAGTACTACGAGATGAAGAATCGCCTGATCAAATTTCCGACTTACCGTAACCCCGATCGGATTAAAAGCAAGTTGACTCTTATAGAGGAAGTTTACGAGAAGTTCTTTGACATCCTTCCGGAAGAGGAGTTACTTACATTAGACATACTGGAGAACATCCTGTCATTCACGAGTTGGGAAGAATCGCCCAAAGTCGAGGAAATATATGAGGACTTATTCGAACAAGTAAAGCGCAAGCGTAAATTCAGCACAAATGATCTGCTTGTCATCGATTACTATTTCTTCCACCTGTATGGCCGTAAGCAGTATGACAAGAAGTTGTTCGAGCGTATCATTAAAAGAGTTTTGAACCAAGAGATATGGACCGACGACGTCTACAATATCGTGTTGTTTAACGATTTAATGGCCATCGCGGCGCTCAAGATATTCCACAATAGTTTTAGCGACTTTCTGACGGTTGTTGACAAGGCGTTGGCCGTGATTGAAAAGTCTCAACTGTATTCCTATAAGCCTTCTGTGTTCGTTTTAAAAGCTAAGTACGAACTGCTGCACAAGGAAAACAAGAAGGAAGCCGCCGAAAACTATGACAAGGCTATAGTATTTGCGTCGGTGCTGGAAGACAGTGTTTTAGAGGAGTCTATTAAAGCGGGGAAGCTTGCAGACGGTTTA |
| Transcription factor | **ComR_SI** | ATGGAAAAGTTTGGAGAGAAAATTCGCCTGATGAGAGAAGAGAAGGGTATTTCTAGAGAAGAATTCTGTGGGGACGAGACTGAACTGTCCGTTCGGCAGCTCGCACGAATCGAATTGAACCAGTCCATCCCAAACTTATCAAAAGCCAGTTTTATAGCGAATCGGCTGGGCGTGAAGCTGGGGACGCTGACAGACGGGGACTCATTGGAGCTTCCTAAGCGTTACAAGGAGCTGAAGTACCTTCTGCTTCGTACGCCAACCTACGGTGATCAAGTGAGACTGGATCGTAAGAACGACTATTTTGATGAAATCGCTGAAGTCTTCTACGATGTAATTCCAGAGGAAGAACGGCTGATAATAGACTGCCTTCAATCTAAGTTCGACGTACATTTCTCCGAGGATGTGAATTTCGGCGAGGGTATCTTAAACGACTACTTCGGGCAAGTAAACCGGAAGAAAGTTTTCACCATAAATGACTTAATCCTTATTGATTTGTATTTTGCATGCTTGAGTTCCGCCAAGAAGTTCGAGGGGATATACAGTCTGGACTTTTACGACGATCTTATGAAGAGATTAGTCAATCAAAAGCACGTTTCTCCCGAGACAGATCTTATCCTGAACAACGTTCTGTTAAATAACATAGACTTAGCATTCCAATTCAAACGAGAGTACTACATTGAACGTGTCATCACCATAAGCGAGAAGATTATGACTGAGATCCACGACTTCCAGCGCCGCCCAATCCTGTCACTGGTCGAATGGAAGTATTACCTTAAGCTCAAGCACGATTTTGCGCTGGCGGAACAAAGCTTCACAAATGCTACTCTTTTCGCACGACTGGTTGGAGACACATACTTGGAGAACAAGCTCAAGGAAGAATGGCAACTGGACATCGACGCGGTTGAG |

| Type | Name | Sequence (5′ to 3′) |
|---|---|---|
| Peptide | **SHP3** | ATGGATATCATTATTATCGTGGGTGGTTAA |
| Peptide | **SHP2** | ATGGATATCCTGATTATCGTGGGTGGT |
| Peptide | **XIP_ST** | ATGCCATACTTCGCCGGCTGTCTGTAA |
| Peptide | **XIP_SI** | ATGCTCCTCACAGCGTGGTGGGGCCTA |
| C-terminal fusion protein | **mScarlet-SHP3**<br>Underline region is SHP3 | ATGGTGAGCAAGGGCGAGGCAGTGATCAAGGAGTTCATGCGGTTCAAGGTGCACATGGAGGGCTCCATGAACGGCCACGAGTTCGAGATCGAGGGCGAGGGCGAGGGCCGCC<br>CCTACGAGGGCACCCAGACCGCCAAGCTGAAGGTGACCAAGGGTGGCCCCCTGCCCTTCTCCTGGGACATCCTGTCCCCTCAGTTCATGTACGGCTCCAGGGCCTTCATCAAGC<br>ACCCCGCCGACATCCCCGACTACTATAAGCAGTCCTTCCCCGAGGGCTTCAAGTGGGAGCGCGTGATGAACTTCGAGGACGGCGGCGCCGTGACCGTGACCCAGGACACCTCC<br>CTGGAGGACGGCACCCTGATCTACAAGGTGAAGCTCCGCGGCACCAACTTCCCTCCTGACGGCCCCGTAATGCAGAAGAAGACAATGGGCTGGGAAGCGTCCACCGAGCGGTT<br>GTACCCCGAGGACGGCGTGCTGAAGGGCGACATTAAGATGGCCCTGCGCCTGAAGGACGGCGGCCGCTACCTGGCGGACTTCAAGACCACCTACAAGGCCAAGAAGCCCGTG<br>CAGATGCCCGGCGCCTACAACGTCGACCGCAAGTTGGACATCACCTCCCACAACGAGGACTACACCGTGGTGGAACAGTACGAACGCTCCGAGGGCCGCCACTCCACCGGCGG<br>CATGGACGAGCTGTACAAGGATATCATTATTATCGTGGGTGGT |
| N-terminal fusion protein | **SHP3-mScarlet**<br>Underline region is SHP3 | ATGGATATCATTATTATCGTGGGTGGTGTGAGCAAGGGCGAGGCAGTGATCAAGGAGTTCATGCGGTTCAAGGTGCACATGGAGGGCTCCATGAACGGCCACGAGTTCGAGATC<br>GAGGGCGAGGGCGAGGGCCGCCCCTACGAGGGCACCCAGACCGCCAAGCTGAAGGTGACCAAGGGTGGCCCCCTGCCCTTCTCCTGGGACATCCTGTCCCCTCAGTTCATGT<br>ACGGCTCCAGGGCCTTCATCAAGCACCCCGCCGACATCCCCGACTACTATAAGCAGTCCTTCCCCGAGGGCTTCAAGTGGGAGCGCGTGATGAACTTCGAGGACGGCGGCGCC<br>GTGACCGTGACCCAGGACACCTCCCTGGAGGACGGCACCCTGATCTACAAGGTGAAGCTCCGCGGCACCAACTTCCCTCCTGACGGCCCCGTAATGCAGAAGAAGACAATGGG<br>CTGGGAAGCGTCCACCGAGCGGTTGTACCCCGAGGACGGCGTGCTGAAGGGCGACATTAAGATGGCCCTGCGCCTGAAGGACGGCGGCCGCTACCTGGCGGACTTCAAGACC<br>ACCTACAAGGCCAAGAAGCCCGTGCAGATGCCCGGCGCCTACAACGTCGACCGCAAGTTGGACATCACCTCCCACAACGAGGACTACACCGTGGTGGAACAGTACGAACGCTC<br>CGAGGGCCGCCACTCCACCGGCGGCATGGACGAGCTGTACAAG |
| Reporter protein | **sfGFP** | ATGGCATCCAAGGGCGAGGAGCTCTTTACTGGCGTAGTACCAATTCTCGTAGAGCTCGATGGCGATGTAAATGGCCATAAGTTTTCCGTACGCGGCGAGGGCGAGGGCGATGCA<br>ACTAACGGCAAGCTCACTCTCAAGTTTATTTGTACTACTGGCAAGCTCCCAGTACCATGGCCAACTCTCGTAACTACTCTGACCTATGGCGTACAATGTTTTTCCCGCTATCCAGAT<br>CACATGAAGCAACATGATTTTTTTAAGTCCGCAATGCCAGAGGGCTATGTACAAGAGCGCACTATTAGCTTTAAGGATGATGGCACCTATAAGACTCGCGCAGAGGTAAAGTTTGA<br>GGGCGATACTCTCGTAAATCGCATTGAGCTCAAGGGCATTGATTTTAAGGAGGATGGCAATATTCTCGGCCATAAGCTGGAGTATAATTTCAATTCCCATAATGTATATATTACCGC<br>AGATAAGCAAAAGAATGGCATTAAGGCGAATTTTAAGATTCGCCATAATGTGGAGGATGGCTCCGTACAACTCGCAGATCATTATCAACAAAATACTCCAATTGGCGATGGCCCAG<br>TACTCCTCCCAGATAATCATTATCTCTCCACTCAATCCGTGCTCTCCAAAGATCCAAATGAGAAGCGCGATCACATGGTACTCCTGGAGTTTGTAACTGCAGCAGGCATTACTCATG<br>GCATGGATGAGCTCTATAAGCTCGAGCACCACCACCACCACCAC |

| Type | Name | Sequence (5′ to 3′) |
|---|---|---|
| Promoter | **pRgg3** | TATGGCTTTCCTTTTCGAACTTTTCTAAGTGTGACACCAAGTTTTGATTTCATATTTTCCCACTTTCCCAACAAAAATCAATTCCGTTTTTTTTTATTGTACAATAAAACG |
| Promoter | **pRgg2** | TGTTTTCCTTGGCGTAGTCGCCTTAAGGTTTTACCGAGTTCTTTTTCCATTTTTCCCACTTTCACAACAAAAAATTCAAATCCGTCAAATCCATTATATAATACAAGTA |
| Promoter | **pComR_ST** | GTGGTTTTGACTCTATCTCGCTTATTTAATTGTTTTAGTCAAGTTATTTATAGTGACATATATGTCTCTATTTTATTTTTCTACATTCTTTTATAATAGGTCTA |
| Promoter | **pComR_SI** | AAGCGGTAGACGGAGAGCAACTCAGATGAGTTGCTTTTTTTGTGCATCAATTTGGACATTTTTGTCATGTTCCTTTTCCCTAATCCTTTCTATAATAAGGGTA |

#### Sequence Table 5. Mutations identified from the less stringent screen and subcloned for dose-response analysis.

| Type | Name | Sequence (5′→3′) |
|---|---|---|
| Mutation | **pREG+Rgg3+SHP3D2R** | CGG |
| Mutation | **pREG+Rgg3+SHP3D2A** | GCC |
| Mutation | **pREG+Rgg3+SHP3I3L** | TTG |
| Mutation | **pREG+Rgg3+SHP3I4T** | TGG |
| Mutation | **pREG+Rgg3+SHP3I5V** | GTC |
| Mutation | **pREG+Rgg3+SHP3I5L** | CTC |
| Mutation | **pREG+Rgg3+SHP3I6L** | TTG |
| Mutation | **pREG+Rgg3+SHP3V7T** | ACG |
| Mutation | **pREG+Rgg3+SHP3G8T** | ACC |
| Mutation | **pREG+Rgg3+SHP3G8H** | CAC |

| Type | Name | Sequence (5′→3′) |
| --- | --- | --- |
| Mutation | **pREG+Rgg3+SHP3G9M** | ATG |
| Mutation | **pREG+Rgg3+SHP3G9I** | ATC |

**Sequence Table 6. Mutations identified from the more stringent screen and subcloned for dose-response analysis.**

| Type | Name | Sequence (5′→3′) |
| --- | --- | --- |
| Mutation | **pREG+Rgg3+SHP3D2L** | TTG |
| Mutation | **pREG+Rgg3+SHP3D2G** | GGG |
| Mutation | **pREG+Rgg3+SHP3I3R** | AGG |
| Mutation | **pREG+Rgg3+SHP3I3S** | TCG |
| Mutation | **pREG+Rgg3+SHP3I4D** | GAC |
| Mutation | **pREG+Rgg3+SHP3I4E** | GAG |
| Mutation | **pREG+Rgg3+SHP3I5F** | TTC |
| Mutation | **pREG+Rgg3+SHP3I6T** | ACC |
| Mutation | **pREG+Rgg3+SHP3I6F** | TTC |
| Mutation | **pREG+Rgg3+SHP3V7M** | ATG |
| Mutation | **pREG+Rgg3+SHP3G8W** | TGG |

| Type | Name | Sequence (5′→3′) |
|---|---|---|
| Mutation | **pREG+Rgg3+SHP3G9N** | AAC |
